# PPM1D is a neuroblastoma oncogene and therapeutic target in childhood neural tumors

**DOI:** 10.1101/2020.09.04.283648

**Authors:** Jelena Milosevic, Susanne Fransson, Miklos Gulyas, Gabriel Gallo-Oller, Thale K Olsen, Diana Treis, Malin Wickström, Lotta HM Elfman, Baldur Sveinbjornsson, Falk Hertwig, Christoph Bartenhagen, Susanne Reinsbach, Margareta Wilhelm, Frida Abel, Niloufar Javanmardi, Subazini Thankaswamy-Kosalai, Nina Eissler, Anna Kock, Yao Shi, Keiji Tanino, Jane Y Hehir-Kwa, Arjen Mensenkamp, Godelieve AM Tytgat, Chandrasekhar Kanduri, Johan Holmberg, David Gisselsson, Jan J Molenaar, Marjolijn Jongmans, Matthias Fischer, Marcel Kool, Kazuyasu Sakaguchi, Ninib Baryawno, Tommy Martinsson, John Inge Johnsen, Per Kogner

**Affiliations:** Childhood Cancer Research Unit, Department of Women’s and Children’s Health, Karolinska Institutet, Stockholm, Sweden; Center for Regenerative Medicine, Massachusetts General Hospital, Boston, United States; Department of Laboratory Medicine, Institute of Biomedicine, University of Gothenburg, Gothenburg, Sweden; Department of Immunology, Genetics and Pathology, Uppsala University, Uppsala, Sweden; Department of Medical Biology, University of Tromsö, Tromsö, Norway; Department of Pediatric Oncology and Hematology, Charité-Universitätsmedizin Berlin, Berlin, Germany; Department of Experimental Pediatric Oncology, University Children’s Hospital, Cologne, Germany; Center for Molecular Medicine (CMMC), Medical Faculty, University of Cologne, Cologne, Germany; Department of Microbiology, Cell and Tumor Biology, Karolinska Institutet, Stockholm, Sweden; Department of Medical Genetics and Cell Biology, Institute of Biomedicine, Sahlgrenska Academy, University of Gothenburg, Gothenburg, Sweden; Department of Cell and Molecular Biology, Karolinska Institutet, Stockholm, Sweden; Laboratory of Organic Chemistry II, Department of Chemistry, Faculty of Science, Hokkaido University, Sapporo, Japan; Princess Máxima Center for Pediatric Oncology, Utrecht, The Netherlands; Department of Human Genetics, Radboud University Medical Center, Nijmegen, The Netherlands; Department of Genetics and Pathology, University of Lund, Lund, Sweden; Hopp Children’s Cancer Center (KiTZ), Heidelberg, Germany; Division of Pediatric Neurooncology, German Cancer Research Center (DKFZ), Heidelberg, Germany; Laboratory of Biological Chemistry, Department of Chemistry, Faculty of Science, Hokkaido University, Sapporo, Japan; Department of Stem Cell and Regenerative Biology, Harvard University, Cambridge, USA

**Keywords:** Neuroblastoma, chromosome 17q, p53, WIP1, *PPM1D*

## Abstract

Majority of cancers harbor alterations of the tumor suppressor *TP53*. However, childhood cancers, including unfavorable neuroblastoma, often lack *TP53* mutations despite frequent loss of p53 function, suggesting alternative p53 inactivating mechanisms.

Here we show that p53-regulating *PPM1D* at chromosome 17q22.3 is linked to aggressive tumors and poor prognosis in neuroblastoma. We identified that WIP1-phosphatase encoded by *PPM1D*, is activated by frequent segmental 17q-gain further accumulated during clonal evolution, gene-amplifications, gene-fusions or gain-of-function somatic and germline mutations. Pharmacological and genetic manipulation established WIP1 as a druggable target in neuroblastoma. Genome-scale CRISPR-Cas9 screening demonstrated *PPM1D* genetic dependency in *TP53* wild-type neuroblastoma cell lines, and shRNA *PPM1D* knockdown significantly delayed in vivo tumor formation. Establishing a transgenic mouse model overexpressing *PPM1D* showed that these mice develop cancers phenotypically and genetically similar to tumors arising in mice with dysfunctional p53 when subjected to low-dose irradiation. Tumors include T-cell lymphomas harboring *Notch1*-mutations, *Pten*-deletions and p53-accumulation, adenocarcinomas and *PHOX2B-*expressing neuroblastomas establishing *PPM1D* as a *bona fide* oncogene in wtTP53 cancer and childhood neuroblastoma. Pharmacological inhibition of WIP1 suppressed the growth of neural tumors in nude mice proposing WIP1 as a therapeutic target in neural childhood tumors.

## INTRODUCTION

Childhood cancers differ from adult malignancies in terms of histopatological entities, biological and clinical features and molecular landscapes^1^. Pediatric tumors are much less complex in their molecular makeup and exhibit significantly less genomic abberations and mutations compared to adult cancers^2^. Despite this, childhood cancer is still the leading cause to death of disease in the developed countries. The tumor suppressor gene *TP53* is the most commonly mutated gene in cancer, and *TP53* mutations are detected in more than 50% of adult cance. Pediatric cancers on the other hand, exhibit significantly less *TP53* mutations^3^. Neuroblastoma, a childhood tumor of the peripheral nervous system exhibits infrequent *TP53* mutations^3^. However, p53 activity is commonly impaired and relapsed tumors demonstrate increased incidence of *TP53* mutations^4^. This suggests that inactivation of p53 is important for tumorigenesis and that alternative mechanisms for p53 attenuation are operating in neuroblastoma.

Segmental gain of chromosome 17q is the most common chromosomal aberration and correlates to poor prognosis in neuroblastoma^5-7^. Located within the gained regions of 17q in high-risk neuroblastoma is the Protein phosphatase magnesium-dependent 1 delta *(PPM1D)* gene which encode the nuclear serine/threonine phosphatase WIP1 (wild-type p53 induced phosphatase 1)^8^. WIP1 is a critical regulator of DNA damage response and cell cycle progression by its ability to regulate the activity of p53, ATM, CHK1/2 and other key molecules involved in DNA repair, cell cycle progression and apoptosis^9-13^. In concordance, *PPM1D* mutations, amplifications and WIP1 overexpression has been observed in various cancers^7,14-21^. This suggest that *PPM1D* is important for tumorigenesis although conclusive evidences for *PPM1D* being an oncogene inducing cancer is still missing.

Here we use multiomic approaches to understand genetic development of neuroblastom focusing on the most common aberration, gain of chromosome 17q, and the oncogenic features of *PPM1D* and p53 regulating WIP1 phosphataseWe show that neuroblastoma patients harbor frequent *PPM1D* copy number gain and gene fusions, in addition to activating truncating germ-line and somatic mutations. We further inferred that overexpression of WIP1 in transgenic mice, in conjunction with cellular stress, induce development of an array of tumors including neuroblastoma, demonstrating that the *PPM1D* is a *de novo* oncogene with the potential to induce tumorgenesis without the support of additional oncogene activities. Finally, we show that WIP1 is a potential clinical therapeutic target in childhood neural tumors.

## RESULTS

### Chromosome 17q-gain containing *PPM1D* is the most common genetic aberration in neuroblastoma with additional *PPM1D* copies acquired through clonal evolution

To explore the genetic aberrations in neuroblastoma, we used array-based comparative genomic hybridization (CGH) to detect genetic aberrations in 271 Swedish neuroblastomas. Gain of chromosome arm 17q was the most common genetic aberration observed and either segmental 17q gain or whole chromosome 17 gains were present in 82% of the tumors (**Figure S1A**). The majority of unfavorable neuroblastoma, harbored segmental gains of 17q whereas favorable neuroblastomas commonly display whole chromosome 17 gains. Neuroblastoma patients with segmental 17q gain show worse clinical outcome compared to those without 17q gain (5-year overall survival probability 48.0% vs. 82.8%; *P*=9.8×10^−9^; **Figure 1A**). The children with favorable neuroblastoma without segmental 17q-gain (or *MYCN* amplification and/or 11q-deletion) had favorable long-term survival at 8-20 years from diagnosis (>90%), in most cases without any treatment, compared to those with 17q-gain receiving active multimodal therapy with risk of late sequelae and still poor survival (<40%; **Figure S1B**), further validating 17q-gain as a prognostic marker.

**Figure 1.**
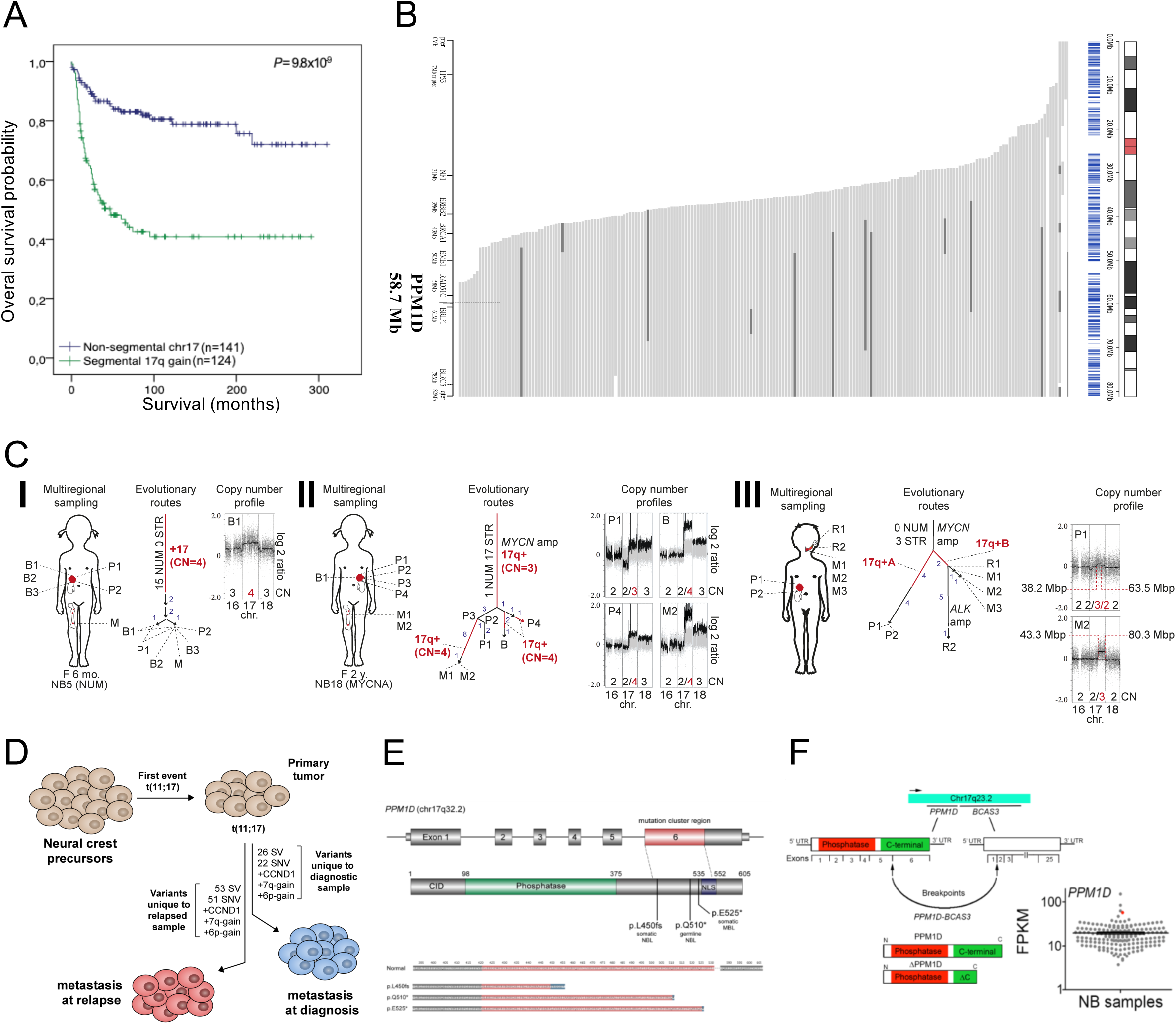
The *PPM1D* gene located on chromosome 17q is frequently altered in neuroblastoma. **A**. Gain of 17q correlates with poor survival in neuroblastoma. Neuroblastoma survival probability according to Kaplan-Meier analysis in a Swedish population-based patient material in relation to chromosome 17 status in the tumor tissue shows worse overall survival (OS) for children with 17q segmental gain (n=124; green line, 48.0% 5 year OS) vs children with no segmental 17q gain (n=141; blue line, 82.8% 5 year OS). For definition of the genomic profile groups (*Carén et al. 2010*). **B**. *PPM1D* is included in all cases of 17q gain. Summary of segmental gains of chromosome 17 in the extended Swedish neuroblastoma cohort (n=435). 208 samples displayed segmental gain of chromosome 17 as indicated by gray horizontal bars with regions displaying additional level of gain (more copies) indicated by black color. The dotted vertical line indicates location of the *PPM1D* locus in relation to the segmental 17q gains in the cohort. An additional 149 cases showed whole chromosome gain of chromosome 17 (not shown in the figure). **C**. Evolutionary trajectories of PPM1D gain. Multiregional tumor sampling reveals *PPM1D* copy number accumulation in high risk neuroblastoma. **(I)** Genome array analysis of pre-treatment biopsies (B1-B3), post-chemotherapy resection specimens (P1-P2) and concurrent bone marrow metastasis (M) from a female (F) 6-month-old (mo.) patient with a low risk neuroblastoma (NB), having a whole genome profile with only numerical (NUM) changes (I, left). Evolutionary reconstruction (I, center) shows 15 aberrations in the phylogenetic stem, i.e. present in >90% of tumor cells in all samples, followed by several additional whole chromosome changes at subclonal levels (arrows with numbers in blue type), distributed over a subset of samples. The stem aberrations include four copies (CN=4) of chromosome 17, as shown by a higher log2 ratio (I, right) than for the trisomic chromosomes (chr. 16 and 18). No further chromosome 17 aberrations were detected. **(II)** Multiregional array analysis of a *MYCN* amplified (MYCNA) high-risk NB in a 2-year-old (y.) with a metastatic relapse in the bone marrow indicates a stem dominated by structural (STR) rearrangements including segmental gain (CN=3) of the *PPM1D* region in 17q. Additional *PPM1D*/17q copies (CN=4) are gained in samples B, M1, M2 and P4 either clonally (>90% of tumor cells, red line) or subclonally (red arrow). **(III)** Another high-risk *MYCN* amplified NB demonstrates parallel evolution of *PPM1D*/17q gain in different tumor regions, as evidenced by different genomic breakpoints (III, right) in different sample sets, including two temporally distinct relapses (R1 and R2). **D**. Gain of 17q including PPM1D is the first and only common aberration in a high-risk metastatic neuroblastoma. Whole genome sequencing of different metastatic sites at diagnosis and relapse revealed an unbalanced translocation t(11;17) as the only common aberration whereas multiple unique but different genetic aberrations unique to respective sample were present in the two metastatic clones sequenced. **E**. Schematic representation of coding and protein sequence of *PPM1D*/WIP1 with mutations shown according location in protein and predicted amino acid sequences resulting from the *PPM1D* variants detected in neuroblastoma (NBL) and medulloblastoma (MBL) patients. Amino acids translated from exon 6 depicted in red with mutated amino acids depicted in blue. **F**. *PPM1D* expression levels are high in neuroblastoma tumor (red) bearing transcript of the *PPM1D*–*BCAS3* fusion, shown by FPKM (Fragments Per Kilobase per Million mapped reads).

Our data for the 271 analyzed Swedish neuroblastoma cases (**Figure S1A**) were extended with multiple samples to a total of 435 analyzed samples showed 208 tumors with segmental gain of chromosome 17q. Given the close propinquity to the proximal breakpoints associated with 17q gain and presence in all tumors with segmental 17q gain, the most prominent gene candidates with tumorigenic capacity are *RAD51C, PPM1D* and *BRIP1* (**Figure 1B**) out of which only *PPM1D* is a previously proposed putative oncogene (http://www.cancer.sanger.ac.uk/cosmic/mutation/).

To further delineate a candidate gene involved in cancer development and to investigate *PPM1D* in neuroblastoma progression, we used evolutionary trajectory analysis^22^. Neuroblastoma copy number status was investigated during tumor progression where we analyzed SNP array data of 100 tumor samples from 23 children with neuroblastoma^22^ with focus on chromosome 17. Most tumors (57%, 13/23) were found to harbor gain of 17q (17q+, always including *PPM1D* in the chromosomal gain) in >90% of the tumor cells in all samples, confining this aberration to the stem of evolutionary ideograms, without further change in copy number (**Figures 1C,I and Tables S1**). However, 30% (7/23) of the tumors showed a successive accumulation of 17q copies as tumors evolved regionally or from primary tumor to metastasis or relapse. In four of these cases, *PPM1D*/17q was gained already in the phylogenetic stem, followed by gain of additional copies of these genes as regional clones evolved (**Figure 1C, II**). In three cases, clones with *PPM1D*/17q gain emerged in subset of samples, where they expanded through selective sweeps to encompass all tumor cells in the samples (**Figure 1C, III**). The remaining 13% of patients (3/23) harbored a regionally fluctuating copy number status of *PPM1D*/17q that precluded conclusive evolutionary analysis. Among the 20 cases informative for evolutionary analysis, successive accumulation of *PPM1D/*17q copies was observed in more than half of high-risk patients (7/13) (**Table S1**). In contrast, additional changes in 17q copy number were not found in any of the low-risk patients, i.e. children <18 months with only numerical changes in the stem (0/7; *P*=0.0445; two-sided Fisher’s exact test). Thus, here we identify *PPM1D* as the strongest gene candidate involved in neuroblastoma progression.

In additional neuroblastoma cases with multiple tumor samples available for analysis we show that gain of 17q including *PPM1D* was the single first event in neuroblastoma development (**Fig 1D, Fig S1C** and **S1D)** in both a child without known predisposition and in a child with a germline PHOX2B mutation^23^.

To examine if mutations are present in neuroblastoma, we next performed whole exome sequencing (WES) or whole genome sequencing (WGS) of 73 neuroblastoma patient samples to screen for germline and somatic mutations. Notably, we identified a pathogenic truncating *PPM1D* mutation in exon 6 (c.1344_1345insT; p.L450fs) in a *MYCN*-amplified tumor sample obtained from an infant girl with neuroblastoma rapidly progressing after diagnosis from localized INSS stage 1 to metastatic INSS stage 4 (**Figures 1E and S1E**) with fatal clinical outcome despite active therapy. In addition, a *de novo* germline DNA mutation located in exon 6 (c.1528C>T; p.Q510*) resulting in a premature *PPM1D* truncation (**Figure 1E**) was identified in a patient diagnosed with a metastatic (stage 4, INSS/stage M, INRGSS) neuroblastoma of the left adrenal gland with bone metastases. The tumor was diagnosed in a 26 months old boy with history and symptoms similar to those previously reported in children with intellectual disabilities and *PPM1D* germline truncating mutations including significant dysmorphic signs (**Figure S1F**). The tumor had no *MYCN* amplification and the patient had a poor clinical outcome after relapse and progression despite multimodal clinical therapy. A detailed description of this patient is given in Star methods. Germline *PPM1D* mutations have previously not been reported in cancer but have been observed in children with intellectual disabilities and neurodevelopmental disorders^24,25^. Both these neuroblastoma-associated *PPM1D* mutations predictably give rise to C-terminal truncated variants of WIP1 similar to those described in other cancers (**Figure 1E**) (http://www.cancer.sanger.ac.uk/cosmic/mutation/). In addition, a gene fusion between *PPM1D* and the breast carcinoma amplified sequence 3 *(BCAS3)* gene at chromosome 17q23.2 was identified in a tumor from a patient enrolled in the Therapeutically Applicable Research to Generate Effective Treatments (TARGET) neuroblastoma dataset, which was associated with high expression of *PPM1D* and a predicted WIP1 isoform with a truncated C-terminal (**Figure 1F**).

### Overexpression of the *PPM1D* gene in neuroblastoma correlates with unfavorable clinical and biological features

Our data demonstrating that the *PPM1D* gene is altered in copy number and/or structure in the majority of high-risk neuroblastomas, prompted us to investigate the clinical prognostic and biological features of *PPM1D* gene expression in neuroblastoma. First, we show that higher expression levels of *PPM1D* was associated with several neuroblastoma risk factors, i.e. metastatic stage 4 disease, increased age at diagnosis (>18 months), *MYCN*-gene amplification and the INRG high-risk (HR) group as defined by combined clinical and biological features associated with unfavorable clinical outcome^26^ (**Figure 2A**). Highest level of *PPM1D* expression was observed in the non-*MYCN* amplified HR-neuroblastoma tumors (nMN HR; **Figure 2A**) mainly corresponding to 11q-deleted tumors that, according the overall genomic landscape, display the most frequent segmental 17q-gain in the *MYCN* non-amplified 11q deleted tumors (**Figure S1A**). Second, quantitative copy number (CN) values for *PPM1D* calculated from CGH array, WES and whole genome sequencing (WGS) was correlated to *PPM1D* expression. CN gains were separated into numerical gains that included whole chromosome 17 and segmental gains that included the sub-region containing *PPM1D*. Correlation analysis revealed a significant stepwise gene dosage dependent expression pattern of *PPM1D*, with the highest expression in tumors with segmental/partial 17q CN gains (**Figures 2A,B**). Moreover, high *PPM1D* expression was related to clinical outcome and children with neuroblastomas showing high expression had worse overall survival (OS; 59% vs. 83% at five years, p<0.001, **Figure 2C**, left panel) and event-free survival (EFS; 47% vs. 68%, p<0.001, **Figure 2C**, right panel). Third, low-risk neuroblastomas characterized by near-triploid DNA content and high TrkA expression (Type 1)^27^ were compared to more aggressive subtypes commonly having 17q gain with 14q- and/or 11q-deletions (Type 2A) or 1p-deletion and *MYCN*-amplification (Type 2B) with principal component analysis (PCA) on gene expression profiles from 30 primary neuroblastoma samples from two published microarray studies^28,29^. High expression of *PPM1D* was associated with aggressive Type 2A and Type 2B neuroblastomas, while low *PPM1D* expression was observed in Type 1 neuroblastomas (**Figure S2A**). Fourth, immunohistochemical staining showed consistent WIP1 expression in all 17q-gained neuroblastoma samples analyzed (**Figure S2B**). Taken together, these data suggest an increased aggressiveness of neuroblastoma and worse survival for patients with high WIP1-expressing tumors, in line with the predictions of the 17q gain survival data (**Figures 1A and S1B**) and the strong *PPM1D-*17q gain correlation (**Figures 1B,C, 2A,B and S2A**).To extend the investigations on the correlation of chromosome 17 gain and *PPM1D* expression levels, we included medulloblastoma that similar to neuroblastoma also presents frequent chromosome 17q gain or isochromosome 17^3,30^ suggesting that there may be a common gene that is involved in the development of both cancers. We analyzed a cohort consisting of 446 human medulloblastoma and 18 normal cerebellum cases and demonstrated that the highest PPM1D/WIP1expression was detected in the clinically unfavorable Group 3 and 4 subgroups (**Figures S2C, Table S2**), that are in line with previous findings^31^. Furthermore, we detected one PPM1D nonsense mutation in exon 6 (E525X; **Figure 1E**) in a medulloblastoma tumor of the WNT subgroup (**Figure S2E**), and amplifications of PPM1D in three patient samples belonging to the clinically unfavorable SHH (4-17 years) subgroup (**Figures S2D and S2F, G**). While, TP53 mutations are significantly associated with the SHH-group and most pronounced to this particular unfavorable SHH subgroup of intermediate ages and adverse outcome^32^, neither the PPM1D mutated nor the PPM1D amplified tumors in this analysis had TP53 mutations despite similar age- and prognostic features, indicating an alternative way of impairing p53 function through amplification of PPM1D in these medulloblastoma tumors.

**Figure 2.**
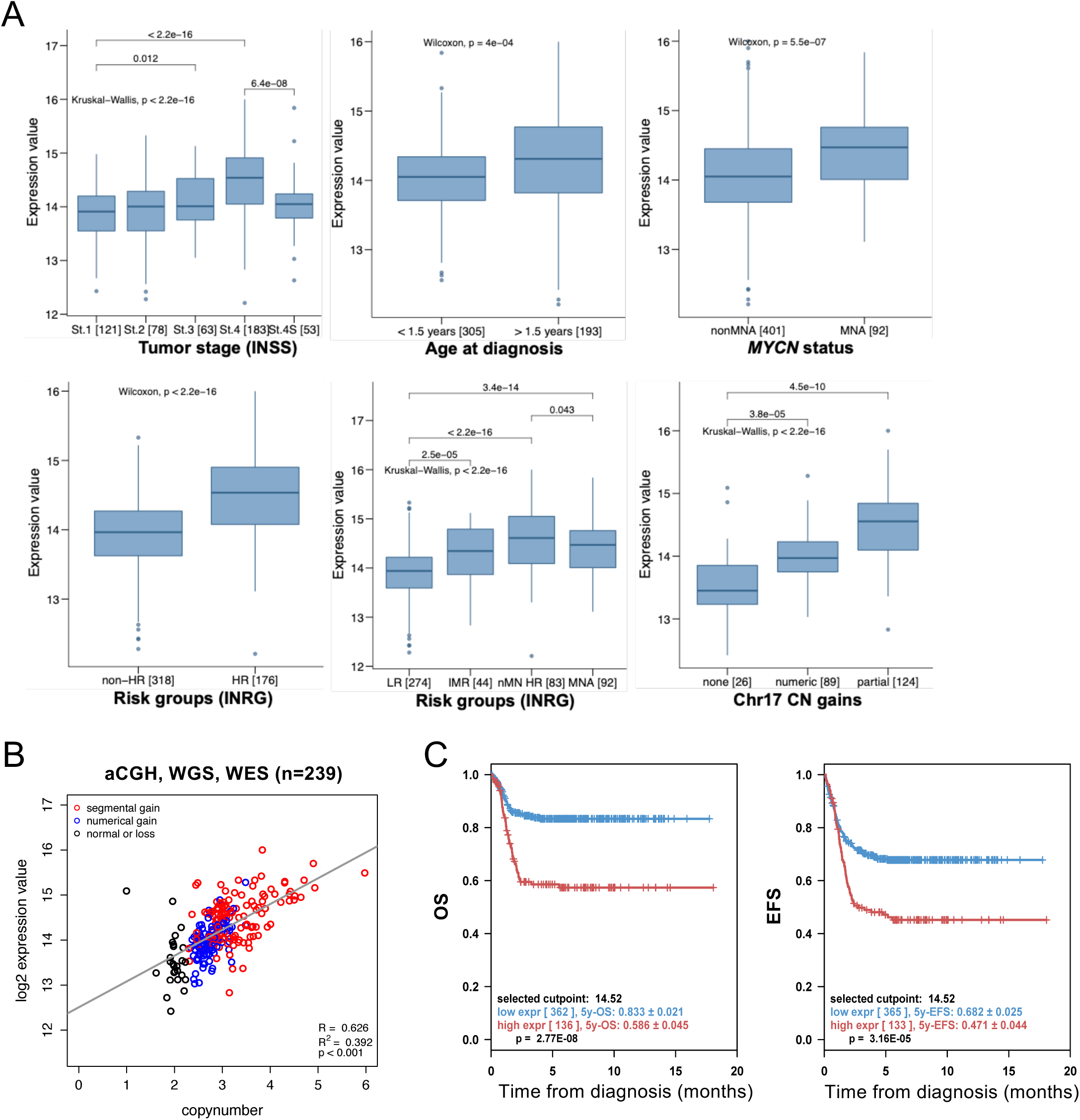
*PPM1D* expression correlates with unfavorable prognosis markers, decreased survival of neuroblastoma patients, and is gene-dosage dependent. **A**. Expression of *PPM1D* is associated with unfavorable prognostic markers. *PPM1D* expression is shown in box-and whisker plots for neuroblastoma tumor stages according to the INSS classification, age at diagnosis, amplification status of *MYCN*, INRG risk group classification (LR=low risk, IMR= intermediate risk, NMN HR= non-*MYCN* amplified high risk, MNA=*MYCN* amplified), and chromosome 17 copy number (CN) gains. **B**. Correlation analysis showing a gene-dosage dependent expression pattern of *PPM1D*. Quantitative copy number information for *PPM1D* based on CGH, whole exome (WES) and whole genome sequencing (WGS) data (x-axis) is shown against normalized log2 expression values from paired RNA-Seq data (y-axis). CN gains are highlighted and separated into numerical gains copying the whole chromosome 17 (blue) and segmental gains affecting only a sub-region involving *PPM1D* (red, min/max/mean size 2.4/70.8/35.8 Mb). Correlation and P-values were obtained by Pearson’s correlation coefficient. **C**. High expression of *PPM1D* is associated with adverse patient outcome. Kaplan–Meier survival estimates are shown for overall survival (OS, left) and event-free survival (EFS, right) in the whole cohort (n=498). P values were obtained by log-rank test. The cohort was dichotomized according to the optimal cut-off expression for *PPM1D*.

### *PPM1D* is important for neuroblastoma development and treatment resistance

To comprehensively assess the function of *PPM1D*/WIP1 in neuroblastoma as well as in medulloblastoma, we used preclinical models of the diseases. First, we examined the expression of *PPM1D* mRNA in a panel of neuroblastoma and medulloblastoma cell lines, as well as the breast cancer cell lines MCF-7 and BT-474 exhibiting *PPM1D* gene amplification^33^, and one sPNET cell line containing isochromosome 17. The cell lines expressed different levels of *PPM1D* mRNA that correlated to their genomic profile^34,35^ with highest expression in *PPM1D*-amplified cell lines and lowest in those without chromosome 17 aberrations (**Figures S3A, B**). All neuroblastomas expressed *PPM1D* mRNA, and the highest expression was observed in SK-N-DZ and IMR32, showing expression levels comparable to the *PPM1D*-amplified breast cancer cell lines MCF-7 and BT-474. Furthermore, the relative expression level of *PPM1D* mRNA was higher in medulloblastoma cell lines with 17q-gained aberrations (D283-MED, D458-MED and MEB-MED8A) compared to cell lines with normal 17q (DAOY, UW228-3 and PFSK-1), further demonstrating a gene-dosage effect on *PPM1D* mRNA expression (**Figures S3A, and S3C**).

We next used genome-scale CRISPR-Cas9 screening^36^, and demonstrated that *PPM1D* with the highest genetic dependency were found in *TP53* wild-type neuroblastoma cell lines compared to *TP53* mutated cell lines (**Figures 3A and 3B; Supplemental Table S3**). Other *TP53*-negative regulators such as *MDM2, MDM4* and *USP7* were ranked as number 3, 21 and 31, respectively. Among the top 40 ranked genes with largest difference between *TP53* wild-type and *TP53* mutated, an enrichment of genes involved in negative regulation of cell proliferation (FDR=0.0117), cell cycle process (FDR=0.0117), cellular response to DNA damage (FDR=0.05) and chromosome organization (FDR=0.05) (**Figure 3C**) was evident.

**Figure 3.**
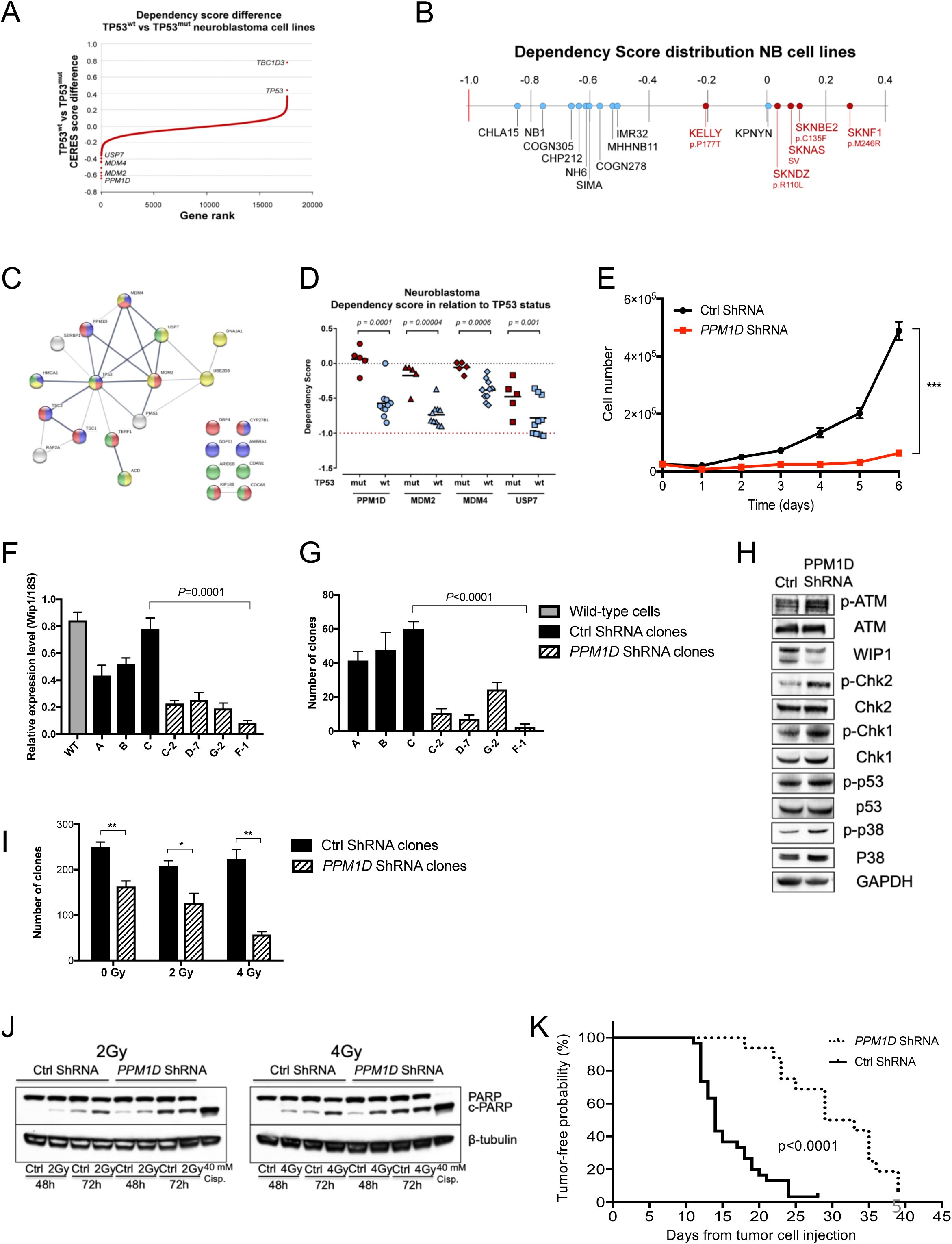
*PPM1D* expression is important for neuroblastoma. **A**. Wild type *TP53* neuroblastoma cells are highly dependent on *PPM1D* expression for survival. Genome-scale CRISPR-Cas9 screening showing ranked average difference in genetic dependencies between wild-type *TP53* and mutated *TP53* neuroblastoma cell lines. **B**. Dependency score showing high *PPM1D* dependency in wild-type *TP53* neuroblastoma cells. Wild-type *TP53* neuroblastoma cell lines (blue, high dependency) and *TP53* mutated cell lines (red, low dependency) with *TP53* status. **C**. STRING database analysis showing *PPM1D* dependency in wild-type TP53 neuroblastoma cells. Among the top 30 genes with largest different in CERES score there was enrichment of genes involved in negative regulation of cell proliferation (indicated in blue), cell cycle process (red), cellular response to DNA damage (yellow) and chromosome organization (green). The width of the edges corresponds to level of confidence (medium confidence STRING scores of 0.4; high confidence STRING score 0.7; and highest confidence STRING score 0.9). **D**. Neuroblastoma cells with wild-type *TP53* are dependent on *PPM1D* expression for survival. Dependency scores of *PPM1D, MDM2, MDM4*, and *USP7* in relation to *TP53* mutational status. **E**. Knockdown of *PPM1D* with shRNA impairs growth of neuroblastoma cell line SK-N-BE(2). Mean with S.D. of three independent experiments are shown (t-test day 6, SK-N-BE(2) *P*=0.0002. **F**. ShRNA-mediated knockdown of *PPM1D*. Normalized *PPM1D* mRNA expression in wild-type SK-N-BE(2) cells (grey bar) compared to three clones transfected with a non-silencing control shRNA (black bars, A, B, C) and four different *PPM1D* shRNA knockdown clones (hatched bars, C-2, D-7, G-2 and F-1). The expression of *PPM1D* mRNA was significantly lower in the clones transfected with the different *PPM1D* shRNAs compared to the control transfected clones (one-way ANOVA with Bonferroni correction *P*<0.05). Mean with S.D. of three determinants are shown. **G**. Knockdown of *PPM1D* inhibits colony-forming potential of neuroblastoma cells. Clonogenic assay of SK-N-BE(2) cells showing decreased colony formation in shRNA *PPM1D* knockdown cells (one-way ANOVA with Bonferroni post test *P*<0.0001) with lowest colony forming ability in the F-1 clone (t-test. P<0.0001). Mean with S.D. of nine determinants are displayed. **H**. Knockdown of *PPM1D* inhibits dephosphorylation of WIP1 target genes. Phosphorylation levels of WIP1 targets increased after *PPM1D* knockdown (clone F-1) compared to control transfected cells (clone C), as shown by western blotting. **I**. *PPM1D* downregulation sensitizes neuroblastoma cells to irradiation. *PPM1D* knockdown of SK-N-BE(2) cells showed an irradiation dose-dependent decrease in clonogenic forming ability compared to control transfected cells. Mean with S.D of three experiments are displayed (t-test, 0 Gy *P*=0.00486, 2 Gy *P*=0.0277, 4 Gy *P*=0.0015). **J**. ShRNA-mediated knockdown of *PPM1D* increases irradiation-induced apoptosis. Protein expression of the pro-apoptotic marker cPARP in *PPM1D* knockdown cells compared to control transfected cells 48 and 72 hours after exposure to irradiation, analyzed by western blot. β-tubulin was used as protein loading control. **K**. Knockdown of *PPM1D* delays neuroblastoma development. Clone F-1 and control clone C were injected in NMRI *nu/nu* mice (shRNA; n=8, control; n=15), 5 million cells sc. bilaterally. Tumor development was significantly delayed (log-rank test *P*<0.0001) showing median tumor development (0.100 mL) to be more than doubled (33 days median, vs. 15 days) after *PPM1D* downregulation (dashed line) compared to animals injected with cells transfected with the non-silencing control shRNA (black line).

Focusing on *PPM1D, MDM2, MDM4* and *USP7*, all of which with pharmacological inhibitors that are in preclinical testing or clinical trials, we showed that neuroblastoma cell lines have preferential dependency for all four genes in *TP53* wild-type cell lines (**Figure 3D**). In cell lines of medulloblastoma origin, *MDM2* showed largest difference in dependency score with negative regulators *PPM1D, USP7* and *MDM4* ranked 6, 41 and 64, respectively (**Figures S3E and S3H**). Among the top 40 genes there is a functional enrichment of pathways associated to cell cycle (FDR=0.000119), chromosome organization (FDR=0.00065) and nucleotide biosynthesis processes (FDR=0.003) (**Figure S3G**). As expected from the ranked differences between *TP53* wild-type and *TP53* mutated medulloblastomas, only *PPM1D* and *MDM2* showed differences in genetic dependency (**Figure S3H**). When comparing the genetic vulnerability of *PPM1D, MDM2, MDM4* and *USP7* in neuroblastoma cells to all the other cancer cell lines in the CRISPR Avana dataset, both *PPM1D* and *MDM4* showed strong selective dependency in neuroblastoma (**Figure S3D**) highlighting the essentiality of intact p53 (**Figure S3F**).

To study the potential tumorigenic function of *PPM1D*/WIP1 in neuroblastoma we next used genetic manipulation through stable knockdown of short-hairpin RNA (shRNA) against *PPM1D*. The majority of neuroblastoma cell lines transfected with *PPM1D* shRNA were non-viable compared to scrambled controls (data not shown), whereas cells that were viable after transfection showed reduced proliferation and increased H2AX phosphorylation, suggesting an effect on cell viability and genomic integrity (**Figures 3E and S3I**). The multi-resistant neuroblastoma cell line SK-N-BE(2), isolated from a patient with recurrent stage 4 neuroblastoma was chosen for further analysis since this was one of the few cell lines that was viable after *PPM1D* shRNA knockdown (**Figure 3E**). Analyzing different shRNAs constructs directed against *PPM1D* mRNA in SK-N-BE(2) cells established *PPM1D*-shRNA F-1 as the most effective in reducing *PPM1D* mRNA levels (**Figures 3F,G**). *PPM1D* knockdown was also confirmed by increased phosphorylation of WIP1 target proteins including ATM, CHK1, CHK2, p53 and p38, all involved in cell cycle regulation and the DNA-damage response (DDR) (**Figure 3H**). To test the role of *PPM1D* in cellular stress we subjected SK-N-BE(2) cells to irradiation, and showed that cells lacking *PPM1D* exhibited increased sensitivity to irradiation compared to control cells (**Figure 3I**). Moreover, higher levels of PARP-cleavage were detected in *PPM1D* shRNA transfected cells compared to controls, suggesting an increased vulnerability to apoptosis following irradiation (**Figure 3J**). Furthermore, to evaluate the effect of *PPM1D* knockdown on tumor development *in vivo*, SK-N-BE(2) cells with either *PPM1D*-knockdown or control shRNA were injected subcutaneously in mice and tumor growth was compared. Tumor development was significantly delayed showing median time to tumor development (0.1 mL tumor volume) to be more than doubled (33 days median, vs. 15 days) in the *PPM1D* shRNA knockdown group compared to animals injected with control shRNA-transfected cells (*P*<0.001) (**Figure 3K**).

### Mice carrying the *PPM1D* transgene develop tumors in response to cellular stress

Demonstrating that *PPM1D*/WIP1 is a strong gene candidate in the development of neuroblastoma and has a critical function in the growth of tumors and their response to cellular stress, we next investigated the oncogenic potential of *PPM1D*. Hence, we constructed genetically engineered C57BL/6N mice to overexpress WIP1 through pronuclear injection of the human *PPM1D* gene controlled by the rat tyrosine hydroxylase (TH) promoter (**Figure S4A**). Three founder lines were generated with intact transmission of the *PPM1D*-transgene to following generations.

Elevated WIP1 protein expression was detected in spleen, ovary and intestine in mice heterozygous for *PPM1D* compared to wild-type mice (**Figure S4B**). However, no increase in tumor development was observed in these *PPM1D-*transgenic mice. We therefore backcrossed the C57BL/6N*PPM1D* +/- mice for eight generations with 129×1/SvJ mice to establish a near 100% 129×1/SvJ background which is more prone to tumor formation. In addition, given the wide-ranging actions of WIP1 in DNA damage, it is likely that overexpression of WIP under cellular stress has the potential to induce neoplasms arising in response to extrinsic cellular DNA damage. Therefore, *PPM1D*-overexpressing animals and their wild-type littermates were exposed to 4.5 Gy of sub-lethal whole-body irradiation at different ages (1-314 days old). Irradiated transgenic mice carrying the *PPM1D* gene had significantly higher tumor incidence (34% tumor probability) compared to wild-type mice (7.8%) (*P*<0.0001), 100-300 days post-irradiation (**Figure 4A**). The majority of cancers detected in *PPM1D*-transgenic mice were similar to those reported in irradiated p53-mutant mice post-irradiation^37-39^, indicating a similar tumor phenotype caused by p53 impairment rather than p53 absence. Thymic lymphoblastic lymphoma was the most frequently malignancy observed (n=41; **Figures 4B,C, Table S4**), compared to four thymic lymphomas detected in irradiated wild-type mice (*P*<0.001). Other malignancies manifested in *PPM1D*-transgenic mice were leukemia/lymphomas (n=9), different adenocarcinomas/adenomas (n=18) and sarcomas (n=4), (**Table S4**). Importantly, mice also developed neural crest-derived PHOX2B-expressing primary tumors of the adrenal gland with neuroblastoma-like features mixed with paraganglioma/pheochromocytoma-like morphology (one of these mice also displayed PHOX2B-expressing neuroblastoma-like metastasis) (**Figure 4D**), further supporting that *PPM1D* might be a driver of neuroblastoma development. We also identified mice with multiple primary tumors, one mouse with an adenocarcinoma in the lacrimal gland and osteosarcoma with lung metastasis, a mouse with T-cell lymphoma and ovary carcinoma, lung adenocarcinoma and a mouse with a gastric adenocarcinoma and leukemia (**Table S4**). The odds ratio of developing cancer in mice with the *PPM1D* transgene compared to wild-type mice was 6.3 (95% confidence interval 2.7-14.2). Comparison of age at irradiation and time to tumor development disclosed a positive correlation, showing that younger *PPM1D*-positive animals were more susceptible to developing thymic lymphomas in response to DNA damage (*P*<0.01), the majority (38 out of 41) of which arose <365 days post-irradiation (**Figure 4B**). Protein expression of thymic lymphomas showed high WIP1 expression and increased phosphorylation of p53 compared to non-irradiated thymus from *PPM1D* -positive and wild-type mice (**Figure S4B**). Other tumor manifestations observed in multiple *PPM1D*-positive mice were spleen, liver, lung and lymph node metastases (**Figure S4C and Table S4**).

**Figure 4.**
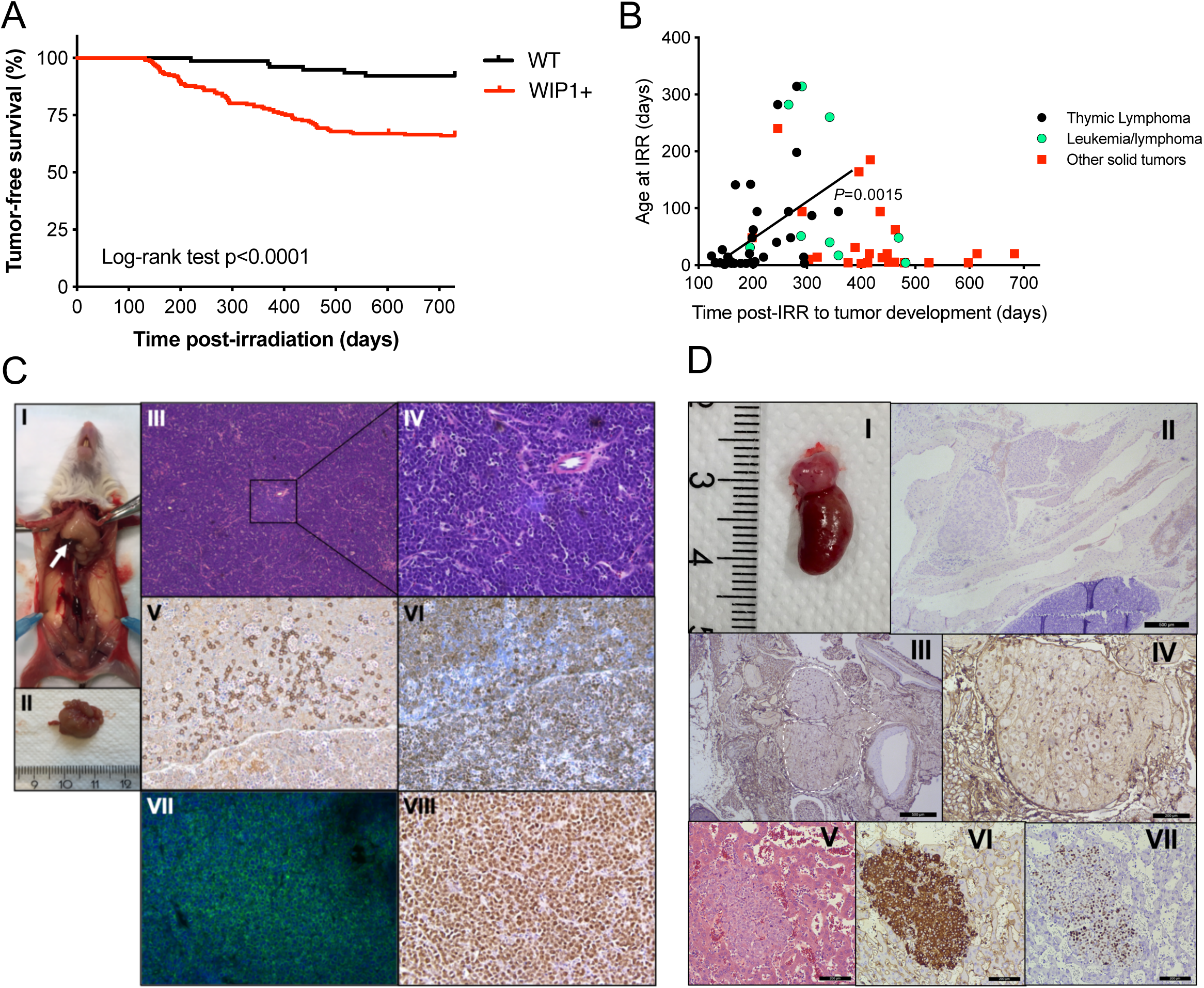
*PPM1D*/WIP1-transgenic mice have increased carcinogenic susceptibility compared to wild-type mice following irradiation. **A**. Kaplan-Meier analysis of irradiated *PPM1D*/WIP1-transgenic mice with the endpoint defined as detection of tumors. Mice were either homozygous or heterozygous for the human *PPM1D* gene. WIP1 positive transgenic mice (n=210) and their wild-type littermates (n=79) with different degrees of 129×1/SvJ-strain background were subjected to one 4.5 Gy sublethal whole-body irradiation dose at different days of age (1-314 days old). Mice that were positive for the human *PPM1D* gene developed frequently more tumors compared to wild-type mice (Log rank test, *P*<0.0001) following irradiation. Among the *PPM1D*/WIP1-positive mice 72 mice developed tumors compared with six mice in the wild-type group (Fisher’s exact test, *P*<0.0001). The odds ratio of developing cancer in the *PPM1D*/WIP1-positive group compared to the wild-type group was 6.3 (95% confidence interval 2.7-14.2). **B**. Time to tumor development of thymic lymphomas post-irradiation (IRR) correlated positively with age at irradiation (Pearson, *P=*0.0015. Spearman, *P*=0.001); the majority of lymphomas manifested <300 days. There was no correlation between age at irradiation and time to tumor development in mice diagnosed with other solid tumors (Pearson *P*=0.0658, Spearman *P*=0.1589) and mice diagnosed having leukemia/lymphoma (Pearson *P*=0.3651, Spearman *P*=0.1977) **C**. Thymic lymphomas were the most common disease manifestation in irradiated human *PPM1D*/WIP1-positive mice. Representative image of thoracic tumor *in situ* indicated by the arrow (**I**) and after dissection (**II**). Microscopy at 4X (**III**) and 20X (**IV**) magnification after hematoxylin and eosin (H&E) staining showed atypical lymphoblastic cells having intense mitotic activity. **V)** Infiltration of B220-positive cells. **VI**-**VII)** The majority of cells are positive for the pan-T cell marker CD3. **VIII**) Staining for Ki-67 was highly positive, indicating high proliferative activity. The thoracic organs, mediastinum (heart base and hilum of the lung), thymus tumor, lymph nodes displayed extensive infiltrates of neoplastic and highly malignant lymphoblasts consistent with lymphoblastic lymphoma. **D**. Adrenal neuroblastic tumor and liver metastasis from a *PPM1D*/WIP1 transgenic mouse **I**) Macroscopic view of the adrenal tumor. **II)** Hematoxylin/Eosin staining of the adrenal tumor. **III-IV)** PHOX2B-staining was positive indicating neural crest origin of this neuroblastoma-like tumor. **V-VII)** Liver metastasis with Hematoxylin/Eosin-staining, Synapotophysin-staining and PHOX2B-staining, respectively.

### *PPM1D*-derived mouse tumors show frequent *Notch1* and *Pten* mutations and activation of Notch signaling

To characterize the tumors that developed in *PPM1D*-transgenic mice, we performed WES on *PPM1D*-derived T-cell lymphomas. Our data showed an average of 18 (range 9-34) somatic protein-changing mutations compared to non-irradiated wild type control mice. Thymic tissues from the two control groups, irradiated wild-type mice, and irradiated *PPM1D*-positive mice without tumors, showed an average of two (range 1-4) and three (range 1-5) somatic protein changing mutation variants, respectively, similar to non-irradiated non-tumorigenic wild type mice (**Table S5**). These data indicate that irradiation alone was not responsible for the accumulation of DNA mutations observed in the tumors. Among the thymic lymphomas, we found *Pten* and *Notch1* to be recurrently altered (**Figure 5**). In total, 15 activating *Notch1* mutations were detected in 9 of 15 lymphomas (60%) and among these, five tumors had more than one mutation of the *Notch1* gene. Similar to human cancers, the *Notch1* missense variants were found to cluster in the heterodimerization domain (HD) while frameshift or nonsense variants were clustered to the proline-glutamic acid-serine-threonine (PEST) domain (**Figure 5A**). Of the 15 sequenced lymphomas, *Pten* aberrations were detected in 12 cases (80 %) consisting of homozygous deletions (9 cases), missense variants (2 cases) and frameshift insertion (1 case) (**Figure 5B**). The three missense and frameshift variants all showed high level variant allele fraction (variant allele fraction 85-97%) suggesting a near homozygous presence. Mutations of *Pten* and *Notch1* were co-occurring in seven cases (46%) (**Table S5**). In addition to *Notch1* and *Pten* mutations, aberrations in genes previously associated to human cancers were also observed in *Ikzf1, Kras, Rac2, Trp53, Ctnnb1, Gli3, Zfp36l2, Mpdz, Dd3x, Foxp1, Pin1* and *Ezh2* (**Table S5**). No mutations were found in *Fbxw7, Notch2, Notch3* or *Notch4* which also have been associated to thymic lymphomas development in radiation-induced murine tumors.

**Figure 5.**
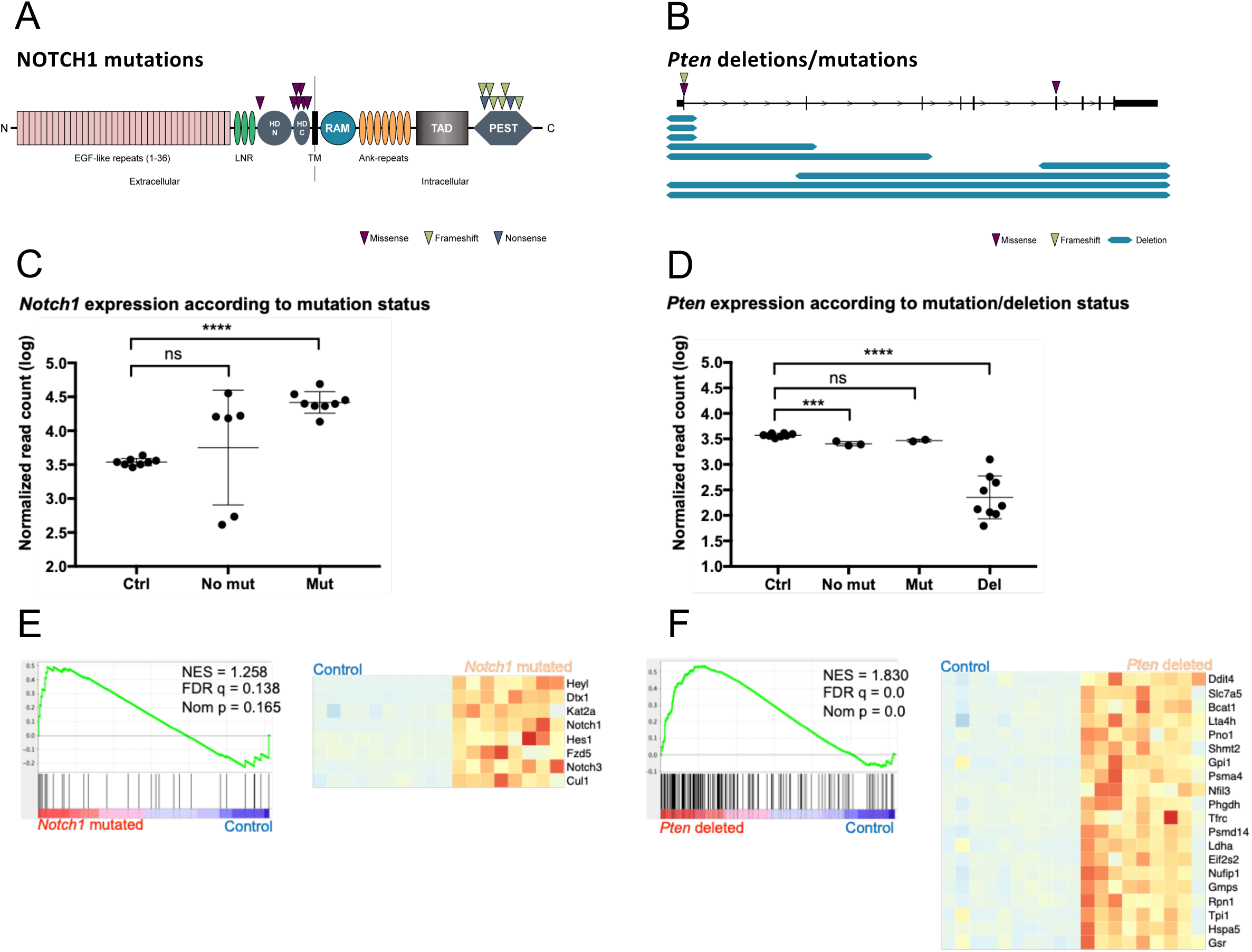
*PPM1D*-induced thymic lymphomas show typical *Notch1* and *Pten* aberrations. **A**. Schematic overview of the NOTCH1 protein showing the distribution of mutations in 15 whole-exome sequenced lymphomas. Purple triangles indicate missense mutations in the intracellular heterodimerization (HD) domain, whereas green and blue triangles indicate frameshift or nonsense mutations in the extracellular PEST domain. LNR: Lin/NOTCH repeats; HD: heterodimerization domain; TM: transmembrane domain; RAM: RBP-JK associated molecule region; TAD: transactivation domain; PEST: sequence rich in proline, glutamic acid, serine, and threonine. **B**. Diagram of the *Pten* gene showing the size and distribution of *Pten* deletions/mutations in lymphomas. The location of the point mutations are shown with purple and green triangles above the gene diagram. The blue lines below the gene diagram shows the size of the deletions identified by exome sequencing. **C**. Expression of the *Notch1* transcript in controls and tumors with and without *Notch1* mutation. The y axis shows the log value of the number of RNA-seq reads mapping to the *Notch1* gene. The difference between controls and tumors without *Notch1* mutations is not significant (ns), whereas *Notch1* expression is significantly higher in the mutated tumors (adjusted p value < 0.0001; Bonferroni multiple comparisons test). **D**. Expression of *Pten* in controls and tumors with and without *Pten* mutation/deletion. Expression is significantly lower in tumors harboring deletions than in controls (adjusted p value 0,0001, Bonferroni multiple comparisons test). **E**. Gene set enrichment analysis comparing *Notch1* mutated tumors to controls demonstrated a significant upregulation of genes related to Notch signalling (MSigDB “Hallmarks” gene set). Left panel: Enrichment plot from GSEA. Right panel: Heatmap showing the expression (z scores) of core enriched genes. Red color indicates positive z scores; blue color indicates negative z scores. **F**. Gene set enrichment analysis comparing *Pten* deleted tumors to controls demonstrated upregulation of genes related to the Mtorc1 pathway (MSigDB “Hallmarks” gene set). Left panel: Enrichment plot from GSEA. Right panel: Heatmap showing the expression (z scores) of the top 20 core enriched genes. Red color indicates positive z scores; blue color indicates negative z scores.

We also performed RNA-sequencing on the tumors and analyzed them using Principal component analysis (PCA) and unsupervised hierarchical clustering based on the 200 most variable genes (**Figure S5A, B**). Differential gene expression analysis (**Table S5**) identified 4138 upregulated and 4378 downregulated genes in tumors when compared to controls. Tumors harboring *Notch1* mutations had significantly higher expression of the *Notch1* gene compared to controls (**Figure 5C**). Also, as expected *Pten* expression was significantly lower in tumors harboring deletions of the gene (**Figure 5D**). Likewise, gene set enrichment analysis (GSEA) showed corresponding increased expression of genes involved in Notch- and Mtorc1-signalling in *Notch1* or *Pten* mutated tumors compared to controls (**Figure 5E,F**). Thus, tumor development was phenotypically and genetically similar to tumors from p53-impaired mice^38,40^ harboring *Notch1*-mutations, *Pten*-deletions and wild-type p53-accumulation.

### WIP1 is a therapeutic target in neuroblastoma

Having established that *PPM1D*-knockdown reduces tumor formation in mice and that high expression of WIP1 is a strong prognostic factor for poor survival in neuroblastoma, we investigated the effects of compounds inhibiting WIP1 activity in neuroblastoma. To assess the cytotoxic effects of different WIP1 inhibitors on cell viability, six neuroblastoma cell lines and the *PPM1D*-amplified breast cancer cell line MCF-7 were treated with different concentrations of the WIP1 inhibitors SL-176, SP-001 or CCT007093^41-44^. Treatment of neuroblastoma cell lines showed highest sensitivity to the specific WIP1-inhibitor SL-176 regardless of p53 or Mdm2 status. SL-176 displayed the lowest mean of IC_50_ value of the three tested WIP1 inhibitors (*P*<0.0001) (**Figures 6A, S6A and Table S6**).

**Figure 6.**
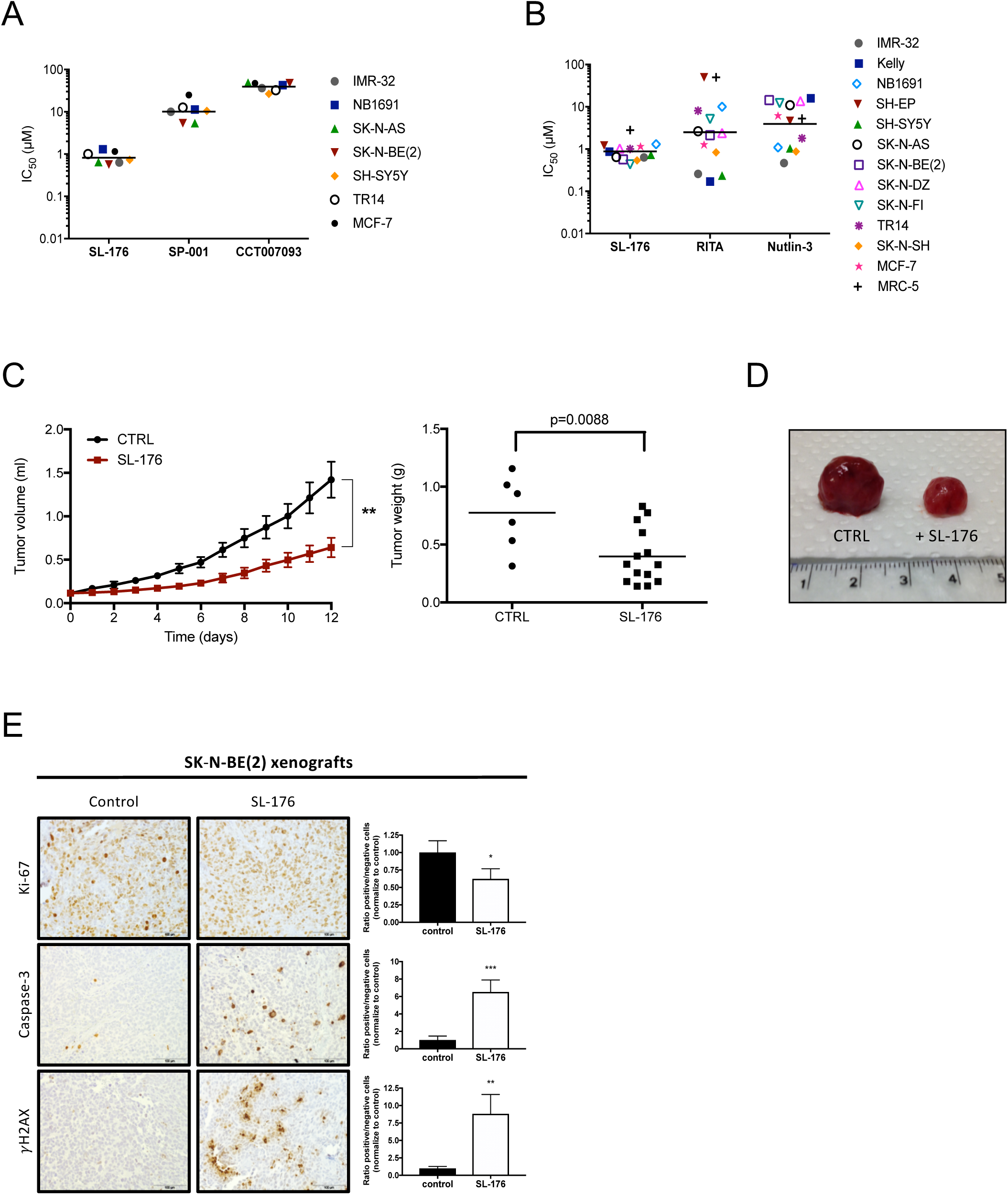
The WIP1 phosphatase inhibitor SL-176 suppresses neuroblastoma and medulloblastoma growth *in vitro* and *in vivo*. **A**. SL-176 is the most efficient inhibitor of tumor cell growth. IC_50_ values for six neuroblastoma cell lines and the *PPM1D*-amplified breast cancer cell line MCF-7 exposed to the WIP1 inhibitors SL-176, SP001 or CCT007093. Horizontal lines indicate mean. IC_50_ values were calculated from cell viability assays performed at least three times. SL-176 displayed the lowest mean of IC_50_ value of the three tested WIP1 inhibitors (one-way ANOVA on log IC50 *P*<0.0001, Bonferroni post-test: SL-176 vs SP-001 *P*<0.0001, SL-176 vs CCT007093 *P*<0.0001). **B**. SL-176 is a potent inhibitor of neuroblastoma cell growth. IC_50_ values for eleven neuroblastoma cell lines, the breast cancer cell line MCF-7 and the fibroblast cell line MRC-5 exposed to the specific WIP1 inhibitor SL-176, the p53-MDM2 interaction inhibitor RITA or the MDM2 antagonist Nutlin-3. Horizontal lines indicate mean. SL-176 displayed the lowest mean of IC_50_s in the neuroblastoma cell lines of the three tested compounds (mean IC_50_ for SL-176: 0.77 µM, RITA: 2.0 µM and Nutlin-3: 3.7 µM), however a significant difference was only evident between SL-176 and Nutlin-3 (one-way ANOVA *P*=0.029, Bonferroni post-test SL-176 vs. Nutlin-3 *P*=0.026). IC_50_ values were calculated from results from cell viability assay WST-1 performed at least three times. **C**. SL-176 inhibit neuroblastoma growth *in vivo*. Nude mice were injected with neuroblastoma SK-N-BE(2) cells to form xenografts on the flank. Daily i. p. injections of SL-176 (n=14) for 12 days, starting at tumor volume 0.1 mL, compared to no treatment (CTRL, n=6) showed that WIP1 inhibition through SL-176 significantly impaired the growth of neuroblastoma xenografts (t-test, day 12 *P=*0.002) (left, horizontal bars, mean weight), and significantly reduced tumor weight after 12 days (*P*=0.0088) (right). Mean with S.E.M. are displayed. **D**. Representative photograph of dissected neuroblastoma xenograft tumor in comparison. **E**. SL-176 decrease proliferation, induce apoptosis and activate γH2AX in xenograft tumors. Immunohistochemical analysis of SK-N-BE(2) xenograft tumors. Tumor sections were stained with anti-Ki-67, anti-Caspase 3, and anti γH2AX antibodies. Representative examples of immunostaining are shown. Images were acquired at 400X magnification. Identification and quantification of positive and negative cells was carry out with ImageJ software.

To further investigate the potency of the WIP1 inhibitor SL-176 on tumor cell viability in monotherapy, a panel of cancer cell lines was treated with different concentrations of SL-176, the p53 modulator RITA, and the MDM2 antagonist Nutlin-3. SL-176 was the most efficient drug against the neuroblastoma cell lines (mean IC_50_ for SL-176: 0.77 µM, RITA: 2.0 µM and Nutlin-3: 3.7 µM), (**Figure 6B, S6C and Table S6**). SL-176 had similar effects on medulloblastoma cell viability as the p53 inhibitors RITA and Nutlin-3. However, no significant differences between the mean IC_50_ values in medulloblastoma cell lines were observed (mean IC_50_ for SL-176: 1.1 µM, RITA: 0.41 µM and Nutlin-3: 3.4 µM, *P*=0.26). The non-cancerous cell lines, MRC-5 derived from human fetal lung fibroblasts and the mouse neural progenitor cell line C17.2 were less sensitive to SL-176 (mean IC_50_ for SL-176: 22 µM, RITA: 9.5 µM and Nutlin-3: 2.0 µM (**Figures S6B, S6C and Table S6**).

Next, we investigated the anti-tumorigenic effect of SL-176 monotherapy in a preclinical *in vivo* model of neuroblastoma Tumor growth inhibition was observed after 1 day of treatment (SK-N-BE(2); *P*=0.01) (**Figure 6C**) while tumor size and weight at the end of the experiment was significantly smaller in SL-176-treated mice compared to control mice (SK-N-BE(2); *P*<0.01, DAOY; *P*<0.05) (**Figures 6C, D**). No adverse effects of SL-176 were observed in the treatment groups. SL-176 treated tumors showed increase in active caspase-3 and phosphorylation of the DNA repair protein γH2AX, whereas reduced levels of the proliferation marker Ki-67 was observed (**Figure 6E**). Similarly, SL176 also suppressed the growth of established medulloblastoma xenografts in nude mice (DAOY; *P*=0.02, **Figures S6D, E**) with increased expression of active caspase-3 and γH2AX, and decreased Ki-67 expression (**Figure S6F**). Taken together, our data supports WIP1 as a druggable target in neuroblastoma and medulloblastoma.

## DISCUSSION

Neuroblastoma is a childhood tumor of the developing peripheral nervous system, which despite intensified multimodal therapy still have a poor outcome when compared to pediatric cancers in general. For further improvement of management, clinical care and chances of cure for these patients, advancements in molecular understanding of tumor biology is essential. We therefore investigated prevalent genetic aberrations in neuroblastoma as well as for medulloblastoma and our data propose the p53 regulating phosphatase WIP1 encoded by the chromosome 17q-located *PPM1D* gene as a novel druggable target for both these pediatric cancers.

Segmental gain of 17q is the most common chromosomal aberration and predictor of poor prognosis detected in neuroblastoma and medulloblastoma^5,6,45^. Also, gain of chromosome 17q is frequently found in tumors of epithelial, neural and hematopoietic origin^46-51^. This suggests that one or multiple genes important for tumorigenesis are located on 17q. Several cancer-associated genes have been identified on chromosome 17q that besides *PPM1D* also include *EME1, BRCA1, ERBB2, NF1, RAD51C, BRIP1* and *BIRC5*. However, our analysis of 271 Swedish neuroblastoma samples shows that only *RAD51C, PPM1D* and *BRIP1* are included in the shortest region of overlap of 17q gains. These genes were also included in the 17q23.2 chromosomal amplifications detected in three SHH-derived medulloblastomas and are similar to observations made in breast cancer^19,52,53^. We also show an accumulation of *PPM1D* gene copies in metastatic and relapsed neuroblastomas compared to low-risk tumors suggesting a clonal evolution of *PPM1D* in high-risk neuroblastoma. High expression of *BIRC5*, at chromosome 17q25.3, encoding the anti-apoptotic protein Survivin has been correlated with poor prognosis in neuroblastoma, whereas in medulloblastoma conflicting results regarding the importance of Survivin have been reported^54-57^. We detected two somatic *PPM1D* mutations, one each from the neuroblastoma and medulloblastoma cohort and one *de novo* germline mutation in one additional neuroblastoma patient. The detected *PPM1D* mutations were all truncating gain-of-function variants and located in exon 6, similar to observations in numerous other malignancies^19,52,53,58^. *PPM1D de novo* germline mutations have previously not been described in cancer. However, in children with intellectual disability syndrome, *PPM1D* germline mutations have been described to occur in the 5^th^ and 6^th^ exons. Notably, an identical c.1528C>T (p.Gln510*) somatic mutation has been reported in a malignant melanoma^59^. The vast majorities of reported somatic *PPM1D* mutationsare located in exon 6 and result in a truncated WIP1 protein with a proposed higher oncogenic capacity compared to the full-length WIP1 protein^19,52,60^. We also detected a gene fusion of *PPM1D* and *BCAS3* in a neuroblastoma patient, predicted to generate a C-terminal truncated WIP1 protein and resulting in high levels of WIP1 expression. Similarly, a gene fusion between *PPM1D* and *C1QTNF1* in a Ewing sarcoma patient resulting in high WIP1 expression has been reported (https://pecan.stjude.cloud). Also, a *PPM1D* fusion with an intragenic region of *RPSK6B1* and a *PPM1D-ZNS655* fusion has been reported in a diffuse cerebellar glioma patient and an AML patient, respectively, without any described effects on WIP1 expression (*https://pecan.stjude.cloud)*. Together these observations suggest that the *PPM1D* gene is frequently altered in a variety of cancers and has important functions during tumorigenesis.

To functionally test the importance of *PPM1D* in neuroblastoma and medulloblastoma we investigated the effect of genetic or pharmacological inhibition of WIP1 and demonstrated that blocking the expression or activity of WIP1 suppressed both neuroblastoma and medulloblastoma growth *in vivo*. These findings together with similar observations^7,61,62^ further support that WIP1 is important for the development and progression of these neural tumors.

WIP1 is a homeostatic regulator of the DNA damage response (DDR) cascade by dephosphorylating and inactivation of ATM, ATR, CHK1, CHK2 and DNA-dependent protein kinase catalytic subunit^63,64^. This and concurrent WIP1-mediated inactivation of p53 will reduce the fidelity of overall DNA repair mechanisms and induce accumulation of DNA aberrations which is a prerequisite for tumorigenesis. Accordingly, we observed increased phosphorylation of DDR proteins and H2AX enhancing the sensitivity to irradiation of *PPM1D* knocked-down neuroblastoma cells. Although mutations of *TP53* are not commonly detected at time of diagnosis in neuroblastoma, the p53 activity is recurrently compromised in these tumors^27,65^ and p53 inactivation has been shown to contribute significantly to neuroblastoma development in specific animal models^66,67^. Individuals with the cancer predisposition syndrome Li-Fraumeni caused by germline mutations of *TP53* may occasionally develop medulloblastoma^30,68^. Although uncommon, primary somatic mutations of *TP53* have been linked to the anaplastic medulloblastoma subset^69^ and more recently it was shown that *TP53* mutations have subgroup-specific prognostic implications with particular enrichment in the childhood SHH-subset linked to a poor clinical outcome^70^. Furthermore, wild type p53 inactivation in medulloblastoma has been suggested to be related to aberrations in the p53-ARF pathway including *MDM2* and *PPM1D*/WIP1^14,71^. Three of our investigated medulloblastoma samples contained *PPM1D* amplification. All three were detected in the SHH subset in child patients and none had *TP53* mutations, which is frequently observed in this unfavorable subgroup of medulloblastoma^32^. This further validates WIP1 as an important player for tumor development in cancers with impared p53. Also neuroblastoma incidence is increased in Li-Fraumeni families and in particular the specific common p.R337H mutation has been shown to increase neuroblastoma development^72,73^.

In primary neuroblastoma, mutations of genes of the p53 pathway are rare, occurring in roughly 5% of the cases^74^. By contrast, *TP53* mutations appear to be more frequent in relapsed neuroblastoma^74^ and in neuroblastoma cell lines established at relapse, indicating a role in the development of a therapy resistant phenotype^71,75,76^. Several additional mechanisms for p53 inactivation in neuroblastoma have been described including *MDM2* gene amplification or increased *MDM2* expression mediated by *MYCN* and hypermethylation or deletions of *CDKN2A*, miR-380-5p mediated repression of p53 expression or p53 inactivation by the methyltransferase SETD8^4,77-80^. The demonstration that *Ppm1d*-deficient mice significantly delayed the formation of ERBB2-induced mammary tumors and that *Ppm1d* null embryonic mouse fibroblast were resistant to transformation by RAS, ERBB2 and c-MYC^81^ prompted us to establish a genetically modified mouse model overexpressing WIP1. These mice developed tumors of different origin including neuroblastoma after low-dose irradiation compared to wild-type control mice. The spectrum of observed tumors was highly similar to tumors observed in p53 deficient mice or p53 deficient mice receiving irradiation^37-40,82-84^. Interestingly, the tumors developed in our *PPM1D* transgenic mice are phenotypical similar to mice with *Trp53* mutations compared to *Trp53* knockout mice. Mice with *Trp53* deletions mainly develop lymphomas and less frequently sarcomas^38,40^ whereas *Trp53* mutant mice, in addition to lymphomas and more frequently sarcoma, also develop carcinomas^39^. This is similar to the spectrum of tumors observed in our *PPM1D* transgenic mice. Thymic lymphoma derived from *PPM1D* transgenic mice expresses high levels of wild-type p53 and WIP1 similar to *Trp53* mutated mice showing high levels of mutated p53^85^.

Also, both the genomic landscape and expression profiles obtained from DNA and RNA sequencing of *PPM1D*-induced tumors had similar features with regard to DNA mutations and expression profiles as observed in tumors deriving from p53 impaired mice. Hence, *PPM1D* is an oncogene that induces tumor development by repressing p53 activity leading to deviant cell cycle arrest, DNA repair and apoptosis.

Taken together, we have dissected the functional role of *PPM1D* in the neural childhood tumors neuroblastoma and demonstrated that *PPM1D* is able to induce tumor growth when cells are subjected to DNA-damaging stress. Hence, *PPM1D* is a *bona fide* oncogene. We also demonstrate that WIP1 constitute a druggable target in neuroblastoma as well as in medulloblastoma that should be further developed and evaluated in combinations with current treatment modalities and investigated for testing in clinical trials given the fact that the majority of patients with poor prognosis have aberrant expression of WIP1. It may be argued that targeting DNA repair mechanisms and phosphatase activity in particular seems a problematic hurdle, but our current data are promising proofs of a principle providing molecular and pharmacological evidence. Furthermore, genetic instability and accumulation of genetic aberrations over time is a major obstacle in metastatic and relapsing pediatric cancers further supporting the potential role and impact of *PPM1D* as a promising therapeutic target for pediatric patients with high-risk neuroblastoma and medulloblastoma as well as a wide range of adult cancer patients.

## ACKNOWLEDGEMENTS

We wish to thank Karolinska Center for Transgene Technologies (KCTT) for the transgenic production service. William A. Weiss and Louis Chesler for providing the construct vectors used for generating transgenic *PPM1D*-mice. Clinical Genomics, SciLifeLab, Gothenburg, Sweden and the Bioinformatics Core Facility platforms at the Sahlgrenska Academy, University of Gothenburg, Gothenburg, Sweden for assistance with the bioinformatical analysis of sequencing data. We thank Hjalmar Ståhlberg-Nordegren, Teodora Andonova, Catarina Träger, Inger Bodin and Susanne Ahlberg for their help and contribution to this work. Special thanks to Jessica Lundgren and Selameyhune Assefa for their excellent assistance with the *PPM1D*-transgenic animals. We convey our gratitude to Drs. E Appella and SJ Mazur for useful critical assessment of our manuscript.

This work was supported with grants from the Swedish Childhood Cancer Foundation, the Swedish Research Council, the Swedish Cancer Foundation, the Swedish Foundation for Strategic Research (www.nnbcr.se), Karolinska Institutet, Märta and Gunnar V Philipson Foundation, and The Cancer Research Foundations of Radiumhemmet.

The study sponsors had no role in the design of the study; the collection, analysis, and interpretation of the data; the writing of the manuscript; or the decision to submit the manuscript for publication.

## Author contributions

Conceptualization, J.M., J.I.J., and P.K.; Investigation and validation, J.M., S.F., M.G., G.G.O., D.T., M.W., B.S., M.W., L.H.M.E., N.E., A.K., J.Y.H., A.M., G.T.; Computational investigation and analysis, J.M., S.F., T.K.O., F.H., C.B., S.R., F.A., N.J., S. K., Y.S., C.K., J.H., D.G., M.J., M.K., T.M., J.I.J and P.K.; Writing – Original draft, J.M, N.B., J.I.J. and P.K.; Writing – Review & Editing, J.M., S.F., C.K., J.J.M., M.F., M.K., N.B., J.I.J. and P.K.; Resources, K.T., K.S., TM, MF, M.K., J.J.M., G.T., M.J., T.M. and P.K.; Final editing and manuscript approval, All authors; Funding Acquisition & Supervision, J.I.J. and P.K.

## STAR METHODS

### Patient-derived tumor material and clinical data

The collection of neuroblastoma (NB) and medulloblastoma (MB) tumors from Swedish patients was performed after either written or verbal consent was obtained from parents/guardians according to ethical permits approved by the local Ethics Committee (Karolinska Institutet and Karolinska University Hospital, registration number 03-736, and 2009/1369-31/1(NB), 2009/1608-31/4(MB)). Fresh tumor tissue was obtained at surgery and snap-frozen and stored at -80°C until analysis. Informed consent for using tumor samples in scientific research was provided by parents/guardians of all minors (<18 years of age). In accordance with the approval from the Ethics Committee the informed consent was either written or verbal. Clinical data on all patients were obtained from hospital records and/or the National Solid Tumor Registry (03-736, 03-642, 2009/1608-31/4 and 2009/1369). All patients were diagnosed, managed and treated according to national and international guidelines and protocols.

### Neuroblastoma patient genomics and transcriptomics

#### Patient material for genetic analysis

DNA was extracted from frozen tumors or blood using DNeasy blood and tissue kit (Qiagen, Hilden, Germany) according to manufacturer’s protocol and evaluated through absorbance measurements, fluorometric quantitation and DNA integrity assessment on Agilent Tapestation (Agilent, Santa Clara, CA).

#### SNP-microarray analysis

Microarray analysis was performed on a consecutive series of samples from an unselected cohort of 271 neuroblastoma patients using Affymetrix gene mapping arrays (Affymetrix Inc., Santa Clara, CA). These analyses include the majority of cases from Sweden. All tumors were staged according to the International Neuroblastoma Staging System (INSS) and INRG criteria. Handling of the microarrays has been described previously^6,86^.

Primary data analysis was performed using GDAS software (Affymetrix) with *in silico* normalization against control samples from healthy individuals with further analyses performed using CNAG (Copy number Analyzer for Affymetrix GeneChip Mapping Arrays) (Genome Laboratory, Tokyo University; http://www.genome.umim.jp). All cases of chromosomal gain, loss, or amplification were scored for both segmental and numerical aberrations, including detailed information about the breakpoint positions when applicable. For practical reasons, all notations of gain and loss were considered in relation to a nominal tumor karyotype and a chromosomal break in a tumor was defined as a clear change in gene dose level in the genomic profile. All genomic position annotations are stated based on the hg19 build (http://genome.ucsc.edu/) of the human genome.

#### Analysis of PPM1D/17q clonal evolution

Single nucleotide polymorphism (SNP) arrays including the Oncoscan (formalin fixed paraffins embedded samples) and Cytoscan HD (fresh frozen samples) platforms (Affymetrix Inc.) were retrieved from and subjected to comparisons of 17q copy number including the *PPM1D* region (pos 17:60,600,183-60,666,280; GRCh38.p12). Evolutionary trajectories were formalized as phylogenetic ideograms along the lines described in the original publication. In all, 100 samples from 23 neuroblastomas from 23 patients were analyzed. Each patient was assigned to one of three simplified clinical-genetic risk groups, including children <18 months with only numerical aberrations found in their tumor’s phyologenetic stem by SNP array (low-risk patients), tumors with *MYCN* amplification in the stem (high-risk patients), and tumors with structural rearrangements in the stem (high-risk patients).

#### Genome sequencing

In total, tumor material from 73 Swedish neuroblastoma patients was subjected to either whole genome and/or whole exome sequencing. Whole exome sequencing was performed through paired-end sequencing on Illumina platforms (Illumina, San Diego, CA) after enrichment with Agilent SureSelect All Exome enrichment kit (Agilent technologies, Santa Clara, CA) on DNA for 18 tumor/normal pairs and additional 20 neuroblastoma tumors without corresponding constitutional DNA. Alignment against hg19 was performed using BWA with GATK local realignment followed by SNV calling using SNPeff. Whole genome sequencing (WGS) was performed on tumor DNA and corresponding constitutional DNA extracted from blood for 35 neuroblastoma patients for an average coverage of at least 60X for tumor and 30X for constitutional DNA using Illumina xTen instrumentation (Illumina, San Diego, CA, USA) located at NGI/Clinical Genomics, SciLife Laboratories, Stockholm, Sweden. Read trimming, mapping to the human reference genome hg19 and variant calling were performed using CLC Genomics Workbench 8.0.3 software (CLC, Aahus, Denmark). Only high quality called variants with a minimum 10% allele frequency and a total read coverage of ten were considered for further analysis. Somatic variants with allele frequency above 3% in either 1000 genomes, Exome Aggregation Consortium (ExAC), Cambridge, MA (http://exac.broadinstitute.org) or NHLBI Exome Sequencing Project (http://evs.gs.washington.edu/EVS/) were discarded as well as excluding all synonymous variants or variants in non-coding regions except those affecting canonical splice sites. Remaining variants were assessed manually through the Integrative Genomics Viewer (IGV)^87^ for removal of calls due to mapping artifacts and paralogs.

For tumors (exome) sequenced without corresponding DNA from constitutional tissue, a systematic filtering approach was used to identify critical variants. This was done by removal of common variants, e.g. present in dbSNP v138 or showing an allele frequency above 0.1 in either 1000 genomes, Exome Aggregation Consortium (ExAC), Cambridge, MA (http://exac.broadinstitute.org), NHLBI Exome Sequencing Project (http://evs.gs.washington.edu/EVS/) or SweFreq (https://swegen-exac.nbis.se/). Variants were annotated based on RefSeq and functional impact was predicted by SIFT and PolyPhen. FusionCatcher was applied to discover *PPM1D*-containing fusion transcripts in the National Cancer Institute TARGET dataset comprising paired-end RNA sequenced neuroblastoma patient samples (dbGap Study Accession:phs000218.v16.p6).

#### Case description, germline PPM1D mutation detection

The patient was born by a spontaneous delivery at gestational week 37+4 after an uneventful pregnancy. His birth weight was 2945 gr. He was diagnosed with a bicuspid aortic valve, which was also present in the maternal grandmother and two of her siblings. The family history was negative for cancer with onset before age 50. In the first year of life he had reflux for which feed thickener and proton pomp inhibitors were described. Psychomotor development was above average with early speech development, early recognition of numbers, letters and colors and walking shortly after his first birthday. At 26 months of age he presented with hemifacial paresis and clinical, radiological and pathological evaluation revealed a metastatic stage 4 (INSS) intermixed ganglioneuroblastoma of the left adrenal gland with bone metastases in the skull base, in the lumbosacral spine with intraspinal extension and expansion to the left-sided ilium and diffuse bone marrow infiltration. The tumor was negative for *MYCN* amplification and negative for LOH of chromosome 1p. Therapy consisted of induction chemotherapy, surgical resection, high dose chemotherapy, autologous stem cell transplantation and radiotherapy which resulted in very good partial clinical remission. Treatment was intended to be completed with immune therapy but evaluation before starting this treatment revealed refractory neuroblastoma in the ilac bone, which was irradiated and again three cycles of chemotherapy (irinotecan/temozolomide) were given. The evaluation that followed showed diffuse leptomeningeal metastases indicative of progressive disease. Palliative treatment was started and the boy died at 3 years and 7 months of age. During treatment it was noted that the boy had a remarkably high pain threshold. Shortly after his cancer diagnosis the parents worried about a possible increased risk for neuroblastoma in the younger brother of the proband and a clinical geneticist was consulted. Fetal fingers pads and a relatively short stature (at cancer diagnosis his age-adjusted length and head circumference were at -1.5 SD) were observed in the proband, but otherwise no dysmorphisms or other notable features were present. Germline *PHOX2B* and *ALK* analysis revealed no mutations. Several years later, the brother developed subcutaneous lesions on his right forearm at three years of age. These were resected and were hard to classify by the pathologist, with benign myofibroblastic proliferation as most likely diagnosis. The parents again consulted a clinical geneticist, in the light of a possible genetic predisposition explaining the tumors in their children. Whole exome sequencing was performed on blood derived DNA of both children and the parents. This revealed a heterozygous c.1528C>T (p.Gln510*) mutation in *PPM1D* in the proband who died of neuroblastoma. This mutation was absent in his brother and his parents. Identification of this mutation in neuroblastoma derived DNA confirmed the germline status of this mutation and excluded that the mutation was restricted to blood and resulted from chemotherapy which has recently been shown for *PPM1D* mutations^88^.

The patient described here has several features in overlap with patients with *PPM1D* intellectual disability syndrome. He had relative short stature, reflux in early childhood, small hands (hand length -2,5 SD at age 9 months), and a high pain threshold. Strikingly, developmental delay was definitely absent in this child. Despite intensive multimodal treatment for his high-risk neuroblastoma, he seemed to have an above average intelligence, which was in concordance with the education levels of his parents and grand parents. Additionally, deep phenotyping revealed overlapping behavioral problems (ASD, ADHD, and anxiety disorders), hypotonia, broad-based gait, facial dysmorphisms, and periods of fever and vomiting. This patient, with the first de novo germ-line *PPM1D* mutation linked to cancer, showed a facial phenotype that significantly correlated (p=0.0164, **Supp Figure S1F**) with eleven previous reported patients with de novo germ-line truncating *PPM1D* mutations of the 5^th^ or 6^th^ exon and intellectual disability^24^.

#### Gene expression analysis

Patient cohort and methodology of RNA-seq data analysis pipeline are extensively described in^89^. We used gene expression data of 498 neuroblastoma patients from seven countries. According to the Neuroblastoma Risk Group (INRG) classification system^26^ we classified patients with *MYCN* amplifications and patients with a metastatic disease and older than 18 months at diagnosis as high-risk patients (n=176). Events were defined according to a revised version of the International Neuroblastoma Response Criteria^90^.

RNA-seq gene expression analyses have been done using the MAGIC-AceView pipeline as previously described^89^. Copy number data for *PPM1D* was derived from array comparative hybridization (aCGH);^91^, whole exome sequencing and whole genome sequencing analyses^74^.

#### Statistical analyses

Gene expression of *PPM1D* was investigated displayed on the gene level (*PPM1D*) or transcript level for two selected transcript variants (AceView annotation, as a result of the analysis, i.e. *PPM1D*.*aAug10* and *PPM1D*.*bAug10*, corresponding to transcripts ENST00000305921 and ENST00000392995 respectively).

*PPM1D* expression in defined NB subgroups (according to INRG high-risk status, INSS stage, amplification status of MYCN, age, CN status of *PPM1D* on chromosome 17) was displayed in Tukey box-and-whisker plots (box range: inter-quartile range, whiskers: 1.5x IQR). Where appropriate, either Mann–Whitney–Wilcoxon or Kruskal-Wallis tests were performed to investigate differences in expression of two or more analyzed groups, respectively. Pearson correlation analyses have been applied to investigate correlation between *PPM1D* expression and copy number.

To investigate the prognostic value of *PPM1D* expression, optimal cutpoint expression values separating the cohort into *PPM1D*-high and *PPM1D*-low groups that correlate best with outcome in terms of event-free survival (EFS; recurrence, progression and death from disease were considered as events), and overall-survival (OS) were calculated by means of maximally selected rank statistics (maxstat package in R v3.4.3)^92,93^ in training sets. Differences in survival of *PPM1D* high and low-expressing subgroups according to dichotomization with the established cutpoints was then tested by Mantel-Cox (log-rank) tests in corresponding validation sets. For this procedure, equally sized training and validation subsets (n=296) were randomly sampled from the entire cohort (n=498). Resampling was done 1000 times and the mode of those established cutpoint values which passed p < 0.05 in maxstat^92,93^ as well as in the log-rank test was eventually selected to define *PPM1D*-high and *PPM1D*-low expressing subgroups (this procedure was independently done for EFS and OS). Subsequently, Kaplan-Meier survival estimates for the entire cohort were calculated and displayed from the time of diagnosis and clinical outcome of respective subgroups was compared using the log-rank test.

#### Principal Components Analysis (PCA) on expression data

Raw data expression: CEL-files (HU133A chips from Affymetrix) from 30 primary neuroblastoma cases from two published microarray studies^28,29,94^ were pre-processed using gcRMA normalization in Bioconducter for R 2.9.2 (library BioC 2.4). For each gene, the mean expression level from probe-sets was calculated, resulting in 8105 genes/variables^94^. Principal Components Analysis (PCA) was performed on the pre-processed McArdle/Wilzén expression data set using Omics Explorer 3.2 from Qlucore (www.qlucore.se). Genes with the lowest variance were filtered out (variance cut-off =0.4), displaying a PCA plot based on 716 genes/variables and 30 samples/cases according to the method previously described^94^. Samples/cases were joined to their nearest neighbor using Euclidean distances, and they subdivided into four separate groups. The differential expression of *PPM1D* between molecular subgroups was calculated by fold change and change and Student’s t-test (2-sided comparison, unequal variance).

### Gene expression data profiling for medulloblastoma

All mRNA expression data of 446 human medulloblastoma and 18 normal cerebellum samples have been generated by Affymetrix U133plus2.0 arrays. Gene expression data come from public sources deposited in GEO (http://www.ncbi.nlm.nih.gov/geo)^95-99^ or have been generated at the German Cancer Research Center DKFZ in Heidelberg. The MAS5.0 algorithm of the GCOS program (Affymetrix Inc) was used for normalization of the expression data. All data have been analyzed using the R2 program for analysis and visualization of microarray data (http://R2.amc.nl). Anova tests were used to test whether there is a significant differential expression of WIP1 between the groups shown in the plots.

### Cell culture and reagents

Twenty-two human cell lines of different origin were used throughout the study: eleven neuroblastoma cell lines (IMR-32, Kelly, NB1691, SH-EP, SH-SY5Y, SK-N-AS, SK-N-BE(2), SK-N-DZ, SK-N-FI, SK-N-SH, TR14), eight medulloblastoma/sPNET cell lines (DAOY, D283MED, D384MED, D425MED, D458MED, MEB-MED8A, PFSK-1, UW228-3), two breast cancer cell lines (MCF-7, BT-474), and one human fetal lung fibroblast cell line (MRC-5). In addition, one neural multipotent progenitor cell line from mouse (C17.2) was used. The cell lines were purchased from ATCC, except D384MED, D425MED, D458MED, PFSK-1, MEB-MED8A and UW228-3 that were kindly provided by Dr. M. Nistér (Karolinska Institutet), NB1691 and TR14 by Dr. D. Tweddle (Newcastle University) and C17.2 by Dr. T. Ringstedt (Karolinska Institutet). The cell lines were cultured in RPMI 1640 (IMR-32, Kelly, NB1691, SH-EP, SK-N-AS, SK-N-BE(2), SK-N-DZ, SK-N-FI, SK-N-SH, TR14, PFSK-1 and MRC-5), Dulbecco’s modified Eagle’s medium (DMEM; MEB-MED8A, C17.2, BT-474), Minimum Essential Media (MEM; DAOY, D283MED, D384MED), Richter’s improved MEM with zinc/DMEM (IMEMZO/DMEM; D425MED and D458MED), DMEM/F12 (SH-SY5Y and UW228-3). Medium was supplemented with 10% (or 15% for C17.2, MEB-MED8A, D425MED and 20% for D384MED) heat-inactivated fetal bovine serum (FBS), 2 mM L-glutamine, 100 IU/ml penicillin G, and 100 μg/ml streptomycin (Life Technologies Inc., Stockholm, Sweden) at 37°C in a humidified 5% CO_2_ atmosphere. To the MCF-7 and D384MED media, 1 mM sodium pyruvate and 1 mM non-essential amino acids solution (Gibco) were also added. All media were purchased from Gibco BRL.

The identities of the cell lines were verified by short tandem repeat genetic profiling using the AmpFlSTR® IdentifilerTM PCR Amplification Kit (Applied Biosystems) in December 2015 and all cell lines were used in passages below 25. All experiments were executed in Opti-MEM (GIBCO) supplemented with glutamine, streptomycin and penicillin (HyClone Thermo Fisher Scientific), except transfection experiments, which were performed without antibiotics. *PPM1D*-knockdown SK-N-BE(2) cells and corresponding control cells were cultured in selection media (standard media according to above supplemented with 0,5-2 µg/mL puromycin).

RITA and Nutlin-3 were purchased from Cayman Chemical Company and Sigma-Aldrich, respectively, and SL-176 and SPI-001 were synthesized as described previously^42,44^. RITA and Nutlin-3 were dissolved in DMSO (Sigma-Aldrich), while SL-176 was dissolved in a mix of DMSO (33%) and ethanol (67%). Further dilutions were made in Opti-MEM or PBS. The DMSO concentration did not exceed 1% v/v in any experiment. For the *in vivo* studies, SL-176 was dissolved in a mix of DMSO (33%) and ethanol (67%) and further diluted in sodium chloride 0.9%.

### Short hairpin RNA (shRNA)

For the transfections, cells were seeded in 6-well plates, left to attach and transfected using Lipofectamine 2000 (Thermo Fisher Scientific) with 4 µg of four pre-designed shRNAs (GIPZ Lentiviral) targeting human *PPM1D* (172_0502-F-1 (Clone ID: V2LHS_262759), 172_0556-D-7 (Clone ID: V2LHS_27794), 172-0447-C-2 (Clone ID: V2LHS_27798) and 172_0496-G-2 (Clone ID: V2LHS_262763), Dharmacon) and non-silencing pGIPZ Lenti Control shRNA (#RHS4346, Dharmacon). Cells were incubated for 6 hours in the transfection media and then replaced with corresponding culture medium. After 24-48 hours, cells were subjected to further analyses. For generating stable transfections, cells were grown in selective medium (0,5-2 µg/mL puromycin selection).

### Viability assays

For evaluation of cytotoxic effect on cell viability, we used the fluorometric microculture cytotoxicity assay as previously described^100^ or the colorimetric formazan-based assay WST-1 (Roche), according to the manufacturer’s description. Briefly, cells were seeded into 96 well plates (5 000 – 10 000 cells/well), left over night and treated with drugs the following day. After 72 hours, WST-1 reagent was added and absorbance was measured at 450 nm. All concentrations were tested in triplicate. The mean out of at least three independent experiments is reported.

To determine colony formation, 100 cells/well in the non-exposure experiments and 300 cell/well in the irradiation experiments were seeded in 60 mm cell+ culture plates (Sarstedt, Sweden) and allowed to attach for 24 h before exposure to ionizing radiation (Cobolt^60^ source) at 2 or 4 Gy, when applicable. After 10-14 days of incubation in medium, cells were washed, fixed in formaldehyde (4%), stained with Giemsa (Gibco, BRL) and colonies (1 clone>50 cells) with 50% plate efficiency were manually counted. The mean out of at least three experiments is reported.

Cell viability of *PPM1D* silencing was assessed by the trypan blue exclusion assay. In brief, cells (4.4 × 10^4^ MEB-MED8A cells/well and 2.5 × 10^4^ SK-N-BE(2) cells/well respectively), were seeded and transfected in 6-well plates; 3 wells for each time point per cell line and transfection group, cultured for 6 days. Cells were stained with 0.4% trypan blue (GIBCO, BRL) and viable (unstained) cells were counted daily to determine the total number of living cells. The mean out of three experiments is reported.

### Irradiation of human cancer cell lines

SK-N-BE(2), SH-SY5Y, DAOY, Med8a and MCF-7 cells were seeded into six-well cell culture plates (300 000 – 500 000 cells/well) in standard medium with 10% FBS and allowed to attach overnight, with exception for Med8a cells growing in suspension. Prior to treatment, 60-80% confluency was observed. Medium was removed and replaced with OptiMEM containing a SL-176 concentration equivalent to the corresponding IC_50_ value for each cell line (0.5 - 1.3 µM). Med8a cells were directly seeded in OptiMEM containing SL-176 at IC_50_. After 1h incubation, cells were irradiated with 4 Gy (Cobolt^60^ source) while kept on ice, after which incubation at 37°C continued for the time indicated (0, 4, 8, or 24 hours). Cells were harvested using Cell Dissociation Solution Non-enzymatic (Sigma-Aldrich). For the irradiation of the stably transfected SK-N-BE(2) neuroblastoma cell lines (*PPM1D* ShRNA and control ShRNA), 500 000 cells/well were seeded into six-well cell culture plates as described above, allowed to attach overnight and irradiated with 2 Gy and 4 Gy respectively and harvested after 48 and 72 hours. SK-N-BE(2) non-transfected cells were treated with 40mM cisplatin, used as a positive control for double-strand DNA breaks.

### Western blot

Protein extraction, determination of protein content, SDS-PAGE under reducing conditions, electroblot and immunoreaction detection were carried out as previously described^101^. Briefly, total proteins were extracted using RIPA buffer (Thermo Fisher Scientific) supplemented with Halt™ Protease and Phosphatase Inhibitor Cocktail (Thermo Fisher Scientific). 30 μg (for cell lines) and 50 ug (for mouse tissue) of total protein were resolved by reducing SDS-PAGE and transferred onto a nitrocellulose membrane. The membranes were blocked with 5% non-fat milk or BSA 5% in TBS-T buffer, and incubated with the corresponding antibodies (**Table S7**). Chemiluminescence visualization of antibodies was performed with Amersham™ ECL™ Prime Western Blotting Detection Reagent (GE Healthcare). Visualization and imaging of signal was performed with ImageQuant LAS 4000 (GE Helathcare).

### Quantitative real-time RT-PCR analyses

The mRNA expression levels were quantified using TaqMan® technology on an ABI PRISM 7500 sequence detection systems (PE Applied Biosystems). Primers were selected from the Assay-on-Demand products (Applied Biosystems), including human *PPM1D* (Hs00186230_m1), and 18S ribosomal RNA (Hs99999901_s1). All gene expression assays was designed with an FAM reporter dye at the 5’ end of the TaqMan MGB probe, and a non-fluorescent quencher at the 3’ end of the probe. High capacity RNA-to-cDNA kit (Applied Biosystems) was used to synthesize cDNA from 100 ng of RNA per sample. The PCR reaction was done in a total reaction volume of 25 µl containing 1 x TaqMan® Universal PCR Master Mix, 1 x TaqMan® Gene Expression Assays (Applied Biosystems) and 10 µl of cDNA from each sample as a template, in MicroAmp optical 96-well plates covered with MicroAmp optical caps (Applied Biosystems). Firstly, samples were heated for 2 min at 50°C and then amplified for 40 cycles of 15 s at 95°C and 1 min at 60°C. A standard curve was generated for relative quantification with cDNA synthesized from 1 μg RNA of the cell lines combined. For every sample the amount of target mRNA was normalized to the standard curve and normalized to 18S ribosomal RNA expression. All experiments included a no template control and were performed in triplicate.

### CRISPR-Cas9 loss of function screening

The DepMap Public CRISPR (Avana) 18Q3 gene dependency dataset including 485 cancer cell lines (whereof 15 neuroblastom and 7 medulloblastoma cell lines) as well as mutation call dataset was downloaded from the Broad Institute Cancer Dependency Map (https://depmap.org/portal/) and used for analysis of *PPM1D* and *TP53* genetic vulnerabilities^102^. Visualization and analysis of enriched functional processes associated to *TP53*-dependency was done in the Search Tool for the Retrieval of Interacting Genes/Protein (STRING) database^103^.

### Flow cytometry

Phosphorylation of H2AX was assayed with Alexa Fluor 647-conjugated anti-phospho-H2AX (2F3, Biolegend, San Diego, CA, USA) 24 and 72 hours on cells that were transfected with *PPM1D-*shRNA 172_0502-F-1 and control shRNA, respectively. A minimum of 10 000 events were recorded on Becton-Dickinson FACSCalibur or LSR II flow cytometers (BD Biosciences, San Jose, CA, USA). Data analysis was performed using the Cell Quest software.

### Human tissue samples for immunohistochemistry

Neuroblastoma and medulloblastoma tumor tissue were obtained from the Karolinska University Hospital according to the ethical approval from the Stockholm Regional Ethical Review Board and the Karolinska University Hospital Research Ethics Committee (approval ID 2009/1369-31/1 and 03-736). Informed consent (written or verbal) was provided by the parents or guardians for the use of tumor samples in research. Samples were collected during surgery, snap-frozen in liquid nitrogen and stored at -80°C until further use. Twenty-seven neuroblastoma samples derived from children of different ages and all clinical stages, including different biological subsets^104^, were analyzed.

### Immunohistochemistry

Formalin-fixed and paraffin-embedded human tissue sections were deparaffinized in xylene and graded alcohols, hydrated and washed in phosphate-buffered saline (PBS).

After antigen retrieval in sodium citrate buffer (pH 6) in a microwave oven, the endogenous peroxidase was blocked by 0.3 % H2O2 for 15 min. Sections from human neuroblastoma were incubated overnight at 4 °C with a primary antibody against PPM1D (ab31270, Abcam). Similarly, tissue sections from xenograft tumors were incubated with an anti-Ki-67 antibody (clone SP6, Neomarkers, Fremont), anti-cleaved caspase-3 (ASP175) (#9579; Cell Signaling Technology) and anti phospho-Histone h2ax (Ser139) (#9718 Cell Signaling Technology), respectively. As a secondary antibody, the anti-rabbit-horseradish peroxidase (HRP) SignalStain Boost IHC detection kit was used (Cell Signal Technology; #8114). A matched isotype control was used as a control for nonspecific background staining (not shown).

### Animal studies

#### Ethical permits

The animal experiments were approved by the regional ethics committee for animal research in Northern Stockholm, appointed and under the control of the Swedish Board of Agriculture and the Swedish Court. All animal experiments were in accordance with national regulations (SFS 1988:534, SFS 1988:539, and SFS 1988:541). For specific approval numbers, please refer to the sections below.

#### Xenograft studies

Immunodeficient nude mice (female 4-6 weeks old, NMRI-nu/nu, Scanbur, Stockholm, Sweden) were used for xenograft studies (ethical approvals N304/08 and N391/11). The mice were kept under specific pathogen-free conditions at a maximum of six individuals per cage and given sterile water and food *ad libitum*. All mice were treatment-naïve at the start of the experiment.

Under general anesthesia, each mouse was injected subcutaneously on the rear flank with 10 × 10^6^ SK-N-BE(2) neuroblastoma cells or 17 × 10^6^ DAOY medulloblastoma cells. In the knockdown experiment, mice were inoculated bilaterally with 5 × 10^6^ SK-N-BE(2) cells (clone d) that were knocked down for *PPM1D* (n=8) and SK-N-BE(2) control cells (clone C) that were transfected with non-silencing shRNA (n=15) respectively and followed until the tumor reached 0.1 mL.

In the drug treatment experiments, mice with SK-N-BE(2) xenografts were randomly assigned to three different treatment groups when the tumor reached ≥0.1 mL. For twelve days, mice received either daily intraperitoneal injections (i. p.) of SL-176 at 3 mg/kg (n=8) or 0.5 mg/kg (n=6), or no treatment (n=6). The mean tumor volume at the start of treatment was 0.115 mL.

Mice bearing DAOY xenografts were randomly divided into two different groups, and treatment commenced at tumor volume ≥0.12 mL. Mice received either 3 mg/kg SL-176 as daily i. p. injection for 21 days (n=8), or no treatment (n=7). The mean tumor volume at the start of treatment was 0.124 mL.

In all xenograft-bearing mice, tumors were measured every day and the animals were monitored for signs of toxicity including weight loss. The tumor volume was estimated as (width)^2^ x length x 0.44. At sacrifice, tumors were dissected and either frozen or fixed in formaldehyde, for subsequent analyses.

#### WIP1-overexpressing mouse model

Transgenic WIP1-overexpressing mice were generated by pronuclear injection with random integration of a *PPM1D-*transgenic vector construct, carried out at Karolinska Center for Transgene Technologies (KCTT) and granted according to ethical approval numbers N251-12 and N42-14.

The construct consisted of rat tyrosine hydroxylase (TH) promoter, meant to direct expression towards the neural crest; rabbit beta-globin intron was used to enhance expression; and cDNA for the human *PPM1D* gene. Herpes simplex virus (HSV) thymidine kinase gene sequence was used as a transcription terminator (**Figure S4A**). The plasmid was kindly provided by Prof. William Weiss (University of California, San Francisco, California, USA) and has previously been used to generate a transgenic mouse model with targeted *MYCN* expression giving rise to murine neuroblastoma^67^. In the current study, however, *MYCN* cDNA was substituted for *PPM1D* cDNA (MGC Human *PPM1D* Sequence-Verified cDNA, Clone Id: 5167004, Dharmacon) to complete the construct (GENEWIZ, South Plainfield, NJ, USA).

The linearized and purified construct was diluted to 1.5 ng/ul in microinjection buffer (10 mM Tris-HCl, pH7.4, 0.1 mM EDTA) and injected into the pronucleus of C57BL/6NCrl zygotes using a Nikon TE200 microinjection system with Narishige NT-88NEN micromanipulators and a Warner Instruments PLI-100A pico-liter injector. The microinjected embryos were transferred into the oviducts of pseudopregnant Crl:CD1(ICR) female mice using standard surgery techniques and ear biopsies from the resulting 55 offspring were screened for the presence of the *PPM1D* transgene by PCR using forward primer: 5’-CTGGTCATCATCCTGCCTTTCT-3’ and reverse primer: 5’-GCCTTTCCCCGAGACTTCG-3’ (Sigma-Aldrich). 6 transgenic animals were found of which 4 founder animals were further established based on their ability to pass on the human *PPM1D*-transgene to their offspring.

In order to achieve a more tumor-permissive genetic background, these four transgenic mouse-lines were backcrossed with 129×1/SvJ mice, aiming for ten generations of backcrossing which should result in approximately 100% 129×1/SvJ background.

Backcrossing was carried out in accordance with ethical permit number N641-12. Throughout breeding, mice were monitored closely for development of palpable abdominal tumors or other disease manifestations up to 548 days (1.5 years) of age.

#### Irradiation of mice

Using a linear accelerator (X-RAD 320, Biological Irradiator, North Branford, CT, USA) with a dose rate of 0.95 Gy/min at 320 KV and a radiation field of 20 × 20 cm, *PPM1D*-transgenic mice from three established transgenic lines and their wild-type littermates resulting from heterozygous breeding pairs with different degrees of 129×1/SvJ strain background (3 to 7 generations of backcrossing from C57BL/6N to 129×1/SvJ background) were subjected to whole-body irradiation at a single sublethal dose of 4.5 Gy (ethical approval N290-15), after which they were monitored daily in their usual pathogen-free environment. Mice were exposed to irradiation at different ages (1 to 314 days old). Littermates were always irradiated simultaneously. The mice were followed up to 548 days (1.5 years) of age.

#### Immunohistochemistry of transgenic mouse tissue samples

Formalin-fixed and paraffin-embedded transgenic mouse tissue sections were deparaffinized in xylene and graded alcohols, hydrated and washed in phosphate-buffered saline (PBS). After antigen retrieval in sodium citrate buffer (pH 6) in a microwave oven, endogenous peroxidase activity blocked by 0.3 % H_2_O_2_ for 15 min. Biotin blocking was preformed using an avidin/biotin blocking kit (Vector Laboratories). All washes and dilutions were performed in PBS containing 0.1% saponin. Sections were incubated with an anti-Ki-67 antibody (clone SP6, ab16668, abcam), anti-B220 antibody (clone RA3-6B2, R&D Systems, Minneapolis, MN, USA), or anti-CD3 antibody (SP7, Abcam, Cambridge, UK) containing 3% human serum overnight at room temperature. After blocking with 1% goat serum for 15 min sections were incubated with a biotin-conjugated secondary antibody (goat anti-rabbit IgG or goat anti rat IgG, Vector laboratories) containing 1%goat and 3% human serum at room temperature for 30 min. For detection an ABC complex (Elite ABC kit, Vector laboratories) was used before the sections were developed using diaminobenzidine (DAB Peroxidase Substrate kit; Vector Laboratories) as chromogen. Sections were counterstained with Mayer’s hematoxylin (Histolab). For immunofluorescence detection of CD3 positive cells sections were incubated with anti-CD3 antibody (SP7, Abcam, Cambridge, UK) in 4% goat serum overnight at +4C following incubation with secondary Goat anti-rabbit-Alexa Fluor 488 (Thermo fisher Scientific). Histological assessment of the transgenic mouse tissues was performed by a pathologist. For full list of antibodies used for immunohistological analyses see **Table S7**.

#### Whole exome sequencing, variant calling and copy number alterations in mice

Whole exome sequencing (WES) was performed on DNA from totally 24 samples; 15 independently developed thymic lymphomas and thymus from nine controls. The controls were dived in to three different groups corresponding to a) irradiated *PPM1D*-transgenes, b) irradiated wild type and c) non-irradiated wild type. In respective control group three mice, one from each strain, were subjects for sequencing.

WES was performed by Otogenetics (Otogenetics Corporation, Atlanta, GA, USA) through pair-end sequencing on Illumina platforms (Illumina, San Diego, CA) after enrichment with Agilent SureSelectXT Mouse All Exon (Agilent technologies, Santa Clara, CA) reaching an average coverage of 80X (range 47,5-99,2X) (**Table S5**). Read trimming, mapping to the mouse reference genome mm10 and variant calling were performed using CLC Genomics Workbench 5.0 software (CLC, Aahus, Denmark). Somatic calling of lymphoblastic tumors and irradiated controls was done using the combined sequence from the three non-irradiated wild type mice as normal control. Only high quality called variants with a minimum 10% allele frequency and a total read coverage of ten were considered for further analysis. All synonymous variants or variants in non-coding regions except those affecting canonical splice sites were discarded. Remaining variants were assessed manually through the Integrative Genomics Viewer (IGV)^87^ for removal of calls due to mapping artifacts or paralogs.

Calling and visualization of copy number alterations was done using the software Control-Free (control-FREE Copy Number Caller) that generates rations from normalized read distribution between tumor and normal followed by visualization in a Shiny application as described previously^105^.

#### RNA sequencing of mouse tumors

Bulk RNA sequencing (RNA-seq) was performed on extracted RNA in a total of 24 samples; 10 controls (thymic tissue from healthy mice) and 14 thymic lymphomas. As with mouse tumor WES, controls were divided into three groups based on irradiation status and genotype: wild-type non-irradiated (n = 3), *PPM1D-*transgenic non-irradiated (n = 3), wild-type irradiated (n = 2), *PPM1D-*transgenic irradiated (n = 2). Total RNA extraction, library preparation and sequencing were performed by Otogenetics (Otogenetics Corporation, Atlanta, GA, USA) through paired-end sequencing on Illumina HiSeq 2500 (Illumina, San Diego, CA) after mRNA purification using the TruSeq Stranded cDNA kit. Read length was 100-125 bp for all samples.

Samples were aligned to the mm10 reference genome using STAR in 2-pass mode^106^. Aligned reads were quantified using htseq-count^107^ and differential gene expression analysis was performed using the R/Bioconductor package DESeq2^108^. After having converted mouse Ensembl gene IDs to human orthologs using the R package *gOrth*, gene set enrichment analysis was performed using the GSEA software from Broad institute^109,110^ and the Hallmark collection of gene sets from the Molecular Signatures Database, MSigDB, version 6.2^111^.

### Statistical analysis

Statistical analyses were done with GraphPad Prism software (GraphPad Software, San Diego, CA). The IC_50_ values (inhibitory concentration 50%) were determined from log concentrations-effect curves using non-linear regression analysis. T test was used to compare means between two groups and for comparison of three or more groups, one-way ANOVA followed by Bonferroni multiple-comparisons post-test were used. Survival analysis was examined with log-rank test and Fisher’s test was used to test significance of association between the two categories. Correlations were assessed with Pearson test/Spearman non-parametric test. P<0.05 was considered significant and all tests were two-sided. Survival curves were calculated using the Kaplan-Meier method. Regarding statistical methods for genomics and transcriptomics studies, please refer to the corresponding methods section.

## Figures outline

1. Fig 1A-F

2. Fig 2A-C

3. Fig 3A-K

4. Fig 4A-D

5. Fig 5A-F

6. Fig 6A-E

