## Supplemental Figures for "PPM1D is a neuroblastoma oncogene and therapeutic target in childhood neural tumors"

### Supplemental Figures outline

1. Fig S1A-F
2. Fig S2A-G
3. Fig S3A-I
4. Fig S4A-C
5. Fig S5A-B
6. Fig S6A-F

Figure S1

A

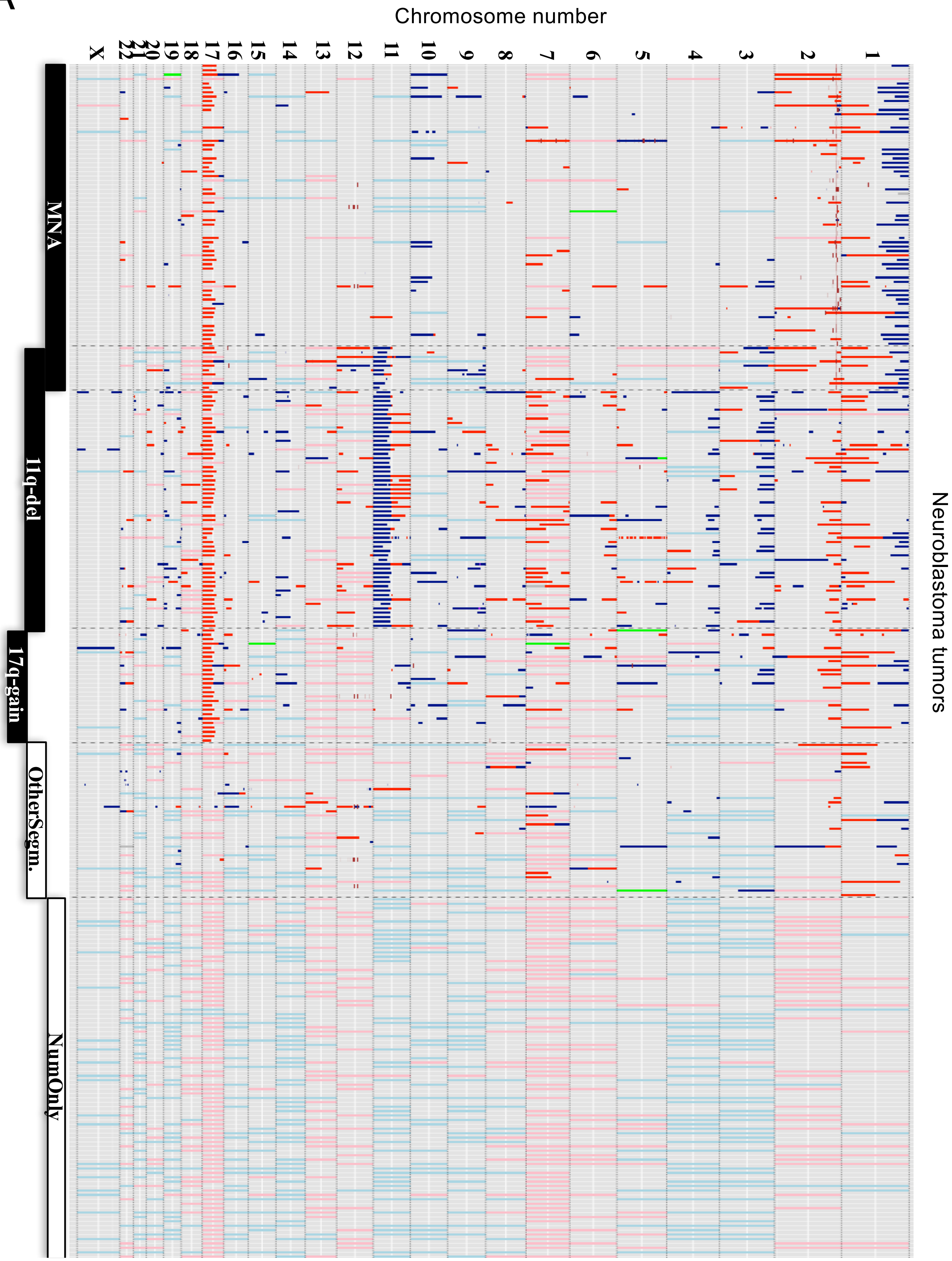
Chromosome number
X
22
21
20
19
18
17
16
15
14
13
12
11
10
9
8
7
6
5
4
3
2
1
MNA
11q-del
17q-gain
OtherSegm.
NumOnly
Neuroblastoma tumors

### Figure S1

B

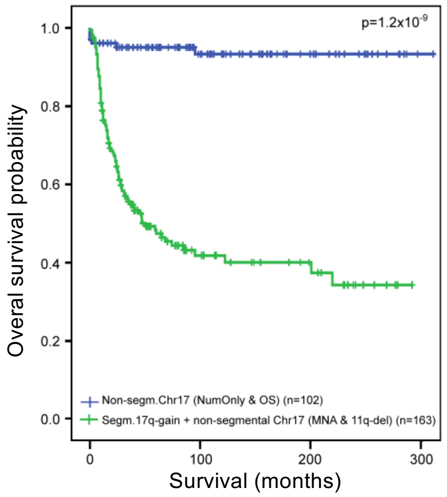


C

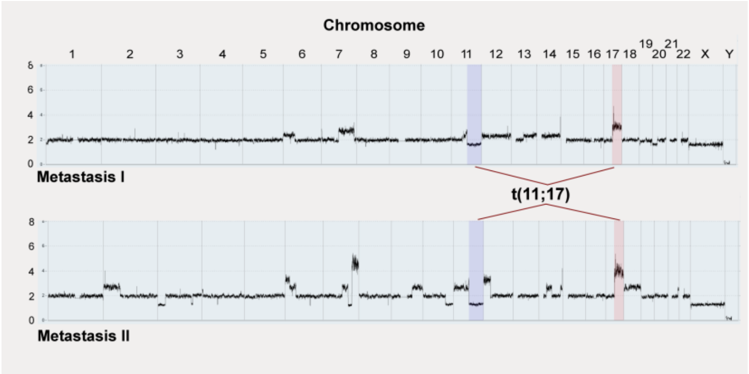


D

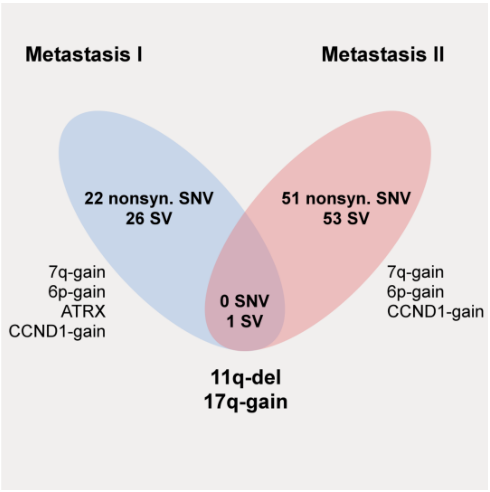


E

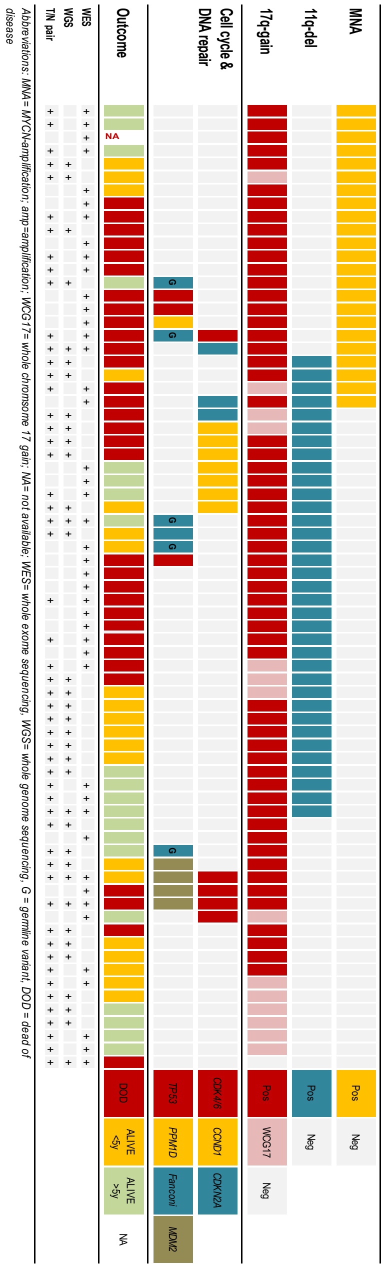


F

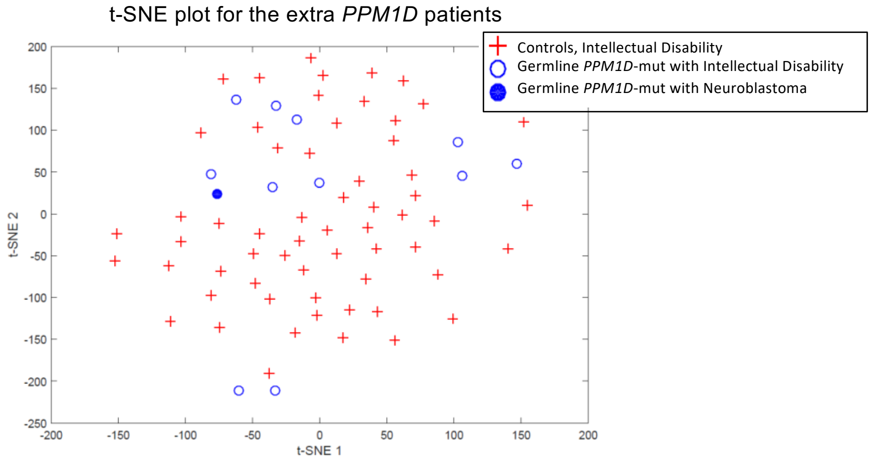

**Figure S1, related to Figure 1. Unfavorable chromosomal 17 q gain is common in neuroblastoma. A.** Gain of chromosome 17q is the most common genetic aberration in neuroblastoma. Summary of SNP array data of gains and losses for neuroblastoma tumors from an unselected cohort of 271 Swedish patients included and analyzed during 25 years. Horizontal lines show segmental loss (clear blue) and gain (clear red) and whole chromosome loss (pale blue) and gain (pale red). Short vertical lines show amplification (red) or homozygous loss (dark blue). Green lines indicate chromothripsis**.** The genomic profile group is indicated to the right. The inclusion features for the three high-risk groups; *MYCN* amplification, 11q deletion and 17q gain-are indicated by ovals. For definition of the genomic groups see Carén et al., 2010. **B.** Gain of 17q correlates with poor survival in neuroblastoma. Neuroblastoma survival probability according to Kaplan-Meier analysis in a Swedish population-based patient material in relation to chromosome 17 status in the tumor tissue, shows significantly worse event-free survival (EFS) for children with 17q segmental gain. Abbreviations; MNA, *MYCN* amplification; 11q-del, 11q-deletion; NumOnly, numerical only (whole chromosome loss or gain); OS, Other segmental (no MNA, no 11q-del, no 17q-gain). **C.** Copy number profiling based on normalized sequencing coverage from two different metastatic sites, at diagnosis (Met 1) and relapse (Met 2), respectively, display different segmental aberrations except for the common 17q-gain and 11q-deletion resulting for one single unbalanced translocation being the first genetic event in neuroblastoma tumor development. **D.** Beside the primary unbalanced translocation t(11;17), all other mutations and structural aberrations showed sequence and/or breakpoint differences although multiple genomic alterations have emerged in parallel in the two different clones (same high-risk neuroblastoma patient as in Fig 1D and Fig S1C). **E.** Genetic landscape of sequenced Swedish neuroblastomas. Diagram of detected genetic defects of specific targets/pathways and clinical parameters (rows) in 73 whole-exome and/or whole-genome sequenced neuroblastoma samples (columns) including one with *PPM1D* mutation as described in detail in Fig 1D and three with *TP53* mutations. **F.** t-SNE Plot of facial dysmorphic features showing the distribution of 12 individuals with germline variants in *PPM1D (n=12, circles)* eleven of which previously described (open circles) (Jansen et al., 2017). and including the patient with neuroblastoma *(filled circle)* versus controls with intellectual disability (ID, red crosses, p=0.0164).

Figure S2

A

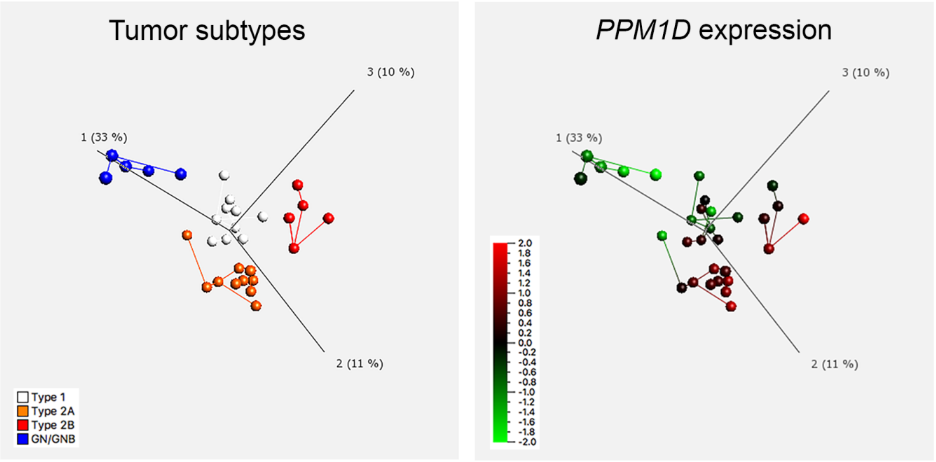


| Group comparison | Fold change | Student's t-test | Significance |
|---|---|---|---|
| Type 2A vs Type 1 | 2.10 | 0.0190 | * |
| Type 2B vs Type 1 | 2.26 | 0.0806 | |
| Type 2A & Type 2B vs Type 1 | 2.15 | 0.0070 | ** |

B

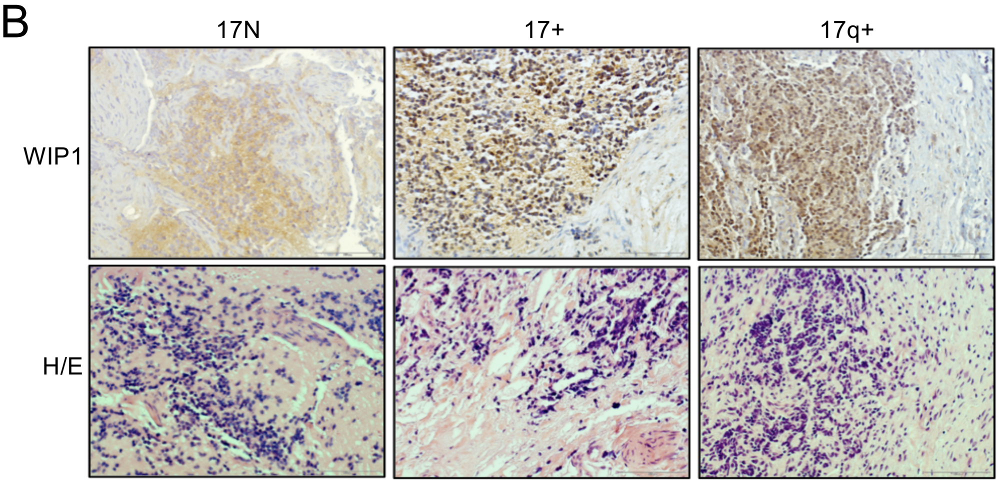


C

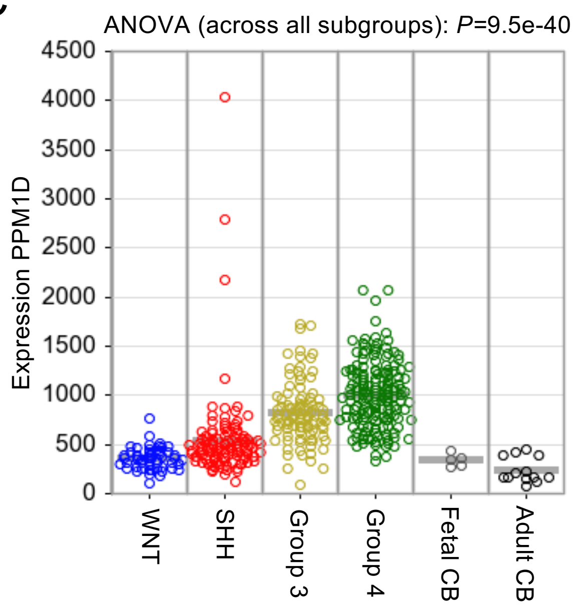


D

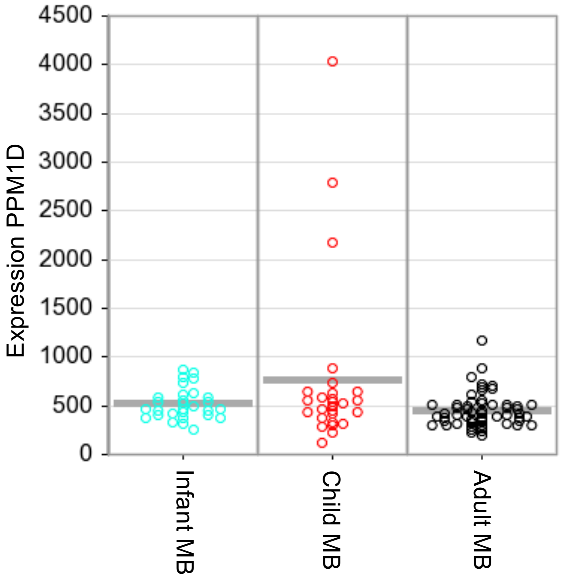

### Figure S2

E

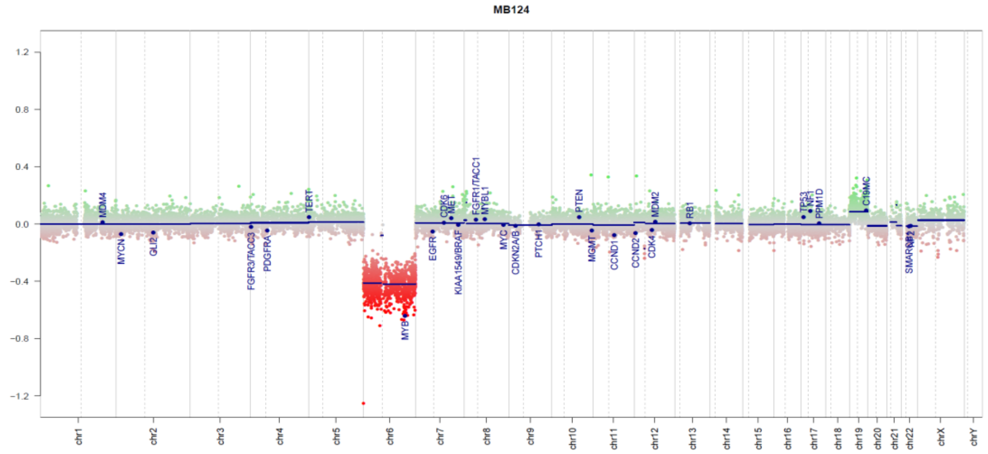


F

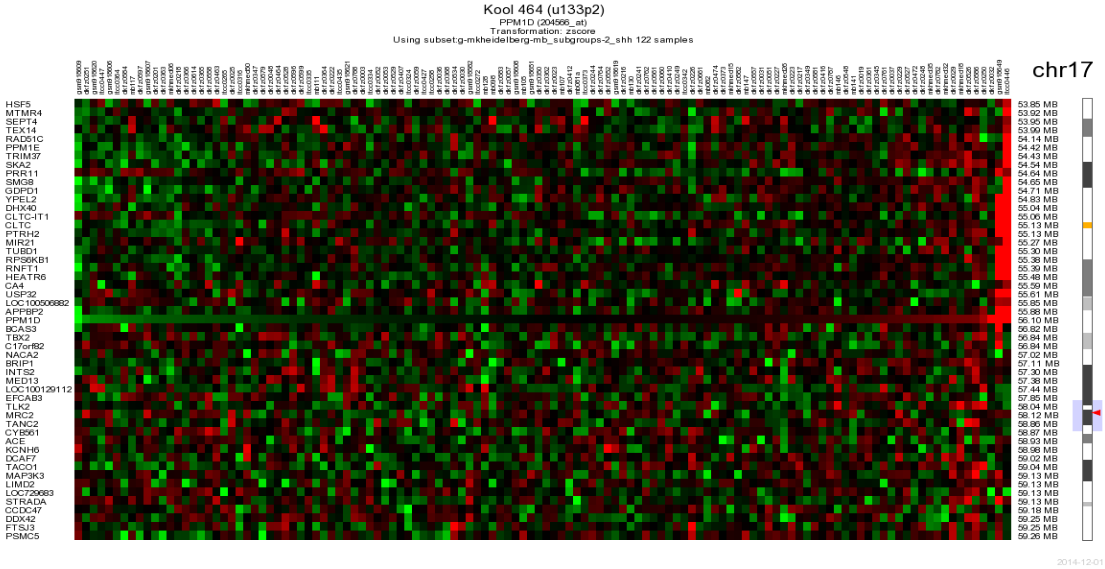


G

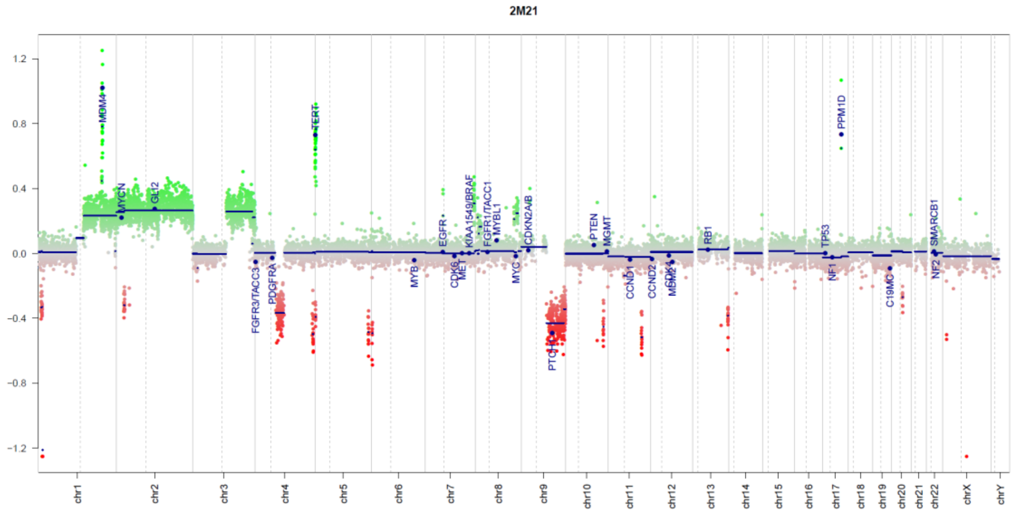

### Figure S2

**Figure S2, related to Figure 2. *PPM1D* expression correlates with unfavorable neuroblastoma and medulloblastoma biological subgroups and is highly expressed in unfavorable chromosome 17q+ tumors.**
**A.** *PPM1D* as a prognostic marker for neuroblastoma. Principal Components Analysis (PCA) of the pre-processed McArdle/Wilzén microarray data sets (716 variables, 30 tumor samples/cases). PCA subgroups corresponding to molecular subtypes of NB; Type 1 (white), Type 2A (orange), Type 2B (red) and ganglioneuroma/ganglioneuroblastoma (GN/GNB; blue), and the expression of *PPM1D* in the four subgroups (Red = High expression; Green = Low expression). Samples/cases are represented as spheres joined by nearest Euclidean neighbor. The color scale of expression is based on standard deviations (SD) ranging from +2 SD (red) to -2 SD (green). Fold change and Student's t-test were calculated and compared between neuroblastoma subtypes; Type 1(n=10), Type 2A (=10), Type 2B (n=5). **B.** Representative images of H/E and Wip1 staining in neuroblastic tumors. Upper images from three neuroblastomas with normal chromosome 17 status (17N), whole chromosome 17 gain (17+), and chromosome 17q gain (17q+). **C.** High *PPM1D* expression correlates with aggressive medulloblastoma subgroups. *PPM1D* expression in medulloblastoma (MB) subgroups (WNT, SHH, Group 3 and Group 4) vs normal fetal or adult cerebellum (CB). *PPM1D* expression is significantly higher in Group 3 and Group 4 medulloblastoma compared to other MB subgroups and to normal adult and fetal cerebellum (Supplementary Table S2). **D.** *PPM1D* expression within SHH MB subgroup only, within the different age groups (infant, child and adult) shows three tumors with high *PPM1D* expression due to high level gene amplification in the SHH-child medulloblastoma subgroup harboring wild-type *TP53*. **E.** Gain-of function PPM1D mutation in medulloblastoma subtypes. Copy number profile (CN) of WNT medulloblastoma case MB124 with a *PPM1D* gain-of-function truncating mutation E525X (allele frequency 0.44). CN figure shows typical profile for a WNT medulloblastoma with loss of chromosome 6. **F.** Amplification of *PPM1D* in medulloblastoma**.** Heatmap of expression data for genes around *PPM1*D in SHH medulloblastomas. Gene amplifications including the *PPM1D* locus were detected in two SHH medulloblastoma samples belonging to the child SHH subgroup with wild-type *TP53*. **G.** Copy number (CN) profile of 2m21 SHH medulloblastoma shows several copy number aberrations, including amplifications of *PPM1D*, *MDM4* and *TERT* and several small deletions across the genome.

Figure S3

A

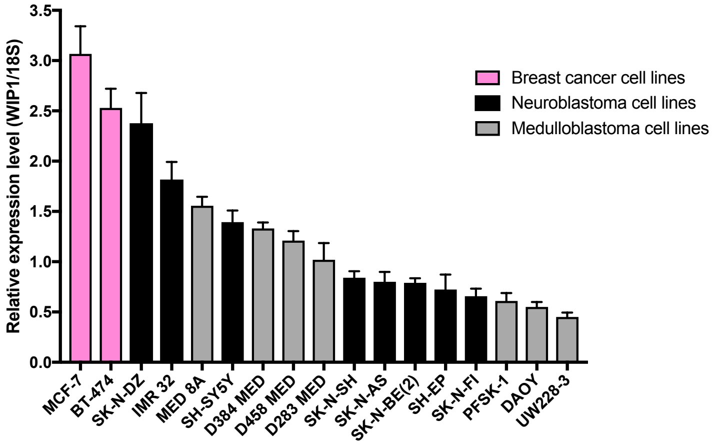


B

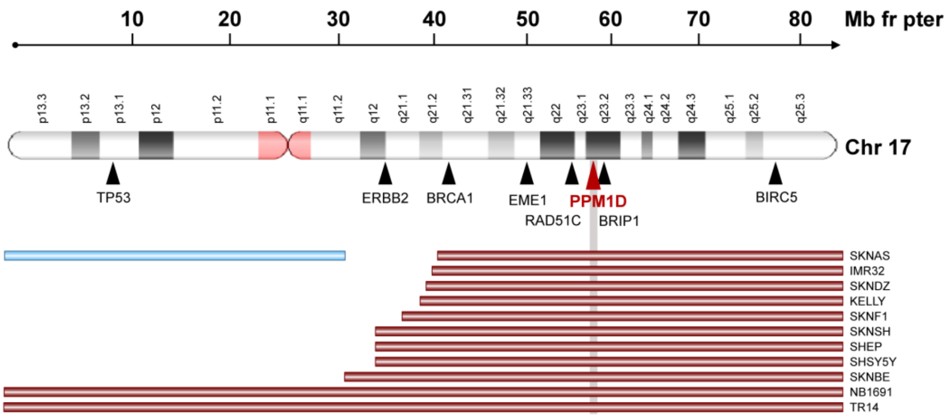


C

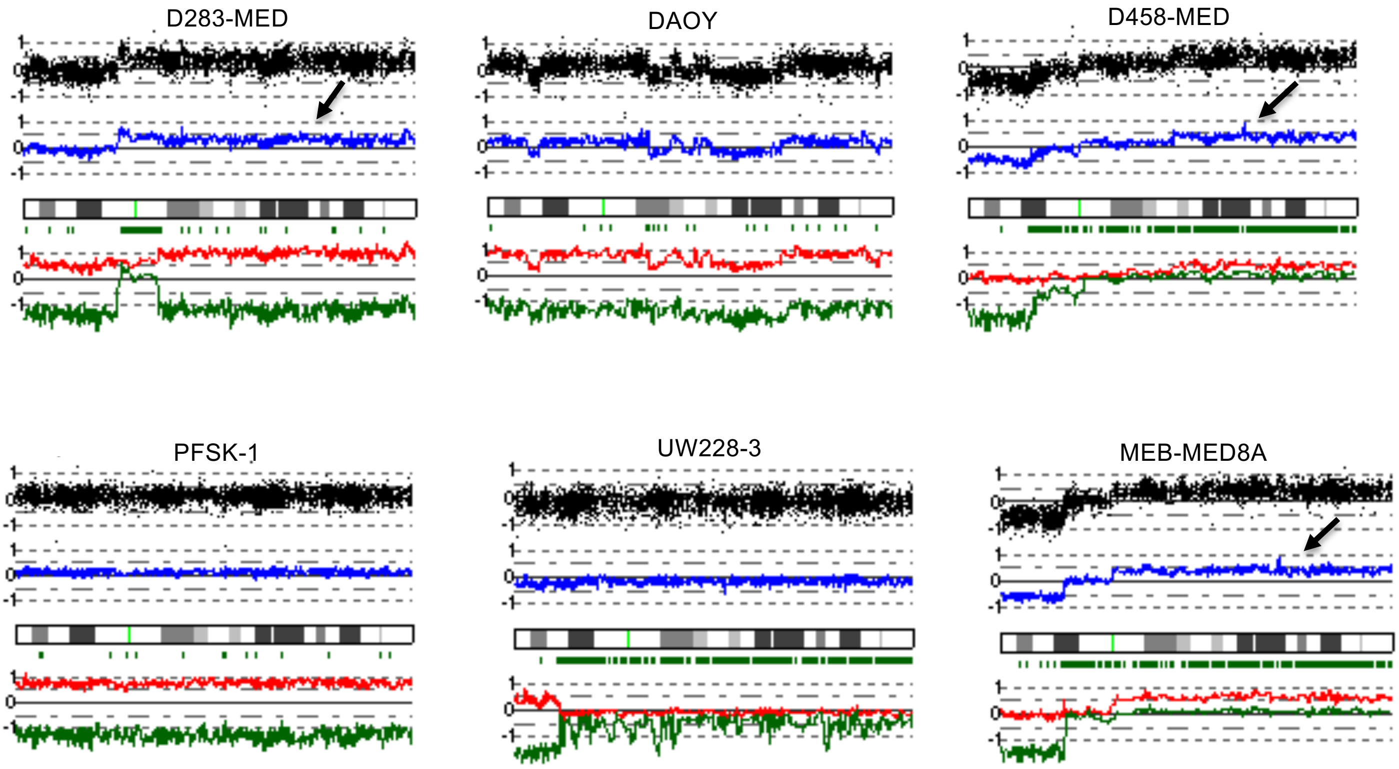

### Figure S3

D

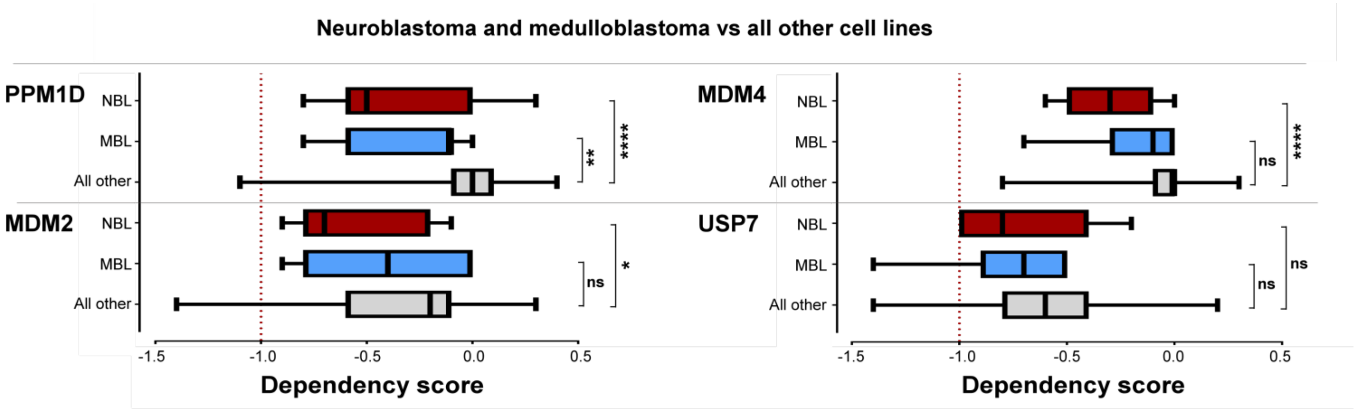


E

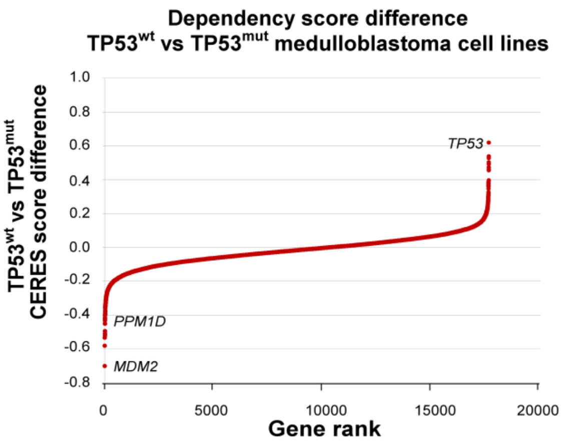


F

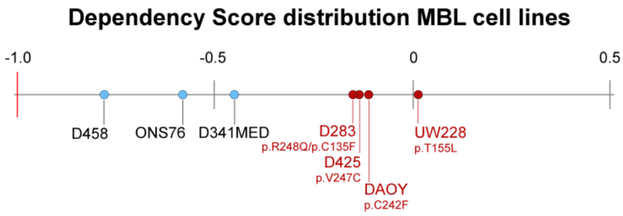


G

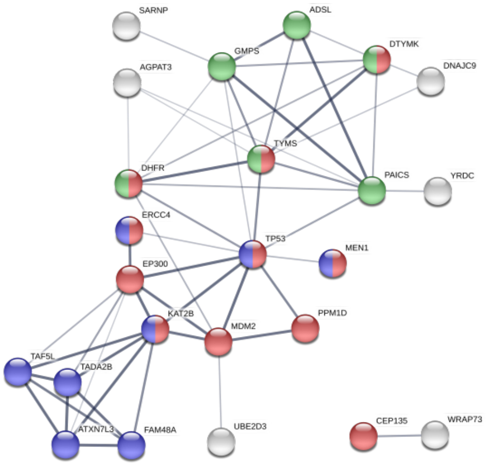


H

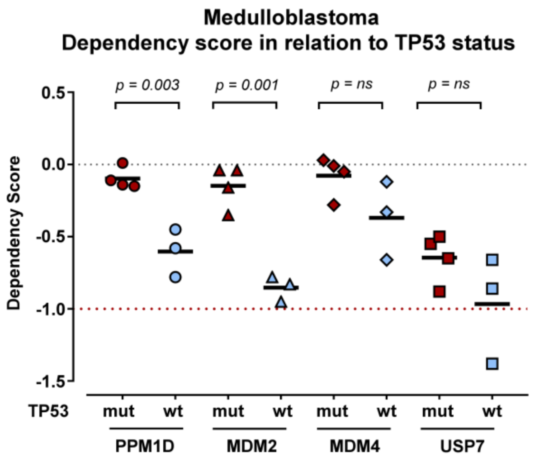

### Figure S3

I

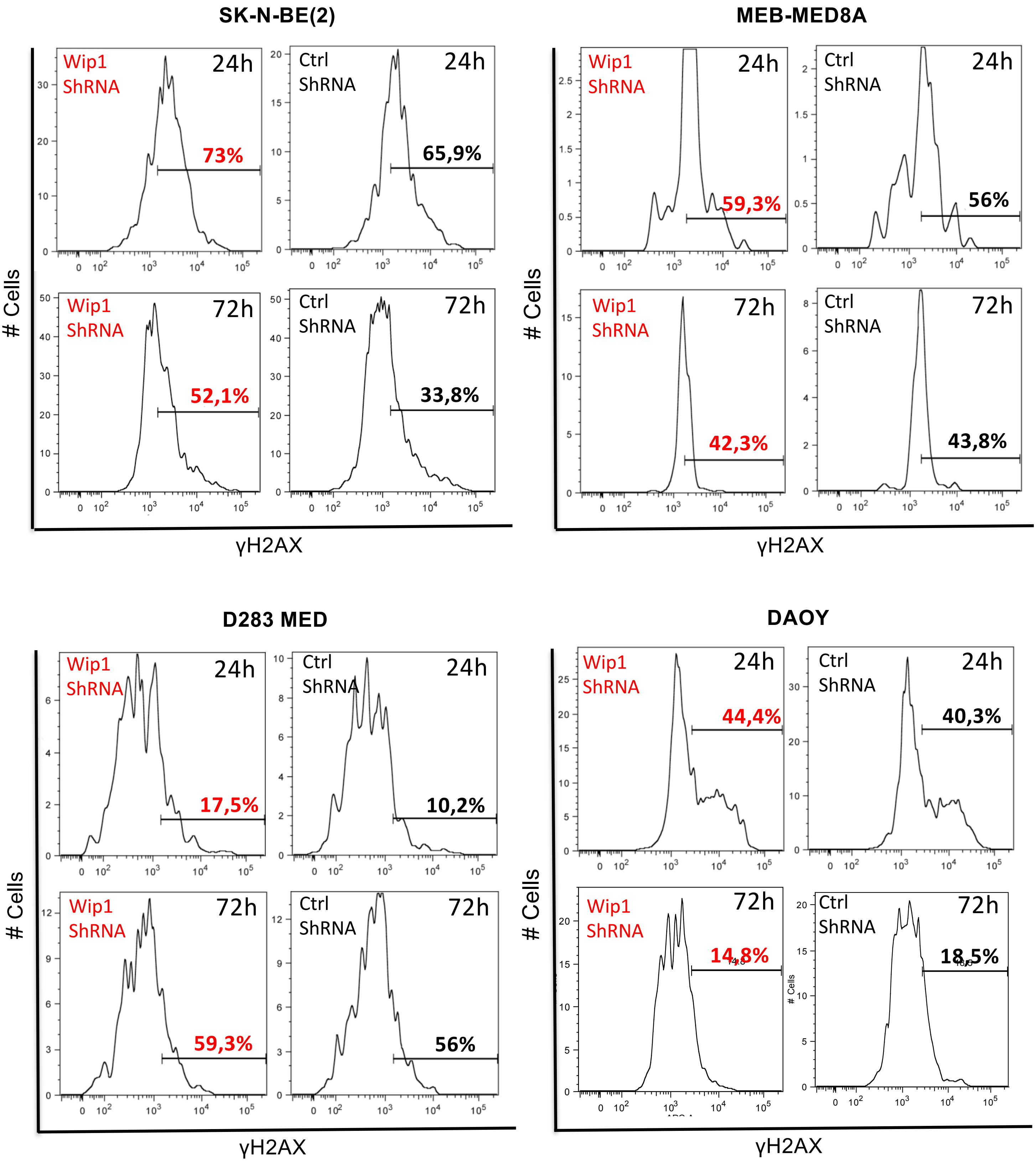

SK-N-BE(2)
Wip1 ShRNA 24h 73%
Ctrl ShRNA 24h 65,9%
Wip1 ShRNA 72h 52,1%
Ctrl ShRNA 72h 33,8%
MEB-MED8A
Wip1 ShRNA 24h 59,3%
Ctrl ShRNA 24h 56%
Wip1 ShRNA 72h 42,3%
Ctrl ShRNA 72h 43,8%
D283 MED
Wip1 ShRNA 24h 17,5%
Ctrl ShRNA 24h 10,2%
Wip1 ShRNA 72h 59,3%
Ctrl ShRNA 72h 56%
DAOY
Wip1 ShRNA 24h 44,4%
Ctrl ShRNA 24h 40,3%
Wip1 ShRNA 72h 14,8%
Ctrl ShRNA 72h 18,5%
### Cells
γH2AX

### Figure S3

**Figure S3, related to Figure 3. *PPM1D* expression correlates to 17q copy number gains and has the highest genetic dependency in *TP53* wild-type neuroblastoma and medulloblastoma cells. Investigating ploidy number of chromosome 17 in neuroblastoma and medulloblastoma cell lines. A.** *PPM1D* is expressed at different levels in neuroblastoma and medulloblastoma cell lines. Relative *PPM1D* mRNA expression in two *PPM1D*-amplified breast cancer cell lines (pink bars), eight neuroblastoma (NB) cell lines (black bars), six medulloblastoma (MB) cell lines (gray bars) and the supratentorial primitive neuroectodermal tumor (sPNET) cell line PFSK-1 (grey bar), analyzed with real-time PCR. The breast cancer cell line MCF-7 is amplified for *PPM1D* and used as a positive control for *PPM1D* mRNA expression. Mean with S.D. of three experiments are displayed. **B.** Chromosome 17q ploidy in neuroblastoma cell lines. Summary of whole or segmental chromosome 17 gain detected or reported in neuroblastoma cell lines (Kryh et al., 2011; Schleiermacher et al., 2004). **C.** Chromosome 17q copy numbers in medulloblastoma cell lines. Comparative genomic hybridization (CGH) array of five medulloblastomas and one supratentorial primitive neuroectodermal tumor cell line PFSK-1, illustrated as a blue line in single chromosome view. The red and green lines show the strongest and weakest allele intensity respectively for each cell line. Arrows indicate gain of chromosome 17q. For allele specific intensity calculations using AsCNAR algorithm see Kryh et al., 2011. **D.** *PPM1D* and *MDM4* show statistically significant differences in genetic dependency in neuroblastoma as compared to all other cell lines. **E.** *PPM1D* expression is important for the survival of medulloblastoma cells lacking *TP53* mutations. Genome-scale CRISPR-Cas9 screening showing ranked average difference in genetic dependencies between wild-type vs mutated TP53 medulloblastoma cell lines. **F.** Dependency score showing high *PPM1D* dependency in wild-type *TP53* medulloblastoma cells. Wild-type *TP53* medulloblastoma cell lines (blue, high dependency) and *TP53* mutated cell lines (red, low dependency) with *TP53* status. **G.** STRING database analysis showing *PPM1D* dependency in wild-type TP53 medulloblastoma cells. Among the top 30 genes with largest different in CERES score there was enrichment of genes involved in negative regulation of cell proliferation (indicated in blue), cell cycle process (red), cellular response to DNA damage (yellow) and chromosome organization (green). The width of the edges corresponds to level of confidence (medium confidence STRING scores of 0.4; high confidence STRING score 0.7; and highest confidence STRING score 0.9). **H.** Medulloblastoma cells with wild-type TP53 are dependent on *PPM1D* expression for survival. Dependency scores of *PPM1D, MDM2, MDM4*, and *USP7* in relation to *TP53* mutational status. **I.** *PPM1D* knockdown increases H2AX phosphorylation. γH2AX was measured with flow-cytometry 24- and 72 hours after transfection with shRNA against *PPM1D* and control shRNA respectively.

Figure S4

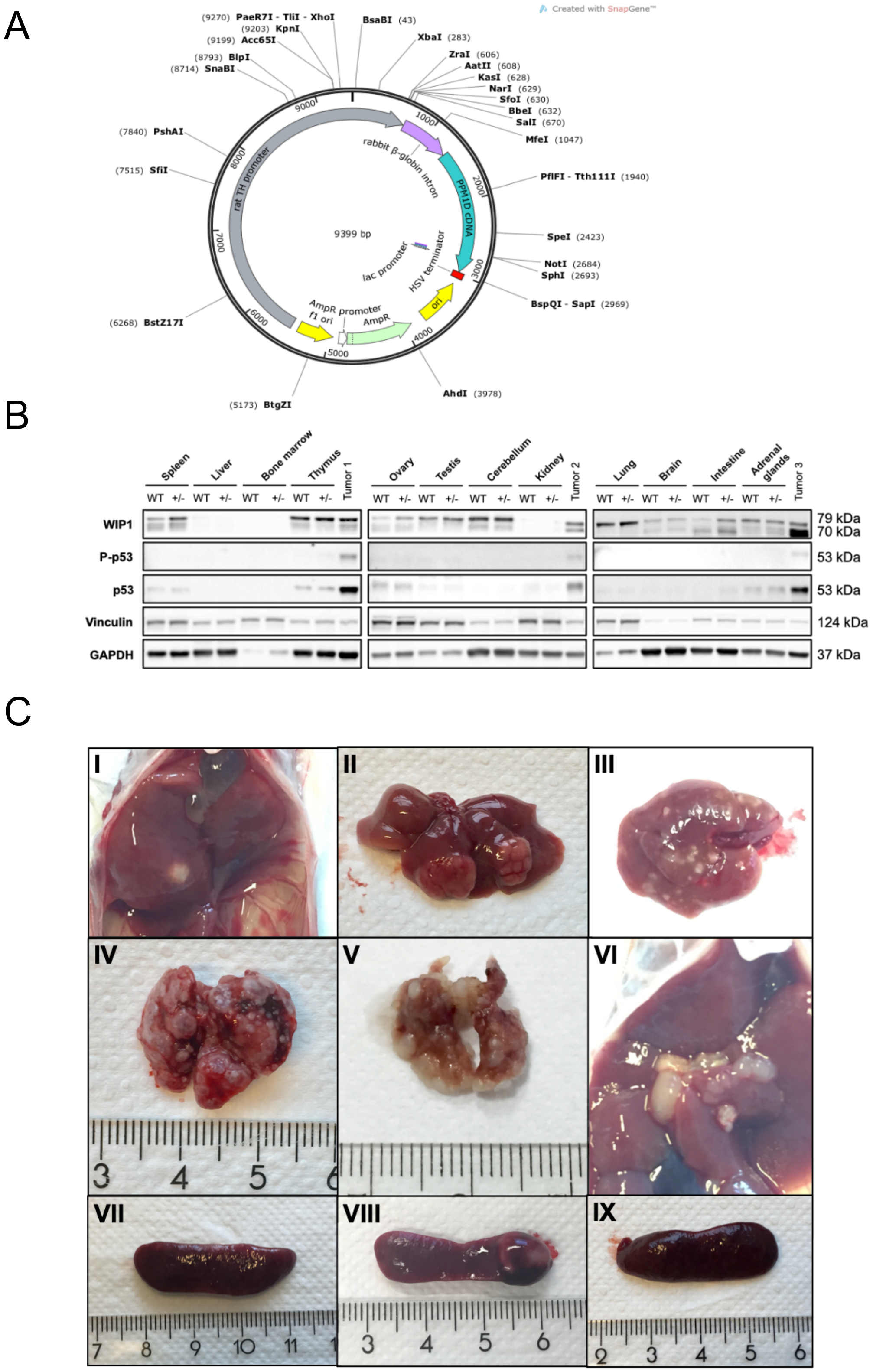


**Figure S4, related to Figure 4. *PPM1D*-transgenic mice show increased WIP1 expression and develop widespread tumors with wild-type p53 accumulation. A.** Transgene-construct. Human *PPM1D* cDNA (blue) *PPM1D,* ligated downstream of the rat tyrosine hydroxylase promoter (gray) and rabbit β-globin enhancer (violet). Herpes simplex virus (HSV) thymidine kinase gene sequence was used as a transcription terminator (red). **B.** Thymic lymphoma shows increased WIP1 protein expression, $p53^{Ser15}$ phosphorylation and accumulation of total p53. Protein expression of WIP1 was analyzed in wild-type and *PPM1D* heterozygous (+/-) mouse tissues. Three different mouse thymic lymphoma tumor samples were also included (Tumor 1-3). Vinculin and GAPDH protein levels were used as loading controls. **C.** PPM1D transgenic mice exhibit widespread tumor malignancies. Representative photographs; tumor mass in the liver (**I**), multiple hepatic metastases (**II, III**), lung carcinosis/metastases (**IV, V**), enlarged lymph nodes (**VI**) and splenomegaly (**VII-IX**) indicative of high tumor burden.

### Figure S5

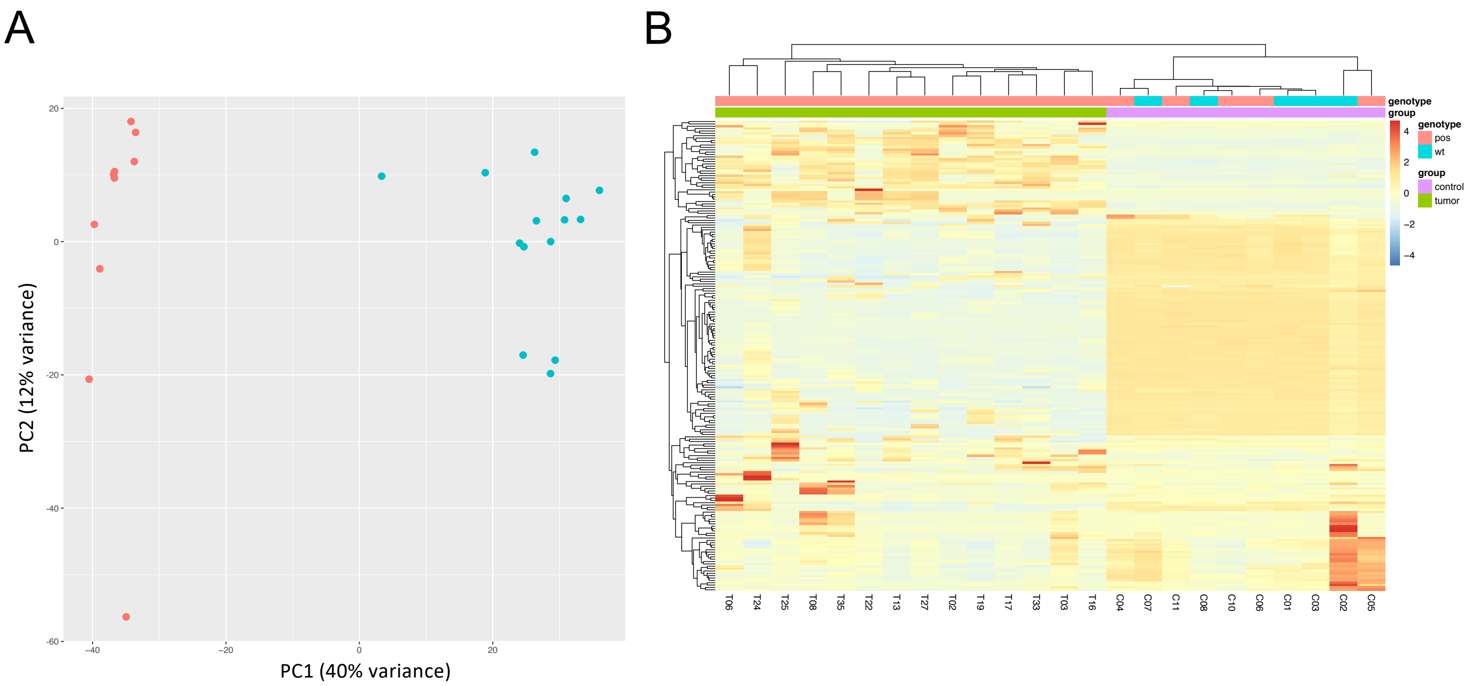


**Figure S5, related to Figure 5. *PPM1D*/WIP1-transgenic mice demonstrate clear distinction between thymic lymphoma tissue and control tissue. A.** Principal component analysis (PCA) plot of all samples, labelled by group (control vs tumor). Red color indicates controls; blue color indicates tumor. **B.** Unsupervised hierarchical clustering showing z scores of the 200 genes with highest standard deviation across all samples. Each column represents one sample (T = tumor; C = control). Each row represents one gene. Samples are annotated according to genotype (*PPM1D* positive or negative) and group (tumor or control). Gene expression values are scaled by row: for each gene, the mean is calculated across all sample. Each sample's deviation from this mean is then indicated by color. Red indicates higher than mean, yellow indicates similar to mean, blue indicates lower than mean.

### Figure S6

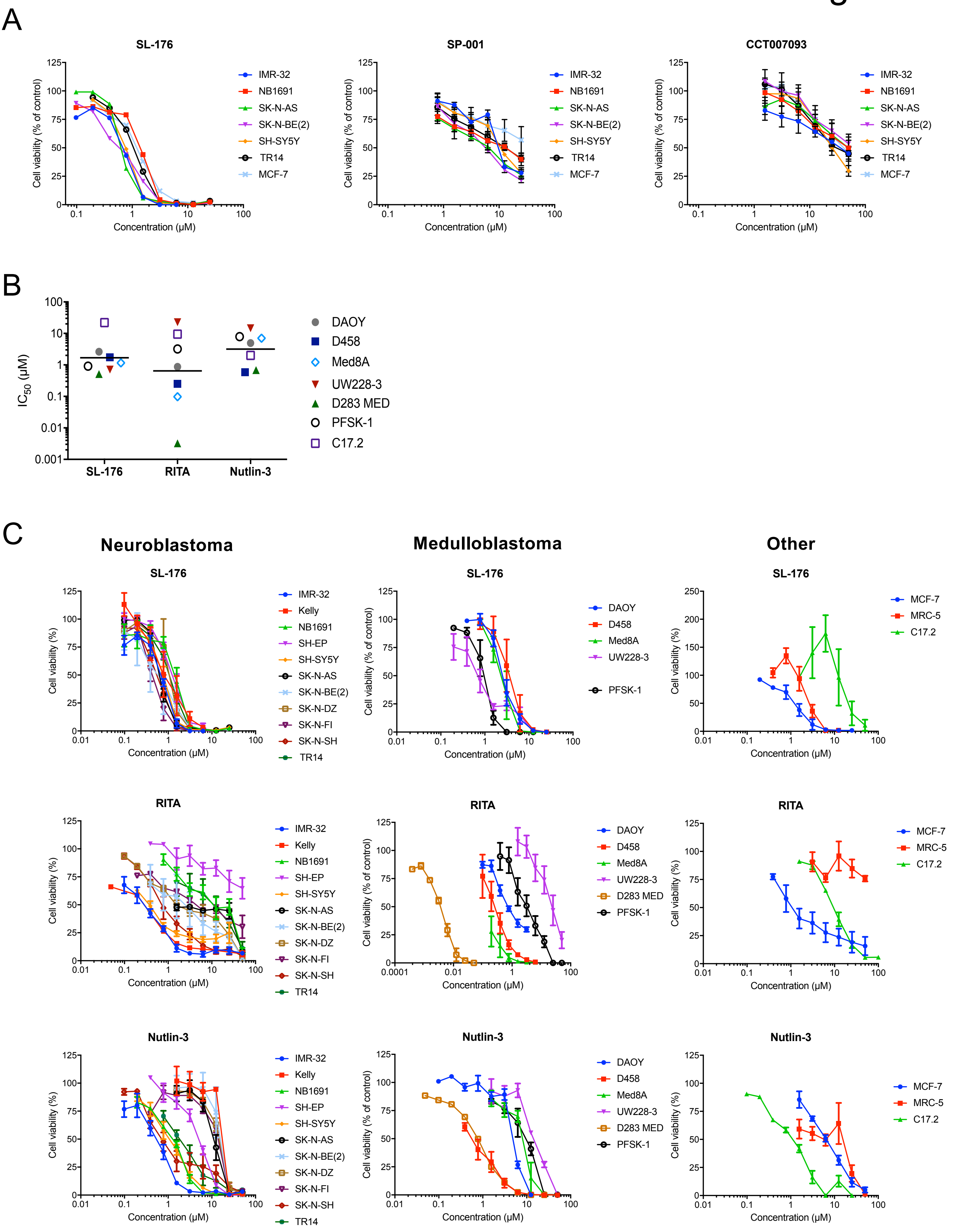
A
SL-176
SP-001
CCT007093
Cell viability (% of control)
Concentration (µM)
IMR-32
NB1691
SK-N-AS
SK-N-BE(2)
SH-SY5Y
TR14
MCF-7
B
IC50 (µM)
SL-176
RITA
Nutlin-3
DAOY
D458
Med8A
UW228-3
D283 MED
PFSK-1
C17.2
C
Neuroblastoma
Medulloblastoma
Other
Cell viability (%)
Kelly
SH-EP
SK-N-DZ
SK-N-FI
SK-N-SH
MRC-5

### Figure S6

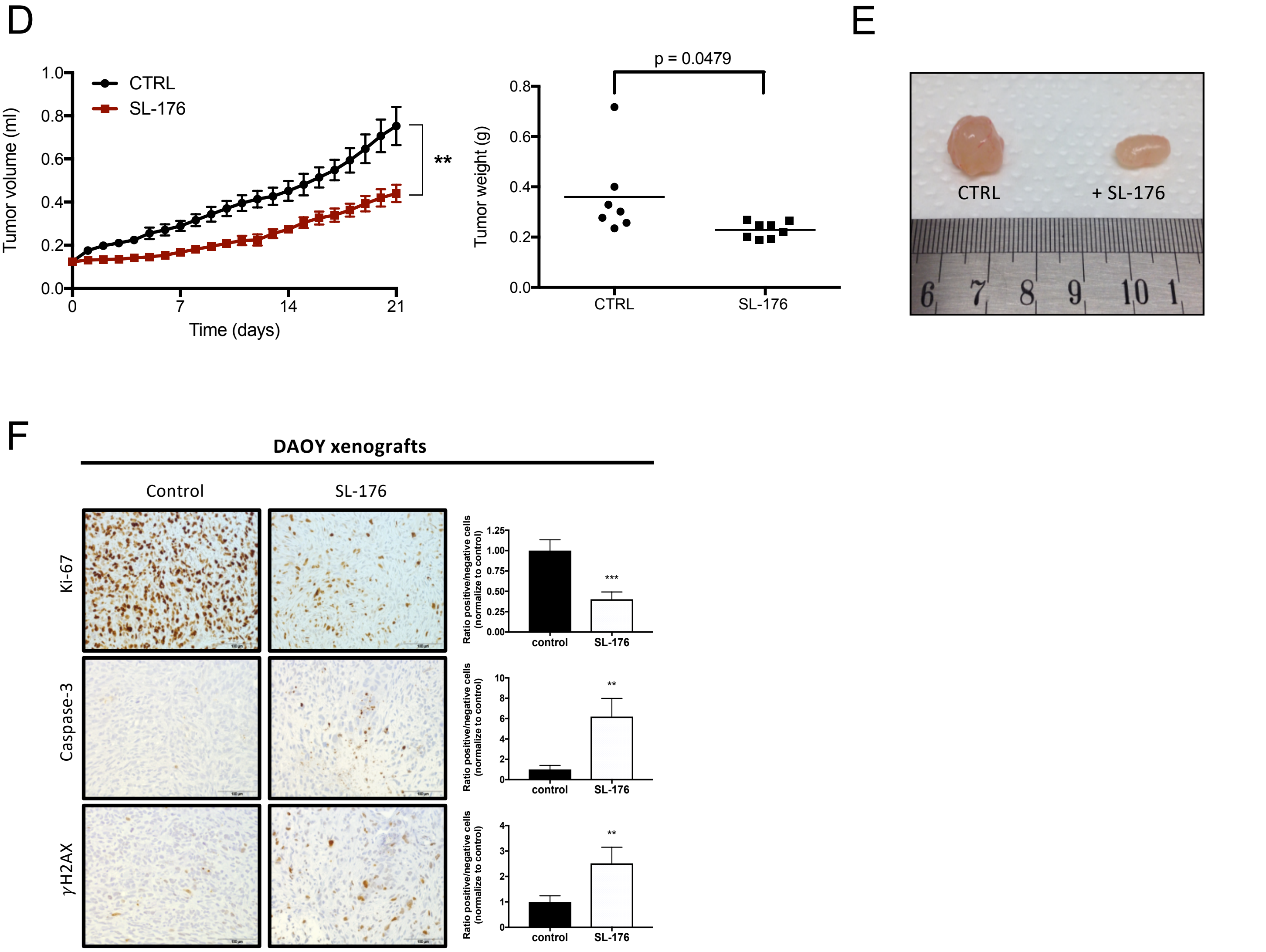


**Figure S6, related to Figure 6. The WIP1 phosphatase inhibitor SL-176 suppresses neuroblastoma and medulloblastoma growth. A.** Dose-response curves for cell viability after 72 hours of treatment with three WIP1 inhibitors; SL-176, SP001 or CCT007093, in neuroblastoma cell lines and breast cancer cell line MCF-7. Cell viability was determined with MTT assay. Data represent the mean ±S.E.M. of at least three experiments. **B.** SL-176 inhibits medulloblastoma cell growth. $IC_{50}$ values for five medulloblastoma cell lines, a supratentorial primitive neuroectodermal (sPNET) tumor cell line PFSK-1 and the murine neural progenitor cell line C17.2 exposed to the specific WIP1 inhibitor SL-176, the p53-MDM2 interaction inhibitor RITA or the MDM2 antagonist Nutlin-3. Horizontal lines indicate mean. No significant differences between the mean $IC_{50}$ values in medulloblastoma cell lines (mean $IC_{50}$ for SL-176: 1.3 µM, RITA: 1.1 µM and Nutlin-3: 4.7 µM, one-way ANOVA $P$=0.26). $IC_{50}$ values were calculated from results from cell viability assay WST-1 performed at least three times. MRC-5 and C17.2 were used as a non-tumorigenic control for drug toxicity. **C.** Dose-response curves for cell viability after 72 hours of treatment with SL-176, RITA or Nutlin-3 in neuroblastoma cell lines, medulloblastoma cell lines, sPNET cell line PFSK-1 and breast cancer cell line MCF-7. The fibroblast cell line MRC-5 and the murine neural progenitor cell line C17.2 were used as non-tumorigenic controls for drug toxicity. Cell viability was determined with WST-1. Data represent the mean ±S.E.M. of at least three experiments. **D.** SL-176 inhibit medulloblastoma growth *in vivo*. Nude mice engrafted with medulloblastoma DAOY xenografts were treated from tumor volume 0.12 mL, receiving either daily i. p. injections of SL-176 (n=8) for 21 days or no treatment (n=7). SL-176 treatment significantly delayed medulloblastoma xenograft growth (t-test, day 21 $P$=0.0051) (left, mean with S.E.M. are displayed), and significantly reduced tumor weight after 21 days ($P$=0.0479, horizontal bars = mean weight) (right).
**E.** Representative photograph of dissected medulloblastoma xenograft in comparison. **F.** SL-176 decrease proliferation, induce apoptosis and activate γH2AX in xenograft tumors. Immunohistochemical analysis of DAOY xenograft tumors. Tumor sections were stained with anti-Ki-67, anti-Caspase 3, and anti γH2AX antibodies. Representative examples of immunostaining are shown. Images were acquired at 400X magnification. Identification and quantification of positive and negative cells was carry out with ImageJ software.
