## Supplemental Tables for "PPM1D is a neuroblastoma oncogene and therapeutic target in childhood neural tumors"

*Table S1. Summary of 17q variation, Related to Figure 1C.*

| | PROPORTION OF CASES | PATIENT ID | NUMERICAL ONLY | MYCN AMPLIFIED | STRUCTURAL |
|---|---|---|---|---|---|
| **17q GAIN IN STEM (NO INTRATUMORAL VARIATION)** | 57% (13/23) | NB1, NB2 NB3, NB5, NB6, NB7, NB8, NB9, NB12, NB13, NB17, NB19, NB22 | 7/13 | 3/13 | 3/13 |
| **INTRATUMORAL 17q ACCUMULATION** | 30% (7/23) | NB11*, NB14*, NB16**, NB18*, NB20**, NB21*, NB23** | 0 | 3/7 | 4/7 |
| **17q WITH REGIONAL FLUCTUATIONS** | 13% (3/23) | NB4, NB10, NB15 | 1/3 | 1/3 | 1/3 |

Evolutionary analysis of *PPM1D*/17q copy number yielded three categories of tumors, those with *PPM1D*/17q gain present only in the stem with no further evolution, those with intratumoral accumulation of PPM1D/17q through clonal evolution and those where PPM1D/17q copy number fluctuated to an extent that precluded analysis. Patient IDs refer to those in Karlsson et al., 2018. Each category is the further subdivided into low-risk patients (numerical only) or high-risk patients, either MYCN amplified or dominated by structural changes (STR).

*Table S1. Copy number status of PPM1D/17q, Related to Figure 1C.*

| Tumor ID | Sample ID | Chr | Start | End | Med LogR | Type | Method | Cytoband/Gene | Clone size (%) |
|---|---|---|---|---|---|---|---|---|---|
| **NO INTRATUMORAL COPY NUMBER VARIATION: 17q GAIN IN STEM** | | | | | | | | | |
| NB1 | ALL SAMPLES (3) | 17 | 0 | 83257441 | NA | GAIN x2 | SNP Array | WHOLE | 100 |
| NB2 | ALL SAMPLES (3) | 17 | 0 | 81041938 | NA | GAIN x3 | SNP Array | WHOLE | 100 |
| NB3 | ALL SAMPLES (2) | 17 | 0 | 81041938 | 0.47 | GAIN x2 | SNP array | WHOLE | 100 |
| NB5 | ALL SAMPLES (6) | 17 | 400959 | 80263427 | NA | GAIN x2 | SNP array | WHOLE | 100 |
| NB6 | ALL SAMPLES (4) | 17 | 400959 | 80263427 | NA | GAIN x2 | SNP array | WHOLE | 100 |
| NB7 | ALL SAMPLES (6) | 17 | 400959 | 80263427 | NA | GAIN x2 | SNP array | WHOLE | 100 |
| NB8 | ALL SAMPLES (3) | 17 | 400959 | 80263427 | 0.70 | GAIN x2 | SNP array | WHOLE | 100 |
| NB9 | ALL SAMPLES (2) | 17 | 28183784 | 81041938 | NA | GAIN | SNP array | 17q11q25 | 100 |
| NB12 | ALL SAMPLES (3) | 17 | 19143975 | 81041938 | 0.70 | GAIN x2 | SNP array | 17q11q25 | 100 |
| NB13 | ALL SAMPLES (4) | 17 | 35914854 | 80956017 | NA | GAIN x3 | SNP array | 17q12q25 | 100 |
| NB17 | ALL SAMPLES (4) | 17 | 36412891 | 80652682.5 | NA | GAIN | SNP array | 17q12q25 | 100 |
| NB19 | ALL SAMPLES (10) | 17 | 43766497 | 80263427 | NA | GAIN x2 | SNP array | 17q21q25 | 100 |
| NB22 | ALL SAMPLES (12) | 17 | 45836832 | 80263427 | NA | GAIN | SNP array | 17q21q25 | 100 |
| **REGIONAL ACCUMULATION OF 17q COPIES COMPARED TO STEM LINE** | | | | | | | | | |
| *17q+ IN STEM FOLLOWED BY ADDITIONAL GAINS IN BRANCHES* | | | | | | | | | |
| NB11 | ALL SAMPLES (4) | 17 | 0 | 81027137 | NA | CNNI GAIN | SNP array | WHOLE | 100 |
| NB11 | B2 | 17 | 0 | 81027137 | 0.05 | GAIN | SNP array | WHOLE | 30 |
| NB11 | B3 | 17 | 400959 | 80263427 | 0.11 | GAIN | SNP array | WHOLE | 55 |
| NB11 | B4 | 17 | 400959 | 80263427 | 0.21 | GAIN | SNP array | WHOLE | 100 |
| NB14 | ALL SAMPLES (4) | 17 | 400959 | 80263427 | NA | GAIN | SNP array | WHOLE | 100 |
| NB14 | B3 | 17 | 0 | 81027137 | 0.18 | GAIN x2 | SNP array | WHOLE | 60 |
| NB14 | M | 17 | 0 | 81027137 | 0.08 | GAIN x2 | SNP array | WHOLE | 100 |
| NB18 | ALL SAMPLES (7) | 17 | 44788545 | 81041937 | NA | GAIN | SNP array | 17q21q25 | 100 |
| NB18 | B | 17 | 33225118 | 81041937 | 0.38 | GAIN x2 | SNP array | 17q12q21 | 100 |
| NB18 | P2 | 17 | 33339133 | 44165802 | 0.13 | GAIN | SNP array | 17q12q21 | 100 |
| NB18 | P2 | 17 | 0 | 33339133 | -0.16 | LOSS | SNP array | 17p13q12 | 100 |
| NB18 | P3 | 17 | 33339133 | 44165802 | 0.22 | GAIN | SNP array | 17q12q21 | 100 |
| NB18 | P4 | 17 | 33339133 | 81195210 | 0.16 | GAIN x2-3 | SNP array | 17q12q21 | 30 |
| NB18 | M1 | 17 | 33339133 | 81041937 | 0.52 | GAIN x2 | SNP array | 17q12q21 | 100 |
| NB18 | M2 | 17 | 33339133 | 81041937 | 0.45 | GAIN x2 | SNP array | 17q12q21 | 100 |
| NB21 | ALL SAMPLES (2) | 17 | 39613689 | 81041937 | NA | GAIN | SNP array | 17q21q25 | 100 |
| NB21 | B1 | 17 | 39488988 | 80263427 | 0.49 | GAIN x2 | SNP array | 17q21q25 | 60 |
| *17q+ IN BRANCHES ONLY, BUT BECOMES FIXED REGIONALLY THROUGH CLONAL SWEEP* | | | | | | | | | |
| NB16 | ALL SAMPLES (2) | 17q normal copy number | | | | | | | |
| NB16 | B1 | 17 | 54967528 | 80263427 | 0.41 | GAIN | SNP array | 17q22q25 | 80 |
| NB16 | B2 | 17 | 54900876 | 81195210 | 0.41 | GAIN | SNP array | 17q22q25 | 100 |
| NB16 | B3 | 17 | 54967528 | 80263427 | 0.48 | GAIN | SNP array | 17q22q25 | 100 |
| NB20 | ALL SAMPLES (4) | 17q normal copy number | | | | | | | |
| NB20 | B2 | 17 | 0 | 83257441 | NA | GAIN | SNP array | WHOLE | 100 |
| NB20 | B2 | 17 | 0 | 83257441 | 0.37 | GAINx2 | SNP array | WHOLE | 30 |
| NB20 | B4 | 17 | 0 | 83257441 | 0.24 | GAIN | SNP array | WHOLE | 100 |
| NB20 | B1 | 17 | 38866764 | 83257441 | 0.30 | GAIN | SNP array | 17q21q25 | 60 |
| NB20 | B3 | 17 | 38866764 | 83257441 | 0.31 | GAIN | SNP array | 17q21q25 | 75 |
| NB23 | ALL SAMPLES (8) | 17q normal copy number | | | | | | | |
| NB23 | P1 | 17 | 38195495 | 63495631 | 0.15 | GAIN | SNP array | 17q21q24 | 100 |
| NB23 | P2 | 17 | 38195495 | 63495631 | 0.13 | GAIN | SNP array | 17q21q24 | 100 |
| NB23 | M1 | 17 | 43262179 | 81041938 | 0.34 | GAIN | SNP array | 17q21q25 | 100 |
| NB23 | M2 | 17 | 43352049 | 80263427 | 0.37 | GAIN | SNP array | 17q21q25 | 100 |
| NB23 | M3 | 17 | 43268712 | 80263427 | 0.40 | GAIN | SNP array | 17q21q25 | 100 |
| NB23 | R1 | 17 | 43266347 | 80263427 | 0.31 | GAIN | SNP array | 17q21q25 | 100 |
| NB23 | R2A | 17 | 43266347 | 80263427 | 0.32 | GAIN | SNP array | 17q21q25 | 100 |
| NB23 | R2B | 17 | 43266347 | 80263427 | 0.31 | GAIN | SNP array | 17q21q25 | 100 |
| **GAIN OF 17q FLUCTUATES IN SUBCLONES** | | | | | | | | | |
| NB4 | ALL SAMPLES (3) | 17q normal copy number | | | | | | | |
| NB4 | B1 | 17 | 40086266 | 81021494 | 0.33 | GAIN x2 | SNP array | 17q21q25 | 40 |
| NB4 | B2 | 17 | 38074518 | 80263427 | 0.41 | GAIN x2 | SNP array | 17q12q25 | 50 |
| NB4 | B3 | 17 | 38074518 | 80263427 | 0.36 | GAIN x2 | SNP array | 17q12q25 | 40 |
| NB10 | ALL SAMPLES (2) | 17q normal copy number | | | | | | | |
| NB10 | B2 | 17 | 400959 | 80263427 | 0.27 | GAIN | SNP array | WHOLE | 100 |
| NB15 | ALL SAMPLES (2) | 17q normal copy number | | | | | | | |
| NB15 | B1 | 17 | 38299547 | 81041937 | 0.57 | GAIN x4 | SNP array | 17q22q25 | 50 |
| NB15 | B2 | 17 | 38299547 | 81041937 | 0.53 | GAIN x2 | SNP array | 17q22q25 | 65 |

Chromosome segments encompassing *PPMID1* (pos 17:60,600,183-60,666,280; GRCh38.p12) filtered from the genomic array raw data of Karlsson et al., 2018.
Chr, chromosomes; Med LogR, median log2 ratio; NA, not applicable (fusion of several samples); Cytoband, cytogenetic position where WHOLE denotes chromosome 17 aneuploidy.

| For Fisher's test | F (NUM ONLY STEM) | U (MYCN AMP/STRU( | Fisher P |
|---|---|---|---|
| 17q+ IN STEM ONLY | 7 | 6 | |
| 17q+ ACCUMULATION | 0 | 7 | 0.0445 |

*Table S2. Differential expression of PPM1D in medulloblastoma, Related to Supplementary Figure S2C.*

| Group comparison | ANOVA *P* -value |
|---|---|
| WNT vs fetal CB | NS |
| WNT vs adult CB | 1.90E-03 |
| SHH vs fetal CB | NS |
| SHH vs adult CB | 0.02 |
| Group 3 vs fetal CB | 1.20E-03 |
| Group 3 vs adult CB | 3.50E-09 |
| Group 4 vs fetal CB | 1.80E-05 |
| Group 4 vs adult CB | 6.30E-14 |
| SHH vs WNT | 2.20E-03 |
| Group 3 vs WNT | 2.50E-19 |
| Group 4 vs WNT | 2.70E-32 |
| Group 3 vs SHH | 3.10E-07 |
| Group 4 vs SHH | 6.20E-21 |
| Group 4 vs Group 3 | 2.20E-05 |
| All MB vs all CB | 1.20E-06 |
| All subgroups | 9.50E-40 |

*Table S3, Neuroblastoma Rang. Related to Figure 3.*

| Rang | GENE | Δ TP53 CERES | Inverse Rang | GENE | Δ TP53 CERES |
|---|---|---|---|---|---|
| 1 | **PPM1D (8493)** | **-0.631131289** | 1 | TBC1D3 (729873) | 0.775875188 |
| 2 | DNAJC9 (23234) | -0.599875082 | 2 | **TP53 (7157)** | **0.439444935** |
| 3 | **MDM2 (4193)** | **-0.561873035** | 3 | TRNT1 (51095) | 0.36862162 |
| 4 | UBE2D3 (7323) | -0.505078942 | 4 | AP2M1 (1173) | 0.358958334 |
| 5 | CCNL1 (57018) | -0.438240784 | 5 | TPRKB (51002) | 0.355877218 |
| 6 | TBC1D3C (414060) | -0.424124013 | 6 | FARSA (2193) | 0.346963178 |
| 7 | TSC2 (7249) | -0.390902601 | 7 | WDR61 (80349) | 0.342824194 |
| 8 | GOLGA6L1 (283767) | -0.385998656 | 8 | GGPS1 (9453) | 0.329380918 |
| 9 | SLC35A1 (10559) | -0.356019496 | 9 | SGOL1 (151648) | 0.327991152 |
| 10 | C17orf58 (284018) | -0.352492248 | 10 | CCT7 (10574) | 0.324015827 |
| 11 | KIF18B (146909) | -0.351675321 | 11 | FBXO42 (54455) | 0.32134656 |
| 12 | DBF4 (10926) | -0.346152435 | 12 | EIF3C (8663) | 0.320357 |
| 13 | ARID1B (57492) | -0.345497012 | 13 | CHMP3 (51652) | 0.318879061 |
| 14 | TCEB3CL (728929) | -0.341972884 | 14 | CKS1B (1163) | 0.314530061 |
| 15 | DNAJA1 (3301) | -0.336090826 | 15 | METTL14 (57721) | 0.305273576 |
| 16 | USP17L5 (728386) | -0.335809357 | 16 | NRF1 (4899) | 0.303539139 |
| 17 | GDF11 (10220) | -0.33373291 | 17 | CTU1 (90353) | 0.301010721 |
| 18 | SERBP1 (26135) | -0.33055467 | 18 | DDX20 (11218) | 0.300524517 |
| 19 | PIAS1 (8554) | -0.32996042 | 19 | PGM3 (5238) | 0.290831529 |
| 20 | ABHD11 (83451) | -0.329150015 | 20 | GRAP (10750) | 0.290196852 |
| 21 | **MDM4 (4194)** | **-0.328117688** | 21 | ATXN10 (25814) | 0.287514623 |
| 22 | ADRM1 (11047) | -0.327744234 | 22 | DDX19B (11269) | 0.281335507 |
| 23 | CELF2 (10659) | -0.322795413 | 23 | FAM25A (643161) | 0.277860595 |
| 24 | HMGA1 (3159) | -0.317861286 | 24 | C10orf71 (118461) | 0.277855566 |
| 25 | CDAN1 (146059) | -0.316361488 | 25 | ANAPC10 (10393) | 0.276814536 |
| 26 | TM2D3 (80213) | -0.314965456 | 26 | VPS52 (6293) | 0.274229534 |
| 27 | NIP7 (51388) | -0.314925046 | 27 | RBM33 (155435) | 0.273833023 |
| 28 | TSC1 (7248) | -0.313363759 | 28 | DOHH (83475) | 0.268896118 |
| 29 | AMBRA1 (55626) | -0.306444369 | 29 | SRP9 (6726) | 0.268234457 |
| 30 | RBM34 (23029) | -0.305637315 | 30 | ELP6 (54859) | 0.261195868 |
| 31 | **USP7 (7874)** | **-0.304581651** | 31 | ZC3H4 (23211) | 0.260887717 |
| 32 | CYP27B1 (1594) | -0.301301017 | 32 | TRMT61A (115708) | 0.260454742 |
| 33 | TERF1 (7013) | -0.300081552 | 33 | C11orf30 (56946) | 0.257133212 |
| 34 | FRG1 (2483) | -0.299988853 | 34 | WARS (7453) | 0.254638686 |
| 35 | CDCA8 (55143) | -0.299967188 | 35 | IGF1R (3480) | 0.25323441 |
| 36 | ACD (65057) | -0.295033425 | 36 | AGPAT1 (10554) | 0.252980027 |
| 37 | RAP2A (5911) | -0.2944985 | 37 | UPF1 (5976) | 0.251354802 |
| 38 | SLC7A11 (23657) | -0.294235542 | 38 | RPRD1B (58490) | 0.249761037 |
| 39 | GRASP (160622) | -0.294186423 | 39 | MAU2 (23383) | 0.248716941 |
| 40 | COMMD6 (170622) | -0.294131838 | 40 | NIPBL (25836) | 0.248679834 |
| 41 | OR4F21 (441308) | -0.293934487 | 41 | ANAPC5 (51433) | 0.24601185 |
| 42 | EIF4EBP1 (1978) | -0.293583933 | 42 | AP2S1 (1175) | 0.245773384 |
| 43 | CSH2 (1443) | -0.292640121 | 43 | RTCB (51493) | 0.244673731 |
| 44 | CCDC137 (339230) | -0.292088423 | 44 | NAA20 (51126) | 0.244332473 |
| 45 | CPNE7 (27132) | -0.291115758 | 45 | ELP2 (55250) | 0.244111002 |
| 46 | BCAS3 (54828) | -0.289571385 | 46 | MAEA (10296) | 0.243886056 |
| 47 | RASGRP1 (10125) | -0.287776037 | 47 | GNGT2 (2793) | 0.243467888 |
| 48 | PRAMEF5 (343068) | -0.285392269 | 48 | TIMM9 (26520) | 0.242709657 |
| 49 | C17orf96 (100170841) | -0.283872068 | 49 | COPS3 (8533) | 0.241580455 |
| 50 | UBE2O (63893) | -0.283467297 | 50 | KBTBD8 (84541) | 0.240848847 |
| 51 | TTC33 (23548) | -0.279363627 | 51 | PIM1 (5292) | 0.239995107 |
| 52 | ZNF468 (90333) | -0.277037763 | 52 | KCNH7 (90134) | 0.238259576 |
| 53 | OTOP1 (133060) | -0.276718028 | 53 | BANP (54971) | 0.237734635 |
| 54 | PSMB6 (5694) | -0.276588303 | 54 | RCL1 (10171) | 0.236245413 |
| 55 | IFITM3 (10410) | -0.275424933 | 55 | SCAP (22937) | 0.23545991 |
| 56 | ABCB10 (23456) | -0.274679509 | 56 | RRAGD (58528) | 0.235073286 |
| 57 | COX17 (10063) | -0.274487502 | 57 | MBTPS1 (8720) | 0.233867318 |
| 58 | JOSD1 (9929) | -0.272740347 | 58 | ZNF331 (55422) | 0.233529554 |
| 59 | MMS22L (253714) | -0.271941323 | 59 | PTPMT1 (114971) | 0.231764804 |
| 60 | DOK2 (9046) | -0.271177009 | 60 | POMP (51371) | 0.231594937 |
| 61 | MST1 (4485) | -0.270682783 | 61 | ZNF44 (51710) | 0.230429336 |
| 62 | ZNF695 (57116) | -0.270489865 | 62 | RPS10 (6204) | 0.230301586 |

| | | | | | |
|---|---|---|---|---|---|
| 63 | EPS8L3 (79574) | -0.269896999 | 63 | ARMC7 (79637) | 0.230192623 |
| 64 | PSPC1 (55269) | -0.269565103 | 64 | C3orf33 (285315) | 0.22857593 |
| 65 | PCTP (58488) | -0.269325376 | 65 | RFPL4AL1 (729974) | 0.227199716 |
| 66 | BCLAF1 (9774) | -0.267905187 | 66 | STK11 (6794) | 0.227108805 |
| 67 | NUDT4 (11163) | -0.266536226 | 67 | ENY2 (56943) | 0.226739738 |
| 68 | CYB561 (1534) | -0.266294257 | 68 | ZNF560 (147741) | 0.225968565 |
| 69 | VEZF1 (7716) | -0.266025953 | 69 | PSMC1 (5700) | 0.22553616 |
| 70 | C12orf71 (728858) | -0.265818025 | 70 | GMPPB (29925) | 0.225530658 |
| 71 | F7 (2155) | -0.265612676 | 71 | HYOU1 (10525) | 0.225503508 |
| 72 | TRIM61 (391712) | -0.265000953 | 72 | SMR3A (26952) | 0.224721165 |
| 73 | RMDN1 (51115) | -0.264960042 | 73 | ZMAT5 (55954) | 0.222454829 |
| 74 | DYRK1A (1859) | -0.264652377 | 74 | CCNK (8812) | 0.222454037 |
| 75 | ANKRD40 (91369) | -0.264582839 | 75 | DPH3 (285381) | 0.221826063 |
| 76 | RHOA (387) | -0.264434785 | 76 | MLLT1 (4298) | 0.221490476 |
| 77 | RUSC1 (23623) | -0.264426395 | 77 | PITRM1 (10531) | 0.221222885 |
| 78 | TBC1D3H (729877) | -0.264191124 | 78 | UBE2V1 (7335) | 0.220667806 |
| 79 | MRTO4 (51154) | -0.264122329 | 79 | CDC37 (11140) | 0.220523965 |
| 80 | METTL15 (196074) | -0.263926132 | 80 | LCE1F (353137) | 0.220173385 |
| 81 | ATG4D (84971) | -0.263585808 | 81 | TXNL4B (54957) | 0.219776337 |
| 82 | HIST1H1D (3007) | -0.263562877 | 82 | TELO2 (9894) | 0.219330863 |
| 83 | PPP4R2 (151987) | -0.263550554 | 83 | ERCC3 (2071) | 0.218896925 |
| 84 | CPT1B (1375) | -0.263507892 | 84 | ELP4 (26610) | 0.218434393 |
| 85 | DAXX (1616) | -0.262460824 | 85 | RARS (5917) | 0.218093863 |
| 86 | CCDC62 (84660) | -0.261461123 | 86 | AREG (374) | 0.218085707 |
| 87 | KRTAP10-9 (386676) | -0.260862765 | 87 | WDR70 (55100) | 0.217608608 |
| 88 | ADAMTS15 (170689) | -0.259265427 | 88 | PHB (5245) | 0.217424915 |
| 89 | NASP (4678) | -0.258956897 | 89 | PMAIP1 (5366) | 0.217419874 |
| 90 | CBWD5 (220869) | -0.258792908 | 90 | LILRB3 (11025) | 0.217385006 |
| 91 | SNAI2 (6591) | -0.258660921 | 91 | INTS2 (57508) | 0.216765431 |
| 92 | RHOF (54509) | -0.257884813 | 92 | TYMS (7298) | 0.21657485 |
| 93 | IPO5 (3843) | -0.257011905 | 93 | HSD17B12 (51144) | 0.216142832 |
| 94 | RNPS1 (10921) | -0.257007842 | 94 | OST4 (100128731) | 0.215833598 |
| 95 | LPA (4018) | -0.255521867 | 95 | CCDC93 (54520) | 0.215620383 |
| 96 | STARD3 (10948) | -0.255295573 | 96 | RBM18 (92400) | 0.214399514 |
| 97 | RPL26L1 (51121) | -0.254252071 | 97 | NELFCD (51497) | 0.214184898 |
| 98 | DND1 (373863) | -0.253953064 | 98 | STEAP1 (26872) | 0.214055986 |
| 99 | TOM1L1 (10040) | -0.253754599 | 99 | LENEP (55891) | 0.213040007 |
| 100 | SUGT1 (10910) | -0.253499557 | 100 | CBWD6 (644019) | 0.213032085 |
| 101 | BRIP1 (83990) | -0.253336105 | | | |
| 102 | HSPA1B (3304) | -0.253213186 | | | |
| 103 | RPP25L (138716) | -0.252687459 | | | |
| 104 | HIST1H3G (8355) | -0.252648445 | | | |
| 105 | DUSP22 (56940) | -0.251807834 | | | |
| 106 | KRTAP4-2 (85291) | -0.251713332 | | | |
| 107 | FBXO22 (26263) | -0.251350403 | | | |
| 108 | ME2 (4200) | -0.251336184 | | | |
| 109 | RNF4 (6047) | -0.250655156 | | | |
| 110 | COG3 (83548) | -0.25010706 | | | |
| 111 | SPRY2 (10253) | -0.249841963 | | | |

*Table S3, Medulloblastoma Rang. Related to Figure S4.*

| Rang | GENE | Δ TP53 CERES |
|---|---|---|
| 1 | **MDM2 (4193)** | **-0.700515819** |
| 2 | WRAP73 (49856) | -0.579775076 |
| 3 | YRDC (79693) | -0.533870687 |
| 4 | SUPT20H (55578) | -0.520397393 |
| 5 | KIF18B (146909) | -0.510145141 |
| 6 | **PPM1D (8493)** | **-0.509476432** |
| 7 | CDAN1 (146059) | -0.50853806 |
| 8 | GMPS (8833) | -0.493104014 |
| 9 | AGPAT3 (56894) | -0.450711844 |
| 10 | TYMS (7298) | -0.450064118 |
| 11 | TAF5L (27097) | -0.431352544 |
| 12 | UBE2D3 (7323) | -0.426474411 |
| 13 | DHFR (1719) | -0.418990151 |
| 14 | TADA2B (93624) | -0.414844598 |
| 15 | DTYMK (1841) | -0.399724411 |
| 16 | DNAJC9 (23234) | -0.397006324 |
| 17 | ATXN7L3 (56970) | -0.394424275 |
| 18 | PAICS (10606) | -0.391497903 |
| 19 | INTS6 (26512) | -0.388452454 |
| 20 | ADSL (158) | -0.37970745 |
| 21 | NOP9 (161424) | -0.374645377 |
| 22 | MEN1 (4221) | -0.372236169 |
| 23 | ERCC4 (2072) | -0.358299206 |
| 24 | PHC2 (1912) | -0.358188813 |
| 25 | SPICE1 (152185) | -0.355681707 |
| 26 | SARNP (84324) | -0.350049898 |
| 27 | EP300 (2033) | -0.347226009 |
| 28 | CEP135 (9662) | -0.346400167 |
| 29 | KAT2B (8850) | -0.344744161 |
| 30 | FOXC2 (2303) | -0.344217506 |
| 31 | AXIN1 (8312) | -0.343560514 |
| 32 | DCPS (28960) | -0.343174108 |
| 33 | NXT1 (29107) | -0.343127092 |
| 34 | HDAC2 (3066) | -0.341886354 |
| 35 | FURIN (5045) | -0.339925212 |
| 36 | POT1 (25913) | -0.338235072 |
| 37 | CCDC77 (84318) | -0.337219526 |
| 38 | OR10H1 (26539) | -0.336950848 |
| 39 | TRABD (80305) | -0.33152913 |
| 40 | TRPM7 (54822) | -0.329734466 |
| 41 | **USP7 (7874)** | -0.323401743 |
| 42 | CRKL (1399) | -0.3224495 |
| 43 | EED (8726) | -0.320532327 |
| 44 | CNTNAP3B (728577) | -0.319767698 |
| 45 | SERF2 (10169) | -0.319374678 |
| 46 | AHCYL1 (10768) | -0.316390048 |
| 47 | MVD (4597) | -0.316059439 |
| 48 | FKBP1A (2280) | -0.312227428 |
| 49 | CCDC101 (112869) | -0.311642505 |
| 50 | MINOS1 (440574) | -0.311524661 |
| 51 | ATP1A3 (478) | -0.310413834 |
| 52 | MSRB1 (51734) | -0.309678926 |
| 53 | ARHGEF7 (8874) | -0.307147811 |
| 54 | DOT1L (84444) | -0.306315993 |
| 55 | NDUFA3 (4696) | -0.303483232 |
| 56 | KRTAP10-9 (386676) | -0.30155291 |
| 57 | CYP2B6 (1555) | -0.299886327 |
| 58 | HIST1H2BI (8346) | -0.29977035 |
| 59 | COX20 (116228) | -0.298266054 |
| 60 | KIAA1731 (85459) | -0.297064078 |
| 61 | TKT (7086) | -0.295717022 |
| 62 | PPA2 (27068) | -0.295676483 |
| 63 | SASS6 (163786) | -0.295512331 |

| Inverse Rang | GENE | Δ TP53 CERES |
|---|---|---|
| 1 | **TP53 (7157)** | **0.619256776** |
| 2 | WDR73 (84942) | 0.538995138 |
| 3 | ITGAV (3685) | 0.527889311 |
| 4 | TSC1 (7248) | 0.503368787 |
| 5 | SNRPB2 (6629) | 0.492234608 |
| 6 | TSC2 (7249) | 0.474121507 |
| 7 | MBNL1 (4154) | 0.461316786 |
| 8 | ZFP36L1 (677) | 0.455887289 |
| 9 | RPRD2 (23248) | 0.396298624 |
| 10 | KCTD5 (54442) | 0.385470908 |
| 11 | RAB7A (7879) | 0.384388585 |
| 12 | TGFBR1 (7046) | 0.377145759 |
| 13 | ARMC5 (79798) | 0.3714635 |
| 14 | BRAT1 (221927) | 0.367064467 |
| 15 | FERMT2 (10979) | 0.367043933 |
| 16 | MSTO1 (55154) | 0.360489647 |
| 17 | SRP9 (6726) | 0.346218325 |
| 18 | KCTD10 (83892) | 0.32539193 |
| 19 | MGAT1 (4245) | 0.316316643 |
| 20 | CTR9 (9646) | 0.313935154 |
| 21 | UBC (7316) | 0.309367268 |
| 22 | MAP3K7 (6885) | 0.302309747 |
| 23 | TARDBP (23435) | 0.299858289 |
| 24 | WDR61 (80349) | 0.293164759 |
| 25 | CEPT1 (10390) | 0.289997081 |
| 26 | PIK3R1 (5295) | 0.286136098 |
| 27 | ARL15 (54622) | 0.28605914 |
| 28 | KPNA2 (3838) | 0.284705932 |
| 29 | COPG1 (22820) | 0.281467881 |
| 30 | FOXN3 (1112) | 0.279480162 |
| 31 | EIF2AK4 (440275) | 0.277630296 |
| 32 | EAF1 (85403) | 0.276295542 |
| 33 | PPM1A (5494) | 0.273007381 |
| 34 | BRF2 (55290) | 0.270531816 |
| 35 | GRB2 (2885) | 0.268299144 |
| 36 | ACTR3 (10096) | 0.266945802 |
| 37 | ITGB5 (3693) | 0.266334287 |
| 38 | AHCTF1 (25909) | 0.265503288 |
| 39 | NPRL2 (10641) | 0.263632486 |
| 40 | TXNDC9 (10190) | 0.262622761 |
| 41 | TIMM9 (26520) | 0.261253478 |
| 42 | ANKRD28 (23243) | 0.260709751 |
| 43 | KEAP1 (9817) | 0.260084603 |
| 44 | DEPDC5 (9681) | 0.258831036 |
| 45 | RTFDC1 (51507) | 0.25731604 |
| 46 | THOC1 (9984) | 0.252647784 |
| 47 | TCF12 (6938) | 0.249430987 |
| 48 | PTPN11 (5781) | 0.24810423 |
| 49 | PZP (5858) | 0.247074507 |
| 50 | INTS12 (57117) | 0.244342196 |
| 51 | LPAR1 (1902) | 0.242659458 |
| 52 | SRSF1 (6426) | 0.242629575 |
| 53 | NUP85 (79902) | 0.242278686 |
| 54 | AREG (374) | 0.240903792 |
| 55 | TRAPPC8 (22878) | 0.239312372 |
| 56 | ZMYND8 (23613) | 0.239258435 |
| 57 | RAG2 (5897) | 0.237023455 |
| 58 | PRUNE2 (158471) | 0.236196354 |
| 59 | TIMM23 (100287932) | 0.235542187 |
| 60 | ALG1 (56052) | 0.234987312 |
| 61 | LY6D (8581) | 0.231578197 |
| 62 | SKP2 (6502) | 0.231211019 |
| 63 | EIF2B2 (8892) | 0.230577057 |

| | | | | | |
|---|---|---|---|---|---|
| 64 | **MDM4 (4194)** | -0.2938428 | 64 | HSBP1 (3281) | 0.229953284 |
| 65 | TADA1 (117143) | -0.293467206 | 65 | DHX9 (1660) | 0.229392847 |
| 66 | CST2 (1470) | -0.291078491 | 66 | FBXW11 (23291) | 0.229196572 |
| 67 | CTPS1 (1503) | -0.291023646 | 67 | ECD (11319) | 0.228610025 |
| 68 | COX17 (10063) | -0.29055972 | 68 | SCG3 (29106) | 0.227986448 |
| 69 | C16orf89 (146556) | -0.289829563 | 69 | OR4C15 (81309) | 0.227373716 |
| 70 | ZNF486 (90649) | -0.289373213 | 70 | CTDNEP1 (23399) | 0.226901936 |
| 71 | ATIC (471) | -0.289317921 | 71 | E2F5 (1875) | 0.225090252 |
| 72 | REXO2 (25996) | -0.288874356 | 72 | CYFIP1 (23191) | 0.224739732 |
| 73 | NSMCE1 (197370) | -0.287692562 | 73 | ETF1 (2107) | 0.223931749 |
| 74 | DHFRL1 (200895) | -0.286829031 | 74 | THOC5 (8563) | 0.22351144 |
| 75 | CAD (790) | -0.286124966 | 75 | PABPN1 (8106) | 0.223052264 |
| 76 | SMARCA5 (8467) | -0.283787415 | 76 | LRP8 (7804) | 0.22165166 |
| 77 | FAM72B (653820) | -0.283473486 | 77 | TAX1BP3 (30851) | 0.221441984 |
| 78 | CCNF (899) | -0.283290549 | 78 | CPEB4 (80315) | 0.221436235 |
| 79 | MR1 (3140) | -0.2826881 | 79 | SLC38A2 (54407) | 0.219911052 |
| 80 | NPIPB6 (728741) | -0.282580773 | 80 | COBLL1 (22837) | 0.219412161 |
| 81 | SURF2 (6835) | -0.282287881 | 81 | VCL (7414) | 0.219208345 |
| 82 | FGFR1OP (11116) | -0.281340548 | 82 | CHMP3 (51652) | 0.218066531 |
| 83 | PARN (5073) | -0.28093143 | 83 | ERH (2079) | 0.217621702 |
| 84 | HEATR3 (55027) | -0.280340527 | 84 | SMCHD1 (23347) | 0.216435483 |
| 85 | ANKRD52 (283373) | -0.27946987 | 85 | FAM228A (653140) | 0.216262478 |
| 86 | OAZ1 (4946) | -0.279214514 | 86 | CCBE1 (147372) | 0.215505689 |
| 87 | IGF1R (3480) | -0.27910879 | 87 | FRS2 (10818) | 0.215500854 |
| 88 | LIN37 (55957) | -0.278690176 | 88 | LHX8 (431707) | 0.213860158 |
| 89 | CS (1431) | -0.278220978 | 89 | MON2 (23041) | 0.213730016 |
| 90 | SDHD (6392) | -0.275829463 | 90 | PLA1A (51365) | 0.213530126 |
| 91 | TJP1 (7082) | -0.274951676 | 91 | ZNF160 (90338) | 0.213159484 |
| 92 | CYP4F11 (57834) | -0.274755674 | 92 | VPS4B (9525) | 0.212709786 |
| 93 | DYNLT1 (6993) | -0.274450569 | 93 | TMEM165 (55858) | 0.211670913 |
| 94 | CTDSPL2 (51496) | -0.273089232 | 94 | JTB (10899) | 0.211504572 |
| 95 | KRTAP9-1 (728318) | -0.272273165 | 95 | CCNE2 (9134) | 0.211287231 |
| 96 | C16orf72 (29035) | -0.271629812 | 96 | PFN1 (5216) | 0.210882181 |
| 97 | CCND1 (595) | -0.271344827 | 97 | MIA2 (117153) | 0.210846536 |
| 98 | TYW1 (55253) | -0.271334229 | 98 | ZNF329 (79673) | 0.210563192 |
| 99 | CDK6 (1021) | -0.271109854 | 99 | SATB1 (6304) | 0.210191813 |
| 100 | TPPP (11076) | -0.270401359 | 100 | CHMP7 (91782) | 0.209615714 |

*Table S4. List of tumors developed in irradiated PPM1D transgenic mice, Related to Figure 4.*

| Diagnosis | n | Age at irr[b] (days) | Time from irr to tumor development[b] (days) | Metastatic spread[c] |
|---|---|---|---|---|
| ***PPM1D* -positive mice** | | | | |
| **Thymic lymphoblastic lymphoma** | **41** | Mdn: 6(1-314) | Mdn: 196 (124-535) | Systemic |
| **Leukemia/lymphoma** | **9** | Mdn: 48(4-314) | Mdn: 342 (195-482) | Systemic |
| **Other solid tumors:** | **25** | Mdn: 20(3-240) | Mdn: 435(246-683) | |
| Papillary serous ovarian cancer | 5 | Mdn: 20(4-164) | Mdn: 415(389-598) | Splenomegaly |
| Lacrimal gland tumor | 6 | Mdn: 455(291-614) | Mdn: 20 (4-94) | Splenomegaly |
| Gastrointestinal stromal tumor | 1 | 62 | 463 | Splenomegaly |
| Gastric adenocarcinoma | 1 | 62 | 463 | Lung, splenomegaly |
| Adenocarcinoma of the lung | 4 | Mdn: 414(246-463) | Mdn: 34(4-240) | |
| Angiosarcoma: | 2 | | | |
| #1 | | 48 | 199 | |
| #2 | | 14 | 319 | Splenomegaly |
| Osteosarcoma metastasis in the lung[a] | 1 | 10 | 303 | Lung, Splenomegaly |
| Neuroblastoma/Adrenal tumor: | 2 | | | |
| #1 | | 6 | 683 | Liver, splenomegaly |
| #2 | | 6 | 522 | |
| **Total nr of tumors** | **75** | | | |
| **Wild-type mice** | | | | |
| **Thymic lymphoblastic lymphoma** | **4** | Mdn: 16(14-87) | Mdn: 334 (220-596) | Systemic |
| **Adenocarcinoma of the lung** | **1** | 14 | 437 | |
| **Lacrimal gland tumor** | **1** | 17 | 517 | |
| **Total nr of tumors** | **6** | | | |

[a] Presence of proliferative lymphoblastic cells in the spleen indicative of leukemia or lymphoma.

[b] Age in days given as median (Mdn) and range for groups.

[c] And/or other tumor manifestations such as splenomegaly.

Systemic spread implicate infiltrates of atypical lymphocytes/lymphoblasts engaging the spleen, liver and in some cases the kidneys.

In total, 75 tumors were diagnosed in 72 PPM1D-positive mice (three mice were diagnosed having two primary tumors each).

**Available upon request**

*Table S5. Summary of relevant genomic alteration detected through WES, Related to Figure 5.*

*Table S5. Non-synonymous variants in sequenced samples, Related to Figure 5.*

*Table S5. Exome and RNA Sequencing information,  Related to Figure 5.*

*Table S5. Notch & Pten aberrations, Related to Figure 5.*

*Table S5. Differential gene expression analysis, 4378 downregulated genes, Related to Figure 5.*

*Table S5. Differential gene expression analysis, 4138 upregulated genes, Related to Figure 5.*

*Table S6. IC50 µM with 95% confidence interval for SL-176, SP-001 and CCT007093, Related to Fig 6A.*

| Origin | Cell line | SL-176 | SP-001 | CCT007093 |
|---|---|---|---|---|
| Neuroblastoma | NB1691 | 1.3 (1.0-1.7) | 11 (6.2-21) | 43 (35-53) |
| | SH-SY5Y | 0.73 (0.65-0.83) | 10 (8.2-13) | 27 (21-34) |
| | SK-N-AS | 0.65 (0.61-0.69) | 5.4 (4.5-6.4) | 49 (21-110) |
| | SK-N-BE(2) | 0.57 (0.44-0.72) | 5.4 (4.4-6.6) | 48 (38-170) |
| | TR14 | 1.0 (0.93-1.1) | 13 (7.7-21) | 33 (25-43) |
| Breast cancer | MCF-7 | 1.2 (0.91-1.5) | 41 (20-85) | 47 (31-53) |

$IC_{50}$ values were determined from cell viability data from the MTT method.

The cells were incubated for 72 h, tested in duplicates and the experiments were repeated at least three time[illegible]

*Table S6. IC50 µM with 95% confidence interval for SL-176, RITA and Nutlin-3, Related to Fig 6B and Supp. Fig S6B,C.*

| Origin | Cell line | SL-176 | RITA | Nutlin-3 |
|---|---|---|---|---|
| Neuroblastoma | IMR-32 | 0.63 (0.53-0.75) | 0.26 (0.21-0.32) | 0.47 (0.39-0.55) |
| | Kelly | 0.87 (0.58-1.3) | 0.17 (0.12-0.24) | 16 (5.8-44) |
| | NB1691 | 1.3 (1.0-1.7) | 10 (6.8-15) | 1.1 (0.95-1.3) |
| | SH-EP | 1.2 (1.0-1.5) | >50 [a] | 4.7 (4.1-5.4) |
| | SH-SY5Y | 0.73 (0.65-0.83) | 0.24 (0.12-0.45) | 1.0 (0.89-1.2) |
| | SK-N-AS | 0.65 (0.61-0.69) | 2.6 (0.93-7.5) | 11 (9.2-13) |
| | SK-N-BE(2) | 0.57 (0.44-0.72) | 2.1 (1.5-3.1) | 14 (12-17) |
| | SK-N-DZ | 1.0 (0.80-1.3) | 2.4 (1.7-3.4) | 14 (12-15) |
| | SK-N-FI | 0.44 (0.36-0.52) | 5.1 (3.4-7.9) | 12 (10-14) |
| | SK-N-SH | 0.54 (0.45-0.64) | 0.84 (0.67-1.1) | 0.88 (0.56-1.4) |
| | TR14 | 1.0 (0.93-1.1) | 8.1 (4.7-14) | 1.8 (1.4-2.4) |
| Medulloblastoma | DAOY | 2.6 (2.3-3.0) | 0.87 (0.72-1.1) | 5.0 (4.4-5.5) |
| | D458 MED | 1.8 (1.3-2.4) | 0.25 (0.19-0.33) | 0.59 (0.46-0.77) |
| | MED 8A | 1.2 (0.93-1.5) | 0.098 (0.0035-0.27) | 7.1 (5.6-9.0) |
| | UW228-3 | 0.71 (0.49-1.0) | 23 (16-32) | 15 (12-18) |
| sPNET | PFSK-1 | 0.92 (0.78-1.1) | 3.2 (2.3-4.5) | 7.9 (5.9-11) |
| Breast cancer | MCF-7 | 1.2 (0.91-1.5) | 1.3 (0.64-2.5) | 6.2 (5.3-7.2) |
| Fibroblast | MRC-5 | 2.8 (1.8-4.3) | >50* | 5.3 (2.1-13) |
| Mouse neural stem cell | C17.2 | 22 (10-49) | 9.5 (8.6-11) | 2.0 (1.7-2.4) |

$IC_{50}$ values were determined from cell viability data from the WST-1 method.

The cells were incubated for 72 h, tested in duplicates and the experiments were repeated at least three times.

[a]The highest tested concentration was 50 µM and it did not reach $IC_{50}$.

*Table S7. List of antibodies used in cell line and mouse tissue analysis by western blot, Related to STAR methods.*

| Antibody | Manufacturer | Cat No | Blocking solution | Antibody dilution | Antibody specie |
|---|---|---|---|---|---|
| **phospho ATM (S1981)** | Cell Signaling | 45265 | 5% BSA in TBS-T | 1:500 | MOUSE |
| **ATM** | Cell Signaling | 2873 | 5% BSA in TBS-T | 0.736111111 | RABBIT |
| **PARP** | Cell Signaling | 9542 | 5% milk in TBS-T | 0.736111111 | RABBIT |
| **phospho p53** | Cell Signaling | 9284 | 5% milk in TBS-T | 0.736111111 | RABBIT |
| **p53** | Cell Signaling | 9282 | 5% milk in TBS-T | 0.736111111 | RABBIT |
| **p53 (D2H9O)** | Cell Signaling | 32532 | 5% milk in TBS-T | 0.736111111 | RABBIT |
| **phospho p38 (D3F9)** | Cell Signaling | 4511 | 5% BSA in TBS-T | 1.430555556 | RABBIT |
| **p38** | Cell Signaling | 9212 | 5% BSA in TBS-T | 1.430555556 | RABBIT |
| **WIP1** | Cell Signaling | 11901 | 5% BSA in TBS-T | 0.736111111 | RABBIT |
| **WIP1 (H300)** | Santa Cruz Biotechnology | SC20712 | 5% milk in TBS-T | 0.180555556 | RABBIT |
| **phospho CHK1** | Cell Signaling | 12302 | 5% BSA in TBS-T | 1.430555556 | RABBIT |
| **CHK1** | Cell Signaling | 2360 | 5% milk in TBS-T | 1.430555556 | MOUSE |
| **phospho CHK2 (T68)** | Cell Signaling | 2661 | 5% BSA in TBS-T | 0.736111111 | RABBIT |
| **CHK2** | Cell Signaling | 2662 | 5% BSA in TBS-T | 1:500 | RABBIT |
| **GAPDH** | Abcam | ab8245 | 5% milk in TBS-T | 1:10000 | MOUSE |
| **β-tubulin** | Sigma-Aldrich | T7816 | 5% milk in TBS-T | 1:2500 | MOUSE |
| **VINCULIN** | Abcam | 129002 | 5% milk in TBS-T | 2.819444444 | RABBIT |

*Table S7. List of antibodies used in transgenic mouse tissue analysis by immunohistochemistry, Related to STAR methods.*

| Antibody | Manufacturer | Cat No | Antibody dilution | Antibody specie |
|---|---|---|---|---|
| **Aktin** | Thermo Fischer Scientific | PA5-19465 | 1:500 | RABBIT |
| **CD117/C-kit** | Agilent | A450229-2 | 0.25 | RABBIT |
| **CD20** | Cell Marque | 120R-14 | 0.180555556 | RABBIT |
| **CD3** | Agilent | IS50330-2 | RTU | RABBIT |
| **CD34** | Abcam | ab81289 | 0.180555556 | RABBIT |
| **CD56** | Agilent | M7304 | 0.180555556 | MOUSE |
| **CD79** | Cell Marque | 179R-14 | 0.25 | RABBIT |
| **Chromogranin A** | Abcam | ab15160 | 0.319444444 | RABBIT |
| **Cytokeratin 20** | Abcam | ab181598 | 1.430555556 | RABBIT |
| **Dog1** | Thermo Fischer Scientific | RM-9132 | 0.111111111 | RABBIT |
| **ER** | Agilent | IR151 | RTU | RABBIT |
| **ERG** | Agilent | IR65961-2 | RTU | RABBIT |
| **Faktor VIII** | Agilent | IR52761-2 | RTU | RABBIT |
| **HNF1B** | Thermo Fischer Scientific | PA5-50531 | 1:200 | RABBIT |
| **LCA/CD45** | Abcam | ab10558 | 0.388888889 | RABBIT |
| **Napsin A** | Biocare Medicals | ACI3043A | 1:200 | RABBIT |
| **NF** | Agilent | IR60761-2 | RTU | MOUSE |
| **NF** | Abcam | ab204893 | 0.180555556 | RABBIT |
| **PHOXB2** | Abcam | ab183741 | 0.736111111 | RABBIT |
| **PR** | Ventana | 790-2223 | RTU | RABBIT |
| **S100** | Agilent | IR50461-2 | 0.180555556 | RABBIT |
| **SATB2** | Abcam | ab92446 | 0.180555556 | RABBIT |
| **Synaptofysin** | Abcam | Ab16659 | 1:40 | RABBIT |
| **TDT** | Agilent | IR09361-2 | RTU | RABBIT |
| **TTF1** | Abcam | ab76013 | 0.215277778 | RABBIT |
| **Vimentin** | Abcam | ab92547 | 0.25 | RABBIT |
| **WT1** | Abcam | ab89901 | 0.25 | RABBIT |
